# Ancient DNA indicates 3,000 years of genetic continuity in the Northern Iranian Plateau, from the Copper Age to the Sassanid Empire

**DOI:** 10.1101/2025.02.03.636298

**Authors:** Motahareh Ala Amjadi, Yusuf Can Özdemir, Maryam Ramezani, Kristóf Jakab, Melinda Megyes, Arezoo Bibak, Zeinab Salehi, Zahra Hayatmehar, Mohammad Hossein Taheri, Hossein Moradi, Peyman Zargari, Ata Hasanpour, Vali Jahani, Abdol Motalleb Sharifi, Balázs Egyed, Balázs Gusztáv Mende, Mahmood Tavallaie, Anna Szécsényi-Nagy

**Author notes:** Corresponding authors: MT and ASN.

## Abstract

In this study, we present new ancient DNA data from prehistoric and historic populations of the Iranian Plateau. By analysing 50 samples from nine archaeological sites across Iran, we report 23 newly sequenced mitogenomes and 13 nuclear genomes, spanning 4700 BCE to 1300 CE. We integrate an extensive reference sample set of previously published ancient DNA datasets from Western and South-Central Asia, enhancing our understanding of genetic continuity and diversity within ancient Iranian populations. A new Early Chalcolithic sample, predating all other Chalcolithic genomes from Iran, demonstrates mostly Early Neolithic Iranian genetic ancestry. This finding reflects long-term cultural and biological continuity in and around the Zagros area, alongside evidence of some western genetic influence. Our sample selection prioritizes northern Iran, with a particular focus on the Achaemenid, Parthian, and Sassanid periods (355 BCE–460 CE). The genetic profiles of historical samples from this region position them as intermediates on an east-west genetic cline across the Persian Plateau. They also exhibit strong connections to local and South-Central Asian Bronze Age populations, underscoring enduring genetic connections across these regions. Diachronic analyses of uniparental lineages on the Iranian Plateau further highlight population stability from prehistoric to modern times.

## Introduction

The Persian Plateau in West Asia—defined here as a vast geographical area stretching from eastern Anatolia to Afghanistan and western parts of Pakistan, also including the south Caucasus, South-Central Asia, and north Mesopotamia^1–3^—played a significant role in shaping human prehistory, as a major hub for early *Homo sapiens* migration out of Africa^1,4^. Human populations colonized Eurasia, Oceania, and the Americas in multiple waves from this hub between 70/60 to 46 thousand years BP^1^. It is also recognized as a key region where the early diverging Basal Eurasian genetic lineage likely existed, described as an Upper Palaeolithic *Homo sapiens* genetic ancestry without admixture from *Homo neanderthalensis,* in contrast to other ancient non-Africans^5^. Prehistoric farmers, and later, ancient civilizations thrived on the Plateau, benefiting from its favorable geographic location and abundant resources, which fostered trade and intercultural relations^6^. Recent archaeogenetic studies have started to shed light on the complex nature of these ancient populations who inhabited the Persian Plateau^5,7–17^. Mesolithic hunter-gatherer (HG) remains from the Alborz Mountains in northern Iran show ancestries primarily related to the Basal Eurasian^5,12^. The Early Neolithic (EN) farmers from Ganj Dareh are proposed to be descendants of these Iranian HGs^16^ and are genetically cladal with the Caucasus HGs (CHG)^15^. These groups descend from the inferred Upper Palaeolithic period populations of Western Asia are a focal point for subsequent comparative genetic studies^4,7,8,15,16^. They carry distinct genetic components compared to the Neolithic farmers of the Levant and Anatolia, as well as the European HGs^5^. Further investigations revealed that the Chalcolithic farmers in both the western and eastern Iranian Plateau^5,16^ (defined here as the current territory of Iran) have additional ancestries related to the Natufian and Pre-Pottery Neolithic (PPN) Levant besides the Anatolian Neolithic Farmers (ANF)^13^. As the Chalcolithic period concluded and the Bronze Age (BA) began, more varied genetic components entered the ancient Iranian gene pool from the east and north, including the Andamanese hunter-gatherer-related (AHG), Ancient North Eurasian-related (ANE), and BA Steppe-related ancestries^5,12,14,16^. AHG-related ancestry is first detected in the region as an influx from South Asia (Indus Valley) at the BA Shahr-i Sokhta in eastern Iran^12^, a major population center around 3550 – 2900 BCE^18^ and a civilization center up to 2300 BCE (Supplementary Information). ANE-related ancestry is recognized in West Siberian Hunter-Gatherers (WSHG) and the Central-Asian Eneolithic/Chalcolithic Botai culture’s people, with genetic continuity from more archaic ANE populations of Northern Asia^19^. BA Steppe-related ancestry, which has genetic sharing with the CHG, and also with the ANE-related groups through its East European HG components, is often associated with the Yamnaya culture’s people and descendant groups, who spread across the Eurasian continent from the Pontic Steppe^12^. The diversity and integration of various genetic components in the Iranian Plateau became even more pronounced during the Iron Age (IA), evident in continuous Levant-related gene flow at this period^13^.

Despite this progress, our understanding of past genetic events remains incomplete as many populations are still uncharacterized, and the extensive human mobility in historical periods further complicates the genetic landscape^20^. This highlights the importance of studying the prehistoric and historical genetic makeup of the region to better understand cultural and population developments in later periods. This is particularly relevant for the northeastern and western regions of the Iranian Plateau, two areas with extensive connections to South Central Asia (including Turkmenistan, Uzbekistan, and Tajikistan) and South Caucasus (Azerbaijan, Armenia, and Georgia), respectively, from the Neolithic period to the historical times^6,21^ (see Supplementary information for more details).

These connections further expanded under the first Persian Empire, the Achaemenids, whose core territories included the Iranian Plateau, South Caucasus, South Central Asia, and regions of modern Afghanistan and Pakistan. In 323 BCE, the Seleucid Empire emerged as a Hellenistic state encompassing most of the Iranian Plateau^22^. By the mid-3rd century BCE, the warriors of the Aparni or Parni—a nomadic or semi-nomadic eastern Iranian tribe—rose to power and established the Parthian government in Nisa, from where they extended their control over the Persian Plateau. Accounts of the Parthian Empire and its tribes are documented in the works of Herodotus and Strabo^23,24^.

The catacomb type graves, also typical for the Parthians, were common on the entire territory of the Persian Plateau until the end of the BA. The oldest catacomb graves are dated to 3000–2900 BCE, excavated in the Jiroft area, predate those at Shahr-i Sokhta (after 2700 BCE)^25^ and South-Central Asia (e.g., Turkmenistan). This suggests that initially southeastern Iran influenced the burial practices of Baluchistan and southern Turkmenistan^26^. This tradition re-emerged during the Parthian and Sassanid periods in the north and northeastern Iranian Plateau (i.e., Liarsangbon and Vestemin, see Supplementary Information, Chapters 8 and 9, for more details).

The objective of our research is to conduct a genetic analysis of key archaeological sites in Iran, from the Chalcolithic period to the Medieval era. We focus particularly on the Achaemenid, Seleucid, Parthian, and Middle Sassanid periods (355 BCE-460 CE) in northern Iran, given its strategic location as a migration and trade corridor for millennia. We investigate three historical-period sites, and provide the first genetic evaluation of the northern Iranian Plateau’s “highway”, a key section of the Silk Roads^27^ that facilitated the distribution of diverse materials and artefacts south of the Caspian Sea (Figure 1).

**Figure 1.**
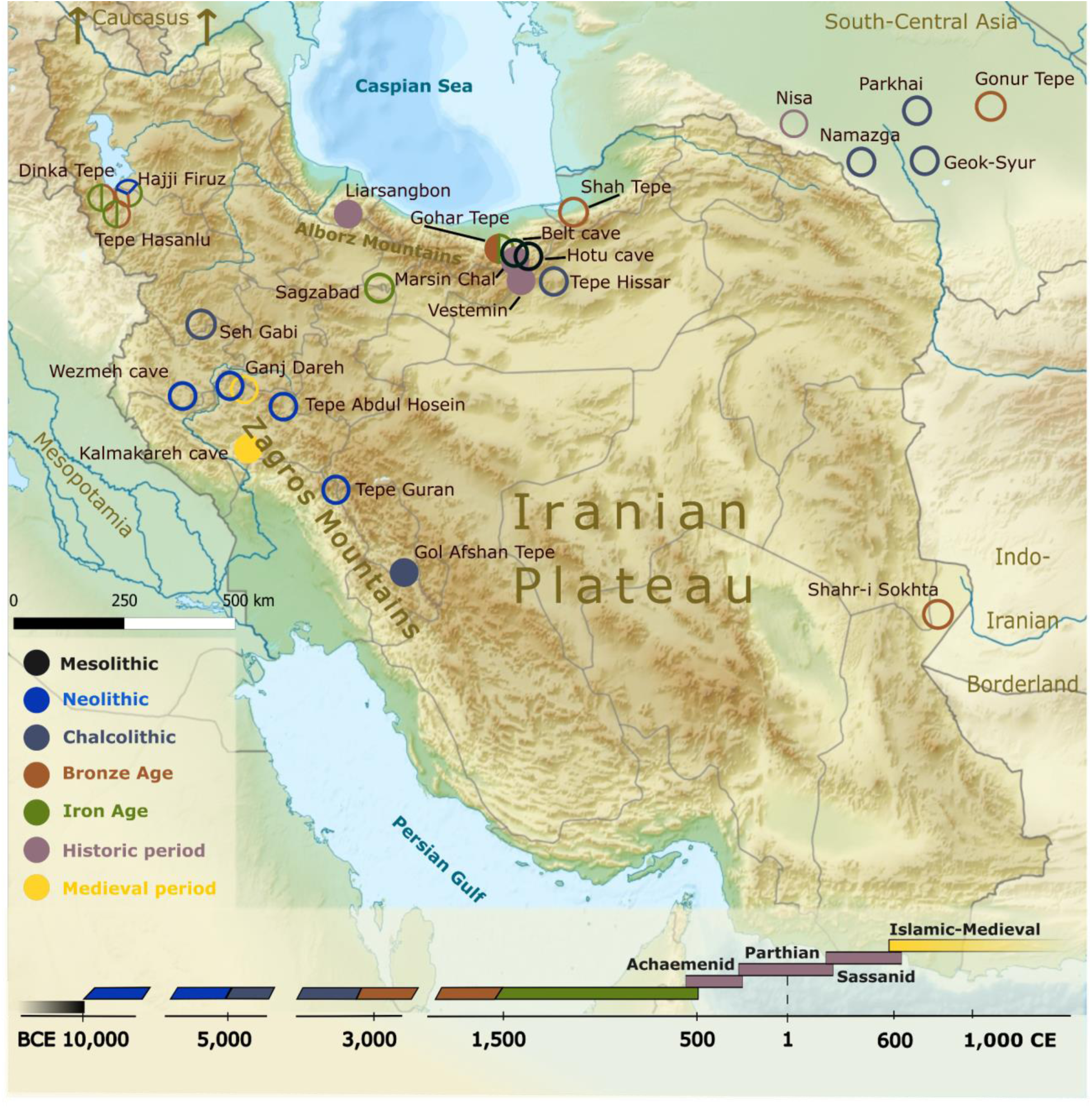
Map of the Iranian Plateau and its surroundings. Present-day Iran is highlighted, with the archaeological sites mentioned in the text. Filled circles represent new samples, empty circles mark sites from which ancient DNA data has been published in previous studies.^5,7–11,13,14,15,17^ For further sites that were processed but not analysed, refer to Supplementary Table S1.

Using genomic data, we aim to uncover genetic variation, as well as possible continuity and change among ancient Iranian populations and their connections to neighbouring populations (e.g., in the South-Caucasus and South-Central Asia), by comparing genetic and archaeological records across different regions. Our analyses seek to explore whether and how human movement, historical events, and cross-cultural interactions, such as trade, have influenced the genetic makeup of the Plateau’s populations over time.

## Results and Discussion

Based on initial screening and shallow shotgun sequencing of 50 human DNA samples, five samples met the criteria for direct deeper shotgun sequencing (Methods), which resulted in genomes with ∼0.20-1.11× average genomic coverage (mean ∼0.61×). The number of detected SNPs from the 1240k panel^28^ varies between 189k and 741k (mean ∼462k).

Initially, we used MyBaits Arbor Complete panel for genomic and mitogenome capture for the other samples. However, following a critical review of this method^29^, we repeated the target enrichment with Twist capture in seven cases^30^ to validate the results. Overall, eight samples were processed with MyBaits Arbor capture, with an average of ∼164k SNPs detected from the 1240k SNP set. For the Twist-captured samples, an average of 182k SNPs was covered, ranging between 11k and 770k SNPs. Finally, we sequenced 14 mitochondrial genomes after a separate mitogenome capture for samples from the Gohar Tepe, Marsin Chal, Liarsangbon, Vestemin, and Kalmakareh archaeological sites (see Methods, Table 1, Supplementary Tables S1-S4). The mean mtDNA coverage was 44.57×.

**Table 1.**
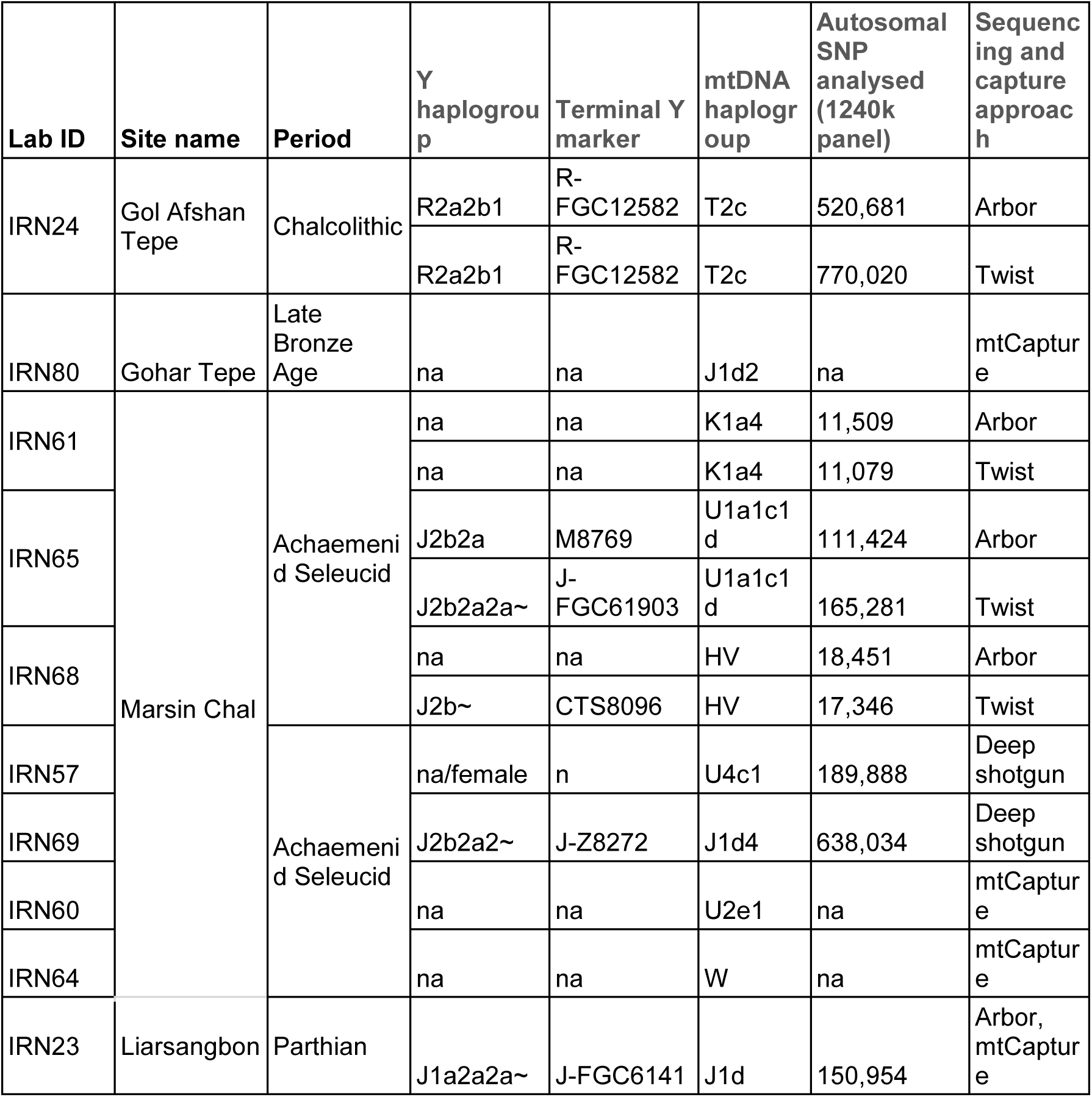
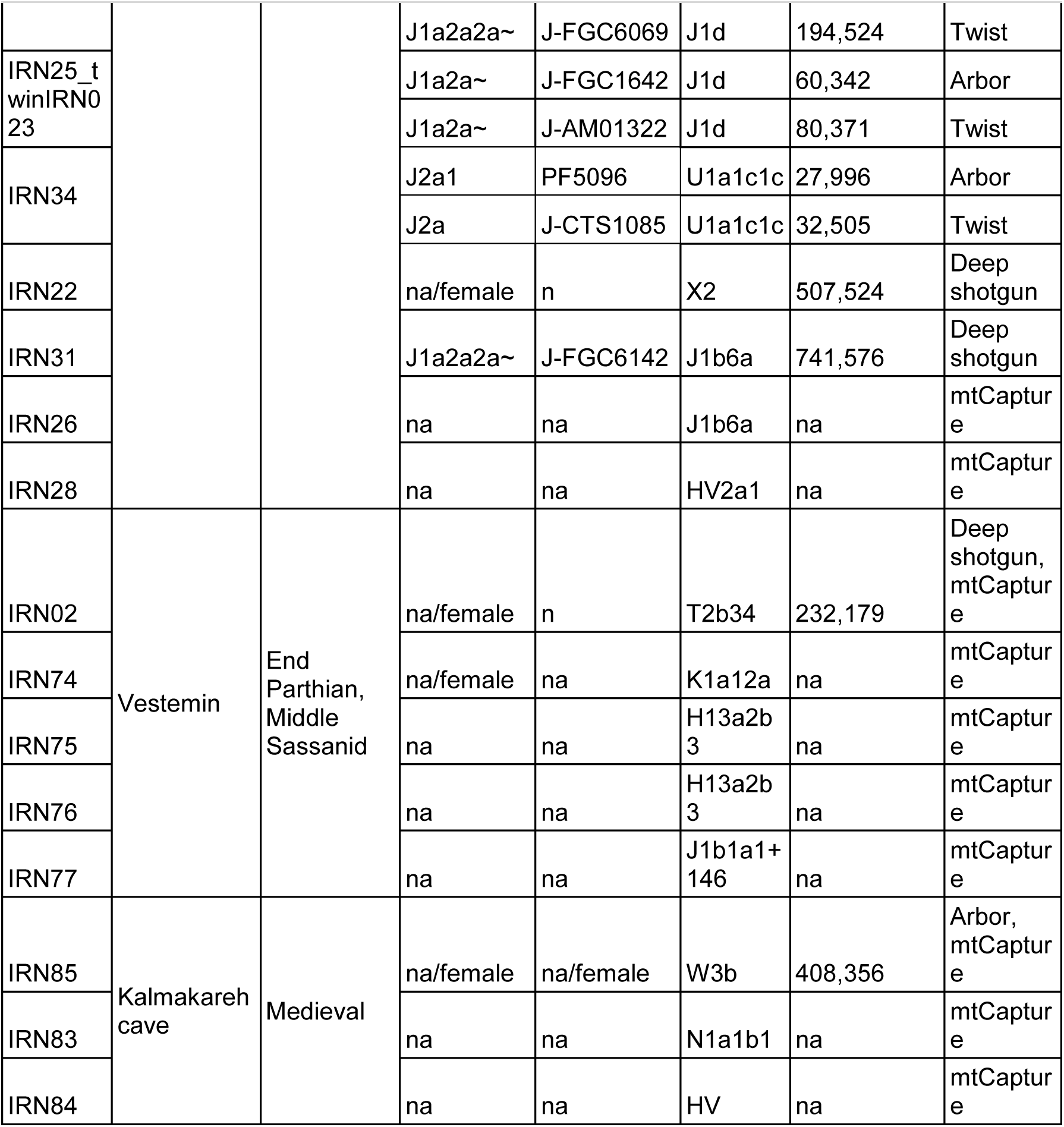
Archaeological and genetic information of the ancient samples sequenced for this study. The abbreviation ‘mt’ stands for mitochondrial in this table. Terminal Y markers were defined based on ISOGG v.15.73^31^. Details are seen in Supplementary Table S1-S4.

We used the BREADR package (Biological Relatedness from Ancient DNA in R) to assess kinship among the whole-genome data samples and found no close relatedness with sufficient SNP overlap between the compared individuals^32^. The only exception is a pair of genetically identical/twin individuals (IRN23 and IRN25) from Liarsangbon (Supplementary Table S8). Anthropological records confirm that these two individuals are twins (Supplementary Information, Chapter 8).

Analysing possible parental relatedness in the new genome-wide dataset from Iran, we detected varying runs of homozygosity (ROH)^33^ signals among individuals from different archaeological sites (Supplementary Table S15, Figure S1). Individual IRN22 from Liarsangbon stands out with the highest ROH, characterized by long homozygous runs (>20cM) indicating a high degree of inbreeding, likely due to parents who were first- or second-degree relatives^33^. In contrast, individuals IRN24 from Gol Afshan Tepe, IRN57 from Mersin Chal, IRN23 from Liarsangbon and IRN02 from Vestemin show minimal ROH, whereas others have nearly no signal, suggesting a more outbred ancestry and a relatively large mating pool (Supplementary Figure S1). Comparing the current dataset with previous analyses of this kind, we infer that signals of consanguinity are sporadically present across the broader Early Neolithic Iranian Plateau (Ganj Dareh, Tepe Abdul Hosein, Wezmeh) and in the Turkmenistan areas, which had cultural contact with Iranian Plateau, including Chalcolithic sites such as Namazga and Parkhai^21,34^ (Supplementary Table S15). Evidence of small effective population sizes are also apparent at the Chalcolithic to Iron Age sites Tepe Hissar, Shah Tepe, Dinkha Tepe, and Hasanlu of the Iranian Plateau, as well as BA Turkmenistan. Considering the available Iranian and Turkmenistan prehistoric datasets, we conclude that the effective population size increased over time from the Neolithic to the Iron Age, where the Neolithic mating pool was 3-6 times smaller than in later periods (Supplementary Table S15).

### A diachronic overview and frequency-based comparisons of the maternal lineages in the Iranian Plateau

To our knowledge, 127 ancient mitogenomes have been previously published from 20 archaeological sites on the Iranian Plateau, spanning the Mesolithic to Medieval times (Supplementary Table S5). We merged our new mtDNA results obtained from three different laboratory approaches (n=23), with these mtDNA data. The combined dataset allowed us to identify 23 distinct maternal macro-haplogroups in the region (Figure 2A).

**Figure 2.**
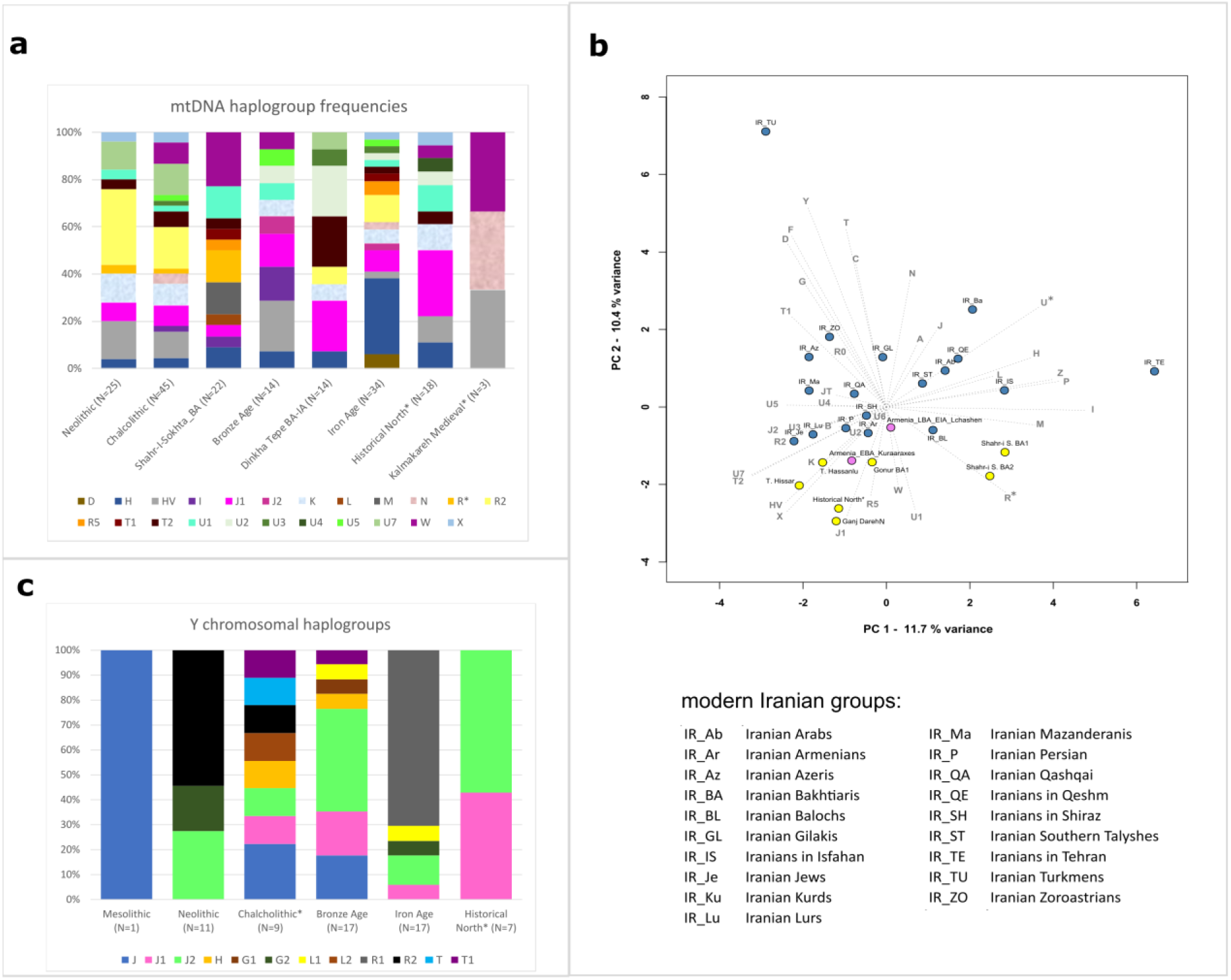
Summary of the uniparental data. (**A)** Distribution of the ancient Iranian mtDNA haplogroups; asterisks denote the samples analysed in this study. (**B)** PCA plot based on mtDNA haplogroup frequencies in modern Iranian and selected ancient Iranian groups. Using the frequencies of 36 haplogroups in modern and ancient Iranian populations, the first two principal components (PCs) capture 22.1% of the total variance (see Supplementary Table S6 for details). To maintain consistency and prevent sampling bias, we selected only those ancient groups that had more than 10 individuals or could be merged geographically and temporally, such as our historical samples. (**C)** Distribution of the ancient Iranian Y-chromosomal haplogroups (details in Supplementary Table S7).

A significant proportion of these ancient haplogroups correspond to the more ubiquitous West Eurasian lineages (70% in total), such as U, R, J, H, HV, and T, which are common even in the modern Iranian maternal gene pool^35^. In general, we detect a predominance of these maternal lineages on the Iranian Plateau during both prehistorical and historical periods. While the Neolithic-Chalcolithic periods show continuity to the BA on the maternal level, the latter period exhibits increasing variability with the appearance of several new mtDNA haplogroups. This trend is evident in the Shahr-i Sokhta dataset, which includes haplogroups such as L and M, R5 or T1^12^, as well as the arrival of additional Eastern Eurasian lineages (A, B, C, D, F, R, U6, Y and Z) in the post-BA times. Our findings align with earlier archaeological descriptions of increased population mobility^6,9^. The Northwestern Iranian IA sample set^7–10^ is dominated by Western Eurasian haplogroups, however the IA also marks the introduction of Eastern Asian haplogroup D to the Iranian Plateau (Sagzabad site in the northern Qazvin province)^11,17^. These patterns are consistent with the mtDNA gene pool of southern Turkmenistan during the Chalcolithic - Bronze Age (except haplogroup L), which was connected geographically and culturally to the eastern Iranian Plateau. The historical sample set shows the abundance of J1, along with an absence of R and Eastern Eurasian mitochondrial haplogroups.

We conducted principal component analysis (PCA) on the maternal haplogroup frequencies derived from a dataset of the first hypervariable region of the mtDNA (Supplementary Table S5-6, Figure 2B). Our data comprises samples from 19 modern Iranian groups (n=1,498)^35^, and some representative ancient populations from different periods of Iran, such as EN Ganj Dareh, Chalcolithic Tepe Hissar, BA Shahr-i Sokhta, IA Hasanlu, and the newly-analysed historical samples from the Marsin Chal, Liarsangbon, and Vestemin sites in northern Iran.

Our analyses reveal a distinct demarcation along PC1 between BA Shahr-i Sokhta samples, and other Iranian and Armenian ancient populations (Figure 2B). The new historical samples cluster together with EN Ganj Dareh, whereas Chalcolithic Tepe Hissar and IA Hasanlu are also positioned closely on the mtDNA PCA. Notably, some modern Iranian groups align with our historical samples along PC2, particularly the Iranian Armenians and Lurs, who share similar frequencies in haplogroups H and HV, and elevated proportions of K and J1 (Supplementary Figure S10.1). The new Medieval sample set from Kalmakareh demonstrates the local continuity of N and HV lineages in Lurestan Province through to the modern era^35^, where Iranian Bakhtiari nomadic people (with 51% N and 15% HV frequencies) have been recorded to inhabit for the last 800 years^36^. Trends in the mitochondrial gene pool are described in detail in Supplementary Information, Chapter 10 (Supplementary Table S5-6).

### Y-chromosomal haplogroup distribution

To achieve an overview of the paternal genomic composition of the region, we combined the new Y-chromosomal results (n=5) with data from previously published Mesolithic to historical-period males (n=54), originating from 16 different archaeological sites in Iran (Supplementary Table S7 and Figure 2C). The Y-chromosomal variation among the ancient Iranian populations mainly consisted of haplogroups J, G, L, R and T. The presence of the South Asian macro-haplogroup H, detected in Late Chalcolithic (Tepe Hissar) and Bronze Age (Shahr-i Sokhta) sites, aligns with the previously described cross-regional interactions characteristic of the Indo-Iranian borderland area^12^ (see more detail in Supplementary Information, Chapter 10). Haplogroups J and R2 were present across the Persian Plateau and its surroundings as early as the Mesolithic and Neolithic periods^5,7^. Our findings align with the previous research, indicating that these lineages have been present in the region for millennia. Haplogroup J1 is widely recognized to have originated approximately 20,000 years ago in a region encompassing western parts of the Persian Plateau^13,37^. In Iran, J1 can be traced back to the Chalcolithic period at the Seh Gabi site in the northwest^5^, and was also detected at the BA northwestern sites of Tepe Hasanlu and Dinkha Tepe (Supplementary Table S7). It persisted into the IA at Dinkha Tepe^13^, and evidence of its presence in the Plateau now extends into historical times with the newly analysed samples. Haplogroup J2 is considered to have formed approximately 31,600 years ago in a region covering northwestern and western Persian Plateau^38^. The frequency of J2 seems to have increased from the Neolithic era to historical periods in the Iranian Plateau, potentially due to greater interactions with regions to the west of the Iranian Plateu^13^ (Supplementary Table S7, Figure 2). More details on the newly analysed dataset can be found in Supplementary Information, Chapter 10.

### Population genomic trends from the Neolithic to Historic periods in the Iranian Plateau

In order to compare the new Iranian sample set to the published ancient and modern Eurasian populations, we applied genomic PCA on ancient and modern Eurasian and North African samples from the AADR Human Origins (HO) dataset v54^39^, supplemented with other relevant published datasets (Supplementary Table S9). *f*_4_-statistics and qpAdm^40,41^ were performed on subsets of the AADR 1240k and HO dataset and other published ancient individuals (see Methods). We performed supervised ADMIXTURE^42^ on a subset of this merged dataset (n=3913), as described in Methods, with results presented in Figure 3, Supplementary Figure S11.3 and Supplementary Table S10.

**Figure 3.**
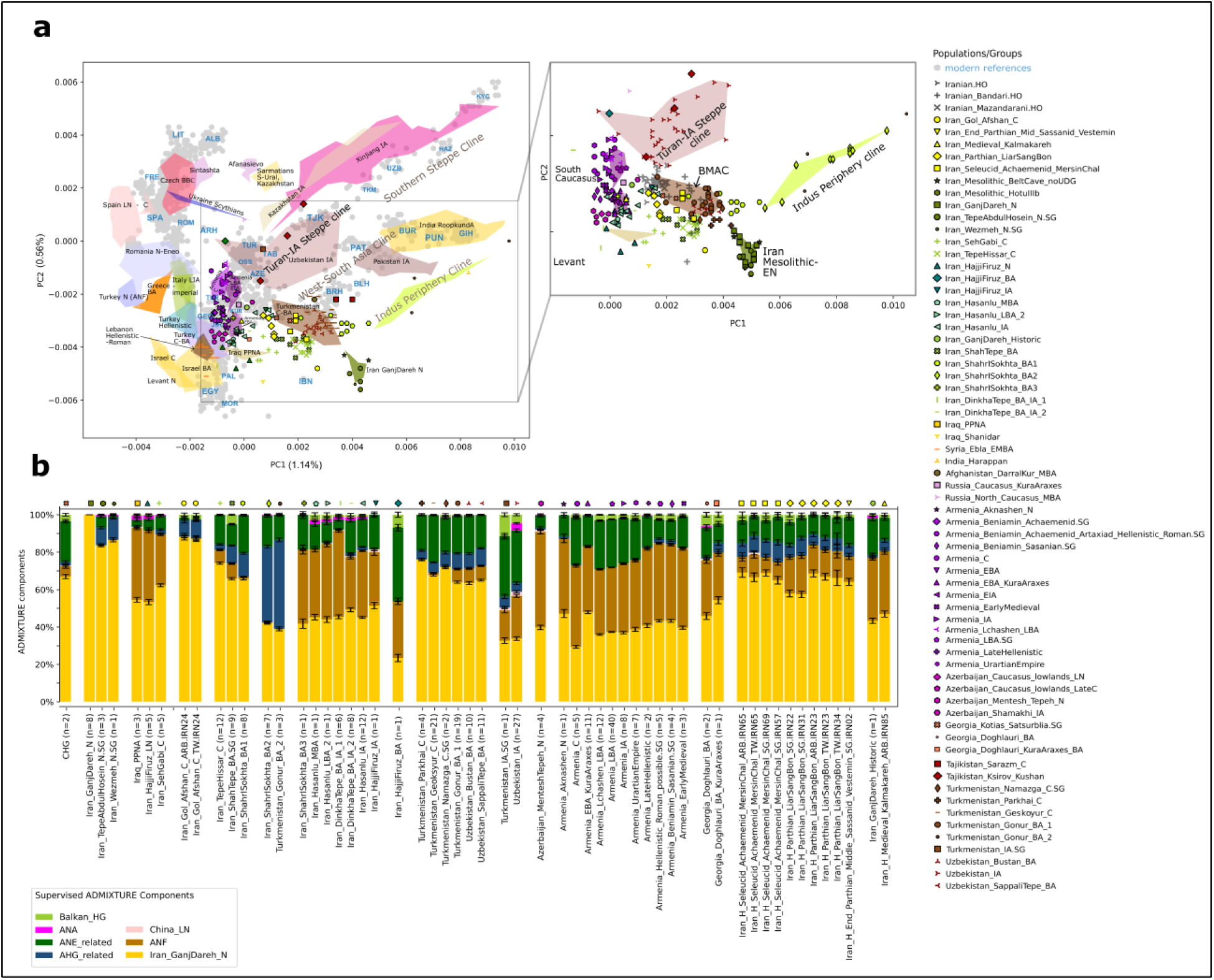
Eurasian PCA and supervised ADMIXTURE focusing on the genetic variation of Western Asia and its adjacent territories. (A) Grey dots represent modern reference populations, with some labelled in blue. Three-letter codes and detailed information on the ancient and modern populations used in the PCA plot are provided in Supplementary Table S9. A detailed view of the PCA is available in Supplementary Figure S11.2. (B) Supervised ADMIXTURE analyses of the most relevant samples and groups with Iranian EN Ganj Dareh, Anatolian Neolithic (ANF), Onge (AHG), WSHG and Botai (ANE-related), Shamanka Eneolithic (ANA), China LN, and Iron Gates HG (Balkan HG) as fixed sources. For more details and references, see Supplementary Table S10. Individual values and standard errors per component were used for the group-based representation, as described in Methods. Results of the full ADMIXTURE run are presented in Supplementary Figure S11.3.

Based on the observations by Davidson et al.^29^ regarding allelic bias of the Daicel Arbor Prime Plus capture data, we treated different capture data from the same individuals and groups separately, and used them to cross-check the allele sharing analyses. Since the Iranian and South-Central Asian reference samples consisted entirely 1240k or shotgun data, potential artificial affinities between Arbor Prime reference and target data were not expected. Considering the PCA, ADMIXTURE, and D and *f*_3_-*f*_4_ statistics conducted with our dataset, we conclude that Arbor and Twist captures did not introduce major discrepancies that could affect the interpretation of the results (Figures 3 and 4, Supplementary Figures S11.4-9, Supplementary Tables S11-12). However, differences were observed in the qpAdm models for these samples (Supplementary Tables S13-14).

**Figure 4.**
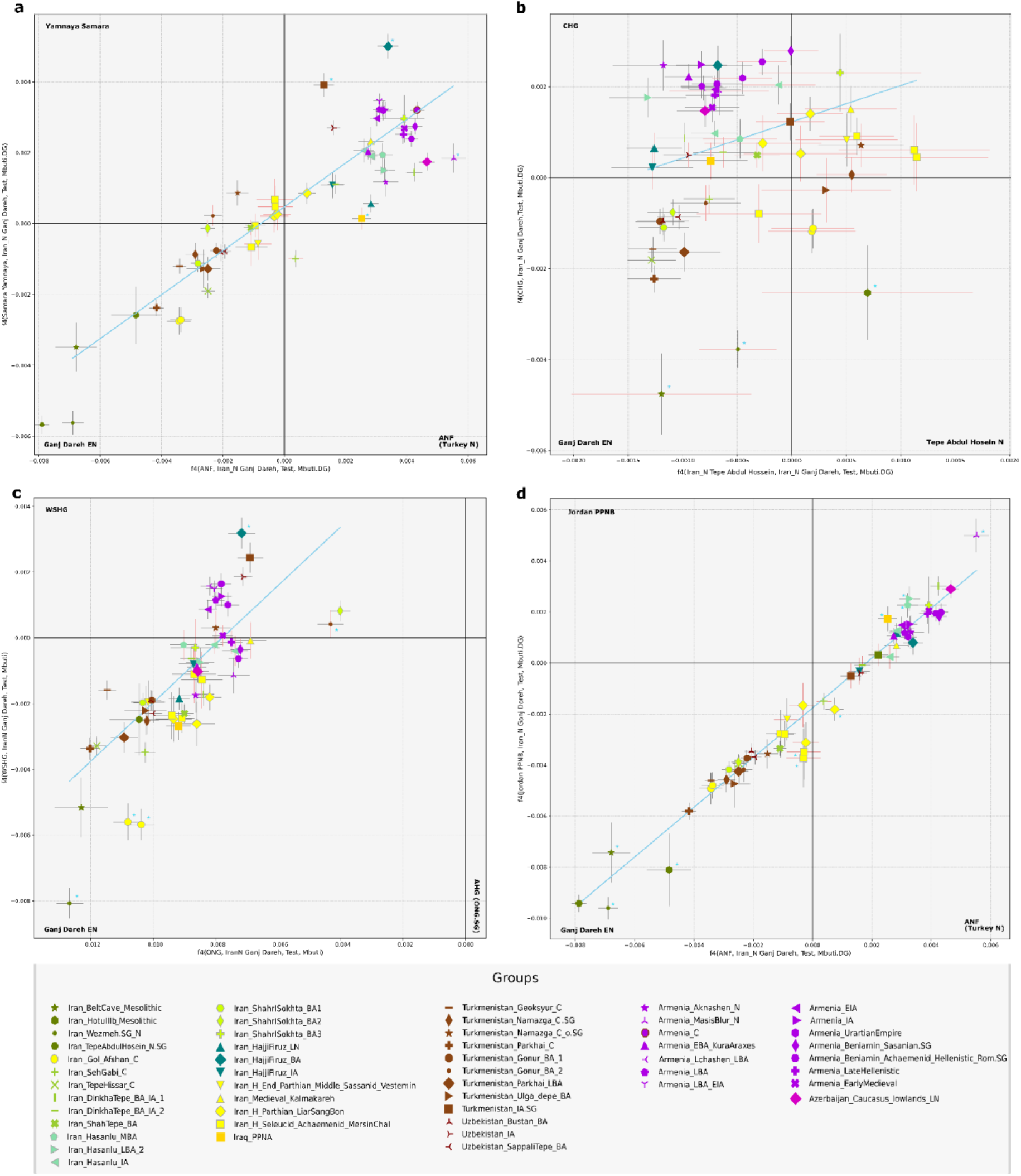
Two-dimensional scatterplots of seven f4-statistics. The four panels represent scatterplots of different f4 model combinations. f4 estimates and corresponding Z-scores are provided in Supplementary Table S11. Non-significant Z-scores are indicated with red error bars. Blue lines represent regression analyses (Weighted Least Squares (WLS) linear regression) and blue stars denote groups that deviate significantly from the regression, based on the model fit and the variability captured by the weighted standard deviation. Data produced using different laboratory methods from the same sample is indicated with a single symbol.

On the genomic PCA, we projected ancient samples onto a subset of the modern Eurasian dataset. We observe notable differences among ancient and modern groups inhabiting the areas within and surrounding the ancient Eurasian Silk Road, which also crossed the Iranian Plateau towards the Mediterranean^43^. Based on principal component 1 (PC1), we identify two distinct clines relevant to our study: (1) the Southern Steppe cline, stretching between Caucasus/Iran and the Central Asian populations^19^, and (2) the Indus the Periphery Cline, extending between Western Asia and Southern Asia^12^. Ancient and recent populations of the Iranian Plateau are clustered in the centre and bottom-centre of the PCA plot, along with South Caucasian groups (Figure 3, Supplementary Figure S11.2).

In the following section, we provide a comprehensive overview of the Iranian prehistoric and historic datasets, highlighting major genetic trends observed through various population genetic methods. The first two principal components position the Iranian Mesolithic and EN samples as distinct from all other ancient and modern samples, indicating their unique base composition, even separate from CHG, which has been previously discussed as cladal to EN Iran^13^. Previous studies have described EN groups (Tepe Abdul Hosein and Wezmeh) as homogenous with each other, using either CHG^13^ or EN Ganj Dareh as unadmixed sources for their modeling^12^. We consider both EN Ganj Dareh and CHG as suitable local sources for the subsequent populations of the Iranian Plateau. In supervised ADMIXTURE, we observe that CHG harbours the pre-defined Ganj Dareh-related and ANE-related components, aligning with Allentoft et al.^15^. We note, therefore, that some of the ANE-related component appearing in the Persian Plateau in our analyses may stem from CHG-related ancestry. In subsequent analyses, we tested the potential differences between EN Ganj Dareh and CHG, as well as the impacts of CHG-related components on the Plateau. We conclude that their contributions vary geographically but cannot be separately quantified within the current dataset (*f*_4_, qpAdm).

In the same supervised ADMIXTURE analysis, we observe that the ancestry composition of the Natufian and pre-pottery Neolithic individuals from the Levant is represented by 20-25% EN Ganj Dareh and 66-78% ANF ancestries (Supplementary Figure S11.3). This suggests that some Levantine ancestry present on the Persian Plateau might be masked within these components. We took these limitations in consideration during the subsequent analyses.

### Genetic structure of the Iranian Plateau during the Neolithic-Chalcolithic transition

We studied a new Early Chalcolithic male genome from the 5th millennium BCE Gol Afshan Tepe site, located on the eastern foothills of the Zagros^44^, which predates all other published Chalcolithic genomes from Iran^5,12^. Supervised ADMIXTURE indicates that he was a descendant of Early Neolithic farmers related to the populations of Ganj Dareh, Wezmeh and Tepe Abdul Hossein, all located in the central Zagros area (Figure 1). On the PCA, the Gol Afshan individual shifts toward more-western ancestry profiles. In *f*_4_-statistics, he still shares more genetic drift with EN Ganj Dareh than with CHG, LN Hajji Firuz, Chalcolithic Tepe Hissar or Chalcolithic Seh Gabi (Figure 4B, S11.5), indicating the EN Iranian component as a more likely source than CHG in this region. Among Neolithic Iranians, the pairwise allele sharing of the Gol Afshan individual with EN Ganj Dareh and Neolithic Tepe Abdul Hosein is not significantly different. Nevertheless, Gol Afshan shares the most alleles with Ganj Dareh and Tepe Abdul Hosein among any other Neolithic and Chalcolithic groups of the Iranian Plateau (Supplementary Figures S11.5). In line with the genetic results, substantial cultural continuity has been detected during the Neolithic-Chalcolithic cultural transition. The Bakun-period Gol Afshan Tepe site was used semi-sedentarily (Supplementary Information, Chapter 1). Furthermore, a ritual ceremony, recorded in the burial type, and the common practice of artificial cranial deformation, also observed at the Gol Afshan Tepe individual, find parallels at the EN Ali Kosh archaeological site in the central Zagros^45^. Moreover, contemporaneous with Gol Afshan Tepe, numerous instances of artificial cranial deformation and other cultural similarities have been recorded at the Chega Sofla archaeological site in the southern Zagros^44,46^ (Supplementary Information, Chapter 1). All these findings suggest long-term cultural and biological continuity in and around the Zagros region.

In supervised ADMIXTURE, we observe approximately 8-10% additional AHG component in the Gol Afshan and Tepe Abdul Hosein groups. This minor Neolithic variation, also identified using Tepe Abdul Hosein in the *f*_4_-statistics and shown in Figure 4B-C, suggests the presence of further ancestral pre-Neolithic or Neolithic genetic elements that warrant exploration in future studies. Statistics such as D tests in the form of D(Gol Afshan, Tepe Abdul Hosein, Shahr-i Sokhta BA1/2, Mbuti), comparing between the allele sharing patterns of the two Shahr-i Sokhta groups, suggest that this variation is not a gene flow from the Indo-Iranian borderland, but reflects unascertained genetic diversity within the prehistoric Iranian Plateau. This diversity appears to distinguish most of the eastern regional variation from the northwestern Plateau and the South Caucasus (Supplementary Table S11, Figure 3, Figure 4B-C). In subsequent analyses, we demonstrate that several two-way and three-way distal qpAdm models fitting Gol Afshan require AHG as a proxy source. However, Gol Afshan can also be modelled without the AHG-related component (Supplementary Table S13).

Besides the dominance of local EN continuity, the eastward expansion of Levant/ANF-related components is also observable in the mid-5th millennium BCE Gol Afshan individual^44^, in addition to previous evidence from northwestern Iranian Late Neolithic-Chalcolithic^12,13^. This individual exhibits a significant excess of allele sharing with ANF when compared to the Neolithic Iranian genomes from the same Zagros region (see D-statistics in Supplementary Table S11, Z=4.182). A series of qpAdm tests indicate additional ancestries to the EN Ganj Dareh component in Gol Afshan, such as ANF (modelled with a 17% contributions), or Neolithic Armenia (Masis Blur and Aknashen), LN Hajji Firuz and Chalcolithic Israel, all of which also show significant ANF ancestries^12,13^. Overall, these show the early admixture events of western influxes into the Neolithic Iranian gene pool. We capture with *f*_4_-statistics that every reference group from the Iranian Plateau shares more allele with ANF than with the Natufian and pre-pottery Neolithic Levant (Supplementary Figure S11.9, Supplementary Table S11). We also find that Iraq PPNA exhibits higher allele sharing with ANF than with pre-Chalcolithic Levant or EN Ganj Dareh (Supplementary Figure S11.9). Meanwhile, for the ancient groups of the Iranian Plateau and its surroundings, we demonstrate a correlational increase in allele sharing with the ANF-related ancestries (Turkey_N) and Levant-related ancestries (Jordan_PPNB) (Figure 4D). These findings indicate the influx of a profile combining both ancestries. This evidence aligns with the commercial interactions of Zagros inhabitants like Ganj Dareh, HajjiFiruz, and Bakun excavation areas with Levantine communities in Syria. These interactions are evidenced by the use of numerical tokens dating back to the Neolithic–Chalcolithic time horizon^47^. We acknowledge (based on Lazaridis et al.^13^ and our plausible qpAdm models) that the amount and direction of potential Levantine influx might be characterized better in future research, if additional high coverage comparative data become available.

The correlated allele sharing of the ancient Iranian and surrounding groups show regional patterns, regardless of the temporal differences, with the sharing with ANF and Levant components increasing from east to west (Figure 4D). Together with the supervised ADMIXTURE analyses, this demonstrates a long-standing East-West genetic cline of the Neolithic Iranian and ANF ancestries^13^. For instance, this is observable in the northern area through two Chalcolithic sites located more than 700 km apart and from different time periods. The Tepe Hissar group (ca. 3600-2000 BCE) exhibits lower ANF-related ancestry (∼7%) and higher EN Ganj Dareh component compared to Chalcolithic western Iranian group, such as Seh Gabi (ca. 4800-3700 BCE), which carries ∼27% ANF-related ancestry, as shown in supervised ADMIXTURE and *f*_4_-statistics (Figure 4A). This increase is also observed in the northwest Iranian sites from the LN-BA periods, such as Hajji Firuz, Hasanlu, and Dinkha Tepe, which harbour even higher levels of ANF-related ancestry than Tepe Hissar and Seh Gabi groups (Figure 3A-B).

### Genetic relations of Chalcolithic and Bronze Age groups of the Persian Plateau

The Mesolithic-EN ancestry forms a predominant basis in the Chalcolithic-Bronze Age genomes from the Iranian Plateau and South-Central Asia. The eastern Iranian BA Shahr-i Sokhta group 1 exhibits a high proportion of EN Ganj Dareh ancestry in its representatives (up to 65% in ADMIXTURE and *f*_4_) and an absence of ANF ancestry. The predominant EN Ganj Dareh related ancestry (∼70% in ADMIXTURE) in the earliest Chalcolithic samples from the Turkmenistan area (Parkhai, Namazga and Geoksyur, dated to ca. 3400-2800 BCE) suggest intense connections with the Iranian Plateau. The ANE-related ancestry observed through ADMIXTURE in these genomes aligns with the detection of these components in South-Central Asia by Narasimhan et al.^12^ and with our *f*_4_ statistics (Figure 4C). Archaeological evidence indicates interactions within the Persian Plateau as early as the EN, through the yet genetically unsampled Jeitun culture’s population, which occupied the territory of modern Turkmenistan^48^. Published Chalcolithic and BA groups of Turkmenistan (represented here by Geoksyur and Gonur sites) and Uzbekistan (Sapalli Tepe and Bustan sites) exhibit strong genetic similarities with the Chalcolithic Tepe Hissar, BA Shah Tepe, and BA Shahr-i Sokhta group 1 from Iran^12,14^. The Sapalli culture from Uzbekistan, which thrived from the first half of the 3rd to the mid-2nd millennium BCE, was closely related in material culture to BA Turkmenistan (Namazga, time periods V and VI) and Iran (Shahr-i Sokhta and Hissar III)^21^, reflecting the observed genetic homogeneity in the region. Meanwhile, ceramic traditions at Shah Tepe and Tepe Hissar demonstrate continuity from the Chalcolithic to BA periods^34^. Numerous specific artifacts —including soapstone beads, semi-precious stones, and various tools such as local blades, saws, and polished grey pottery—have been discovered at both Shahr-i Sokhta and Tepe Hissar sites, located in the northeastern and eastern Iranian Plateau. Subsequently, these artefacts were widely distributed across sites throughout the eastern and western Persian Plateau, suggesting that these two sites functioned as industrial centres facilitating cross-regional interactions^49–51^. Notably, Shahr-i Sokhta also exhibited increased population diversity, possibly coinciding or correlating with its role (Figure 3B, see Supplementary Information, Chapter 10).

### Stability in the Historical periods of the Northern Iranian Plateau

The CHG and EN Iranian ancestries became intermixed by the end of the prehistoric periods, consistent with observations across different periods^13,14,52–56^. These ancestries persist at levels of around 45-51% in the IA groups of the northwest Iranian Plateau (Hajji Firuz, Hasanlu, Dinkha Tepe sites) in our supervised ADMIXTURE analyses. Furthermore, they remain consistently predominant in the northern Iranian Plateau during the historical period, including the Achaemenid to Sassanid era burial sites of Marsin Chal, Liarsangbon and Vestemin. Most of these individuals can be modelled with CHG as a single source in both individual and group-based qpAdm models (including the combined Iran Historical group). However, the Liarsangbon individuals require an additional western component (16-26% ANF, Neolithic Armenia or Chalcolithic Israel). This finding is further supported by significant *f*_4_ results in the form of *f*_4_(Liarsangbon.SG, MersinChal.SG, ANF, Mbuti.DG) and aligns resembles the Liarsangbon site’s closer location to the western Iranian Plateau (Supplementary Table S11). Additionally, it also coincides with the cultural differences of Liarsangbon, evidenced by the discovery of non-local artefacts of Egyptian origin from the Roman times (Figures 3-4)^57^.

Evaluating other possible source populations, we demonstrate through *f*_4_-statistics in the form of *f*_4_(CHG, Test, Samara_EBA_Yamnaya, Mbuti.DG) and qpAdm models, that the BA Steppe affinities is only apparent due to shared CHG-related ancestries, which were previously defined in the BA Steppe communities^52^ (represented in our dataset with Samara_EBA_Yamnaya, Supplementary tables S11,13). The AHG-type ancestry detected in ADMIXTURE persists into the historical period. Moreover, in some deep ancestry qpAdm models of the historical individuals, the AHG (Onge) component reaches detectable thresholds.

After assessing the distal ancestry of the historical-period individuals (Marsin Chal, Liarsangbon, Vestemin), we modelled their proximal ancestries using a more focused approach. On the PCA, ADMIXTURE, and *f*_4_ analyses (Figure 3-4, Supplementary Figure S11.2-3, S11.6-8), these groups display ancestry patterns similar to those of the northeastern Iranian prehistoric samples, as well as to the BA Gonur and Sapalli Tepe from Turkmenistan and Uzbekistan. In accordance, the qpAdm models indicate a shared genetic ancestry of the historical period samples with the prehistoric northeast Iranians (Chalcolithic Tepe Hissar, BA Shah Tepe) or South-Central Asians (represented with Chalcolithic Geoksyur and BA Gonur group 1). Our analyses reveal that everyone out of the seven historical-period individuals yielding sufficient genome-wide data can be modelled either with one of these prehistoric groups as a single source or in a two-way model with an additional western Iranian source. In group-based analyses, shotgun genomes, such as those of the Marsin Chal and Liarsangbon groups (n=2 in each) can also be modelled with the same sources. However, the proportions of the northwestern IA components with elevated ANF/Levant-related ancestries are more prominent in Liarsangbon (Supplementary Table S14). IA Hajji Firuz is the only additional western source that provides plausible models for all historical groups. Using this source, Marsin Chal can be modelled with 78% BA Shah Tepe and additional ∼22% IA Hajji Firuz, while for the Liarsangbon group these amounts are 37% BA Shah Tepe and 63% IA Hajji Firuz. The combined group, referred to as Iran_North_Historical, produces similar results and can be modelled with 52% BA Shah Tepe and 48% IA Hajji Firuz-type contributions (Supplementary Table S14). We interpret these findings as evidence that the genetic profiles of historical-period groups of the northern Iranian Plateau reflect their position along the broader east-west genetic cline, rather than resulting from specific admixture events.

### Scattered evidence on Medieval and Modern Iranian Populations

The Medieval period individual from the Kalmakareh Cave in western Iran has a genetic profile similar to those of the northwestern groups of the Iranian Plateau, such as Hasanlu and Hajji Firuz, as well as a Medieval sample from Ganj Dareh, as observed both on the PCA and in ADMIXTURE. This, combined with *f*_4_ and distal qpAdm results, indicates a reduction of Iran Neolithic ancestry in the western Iranian Plateau over time. Meanwhile, most of the genetic ancestry of the newly published individual can be modelled in qpAdm with the mentioned IA variety in the Plateau, such as IA Hajji Firuz as the single source or as a mixture of IA Hasanlu (∼80%) and another source (e.g., 20% Shah Tepe from the northeast). The lack of published ancient individuals from the southwestern Iran limits further investigation into the regional genetic ancestries.

On the PCA, modern Iranians align with the majority of the Iranian prehistoric and historic genetic variation, but do not exhibit the Indus Periphery Cline characteristics previously described in the BA by Narasimhan et al^12^. In addition to clustering together in the PCA, the Mazandarani and Fars groups of Iran also show similar population models, albeit in different proportions, in the qpAdm analyses performed on the Human Origin dataset (Supplementary Table S14). Further Medieval and modern genome-wide sample pools are required to better understand the formation of the modern Iranian genetic variation.

## Conclusions

In this study, we presented novel ancient mitogenomes and genome-wide data from previously unstudied areas of the Iranian Plateau, with a particular focus on the northern area, southward of the Caspian Sea. We revealed several insights into the genetic history of the ancient Iranian populations and provided a comprehensive overview of the available ancient DNA data from Western Asia.

We explored the influence of major ancestry sources on the new dataset. We reproduced previous results on a prehistoric East-West cline of the EN Iranian ancestry with Neolithic Anatolian (ANF) and Levant-related autosomal ancestries. This cline exerted a lasting imprint on the population of the Iranian Plateau up to the historical period. The allele sharing with both ANF and Neolithic Levant increased towards the western end of this cline. We also discussed varying dual ancestry patterns of CHG and EN Ganj Dareh ancestries in ancient peoples of the Plateau. Furthermore, we found signals for a previously undescribed (AHG-like) ancestry in the Iranian Neolithic farmers that likely distinguished the Iranian Plateau’s population from more westerly groups, such as the contemporaneous South Caucasians. These observations indicate long-term genetic tendencies in the Iranian Plateau.

The new Early Chalcolithic genome from southwestern Iran presented in this study showed closer alignment with Early Neolithic Iranian farmers, with additional contributions from other Neolithic groups in western and northwestern proximities. This finding suggests predominant continuity, but also that the western Iranian region maintained contact with neighbouring areas, facilitating the introduction of western ancestries into the Iranian Plateau during the early stages of the Neolithic-Chalcolithic transition.

We demonstrated a strong Iranian Neolithic and CHG substrate in the historical-period samples from northern Iran, where these genetic components persisted in the pre-Medieval era. We confirmed the continuity from the Chalcolithic-Bronze Age into this period in northeastern Iran, despite this area hosting part of the Silk Road, which facilitated extensive human movement. Bronze Age Steppe ancestry remained relatively minor during the historical period in northern Iran. Instead, the historic period population of the northern Iranian Plateau exhibited strong genetic affinities with the Chalcolithic and Bronze Age communities of Turkmenistan, and northeastern-eastern Iran, forming homogeneous groups in our analyses as a part of the described east-west cline. As only one Iron Age genome is available from Turkmenistan, and there are none from the northeastern Iranian Plateau, further sampling is necessary to investigate the dynamics of this era, particularly to determine whether contacts between the two regions were sustained or disrupted after the Bronze Age.

Although only two Medieval genomes are currently available from the Iranian Plateau (one published in this study), the data indicate that the majority of the ancient Iranian gene pool remained stable over the centuries, with minor changes observed in the contemporary Iranian population. This suggests enduring genetic variation over millennia. Notably, at least half of the genetic heritage in Medieval southwestern Iran originated from Neolithic Iranian farmers, likely transmitted through Iron Age Iranians, despite the region’s exposure to external interactions with Mesopotamia, the Levant, the Caucasus and Anatolia. This highlights the genetic stability of the region’s inhabitants, even in the face of historical migrations and cultural shifts.

In the newly published dataset, we described Y-chromosomal and mitochondrial haplogroups that evolved around the ancient Persian Plateau, and which are still rare in ancient genome databases. We compared these with the available ancient and modern data and showed the long-term continuity in the uniparental ancestries in the region.

In summary, this research provides new evidence enhancing our understanding of the genetic characteristics and connections of ancient Iranian populations, while further comprehensive sampling is still required to uncover their internal diversity.

## Material and Methods

### Sampling strategy

In our genetic analyses, we examined samples from a variety of archaeological sites across Iran, representing a wide range of periods and cultures. We collected a total of 50 samples from nine different cemeteries and sites. From the earliest periods, we sampled from the Early Chalcolithic culture (Middle Bakun period) for the first time, which is represented by the Gol Afshan Tepe site, located in the southern Zagros Mountains^44^. The Late Chalcolithic through to the Late Bronze Age are represented in our dataset by samples from Gohar Tepe in the northern Iranian Mazandaran, Shahr-i Sokhta in southeastern Iranian Sistan and Baluchestan Province, and Cham Papi archaeological site in the southwestern Ilam Province of modern Iran. Details on these sites can be found in the Supplementary Information.

The Iron Age Neo-Elamite period (1100-539 BCE) is represented by samples from the Jubaji burial tomb in Khuzestan Province. Samples from the historical period originate from the Achaemenid to Seleucid periods (355-280 BCE) at Marsin Chal in Semnan Province. From the Parthian Empire (247 BCE-224 CE), the samples were collected from Liarsangbon in Gilan Province. The late Parthian to Middle Sassanid Empire periods (200-460 CE) are represented by samples from Vestemin in Mazandaran Province, all from northern Iran. Lastly, the Medieval Period (1215-1270 CE) is represented by samples from Kalmakareh cave in Lorestan Province (Supplementary Table S1, Figure 1).

### Sampling

For genetic investigation, we prioritised sampling petrous bones if they were available, due their favourable DNA preservation properties^58^. Alternatively, we collected teeth and long bones^59^.

### Radiocarbon dating

Radiocarbon dating was performed at the HUN-REN ATOMKI AMS C-14 facility of the Isotope Climatology and Environmental Research Centre in Debrecen, Hungary. Before Present (BP) dates were calibrated in Oxcal software v4.4, using IntCal 20 calibration curve^60,61^ (Supplementary Table S1).

### Ancient sample preparation and DNA extraction

All steps of sampling, DNA extraction, and library preparation were carried out in the dedicated ancient DNA facility of the HUN-REN RCH Institute of Archaeogenomics. After thorough mechanical surface cleaning and UV-treatment, we drilled the samples to gain powder from them^62^, exception of four samples that were ground in a mixer mill. An additional three samples were processed with the minimally destructive protocol developed by Harney et al.^63^. DNA was extracted from 50 mg bone powder either manually^64^ or using a liquid-handling automated workstation^65^. As for the three minimally destroyed samples, 150 μl lysate was used for DNA extraction. Three very short fragments of the mitochondrial DNA (mtDNA) were amplified with primers from the GenoCoRe22 assay^66^ in order to verify the success of DNA extraction.

### DNA library preparation, hybridisation captures and sequencing

We prepared half-UDG treated, double-stranded libraries with unique barcoded adapters following a well-established protocol, incorporating minor changes for automation^67^. Libraries were amplified with TwistAmp Basic (Twist DX Ltd), followed by purification with AMPure XP beads (Agilent). Finally, unique indexes were added to each library in a PCR reaction for multiplex sequencing^68^.

In some cases, we carried out hybridization capture to selectively enrich specific targeted regions of the genome. Mitochondrial genomes were captured as described in Csáky et al.^69^, while genomic variants were detected with myBaits Arbor Expert Human Affinities “Complete” Kit (Daicel Arbor Biosciences, which includes the ancestral 850K SNPs set in addition to the ones from “Prime Plus” (1240k, Y, and mitochondrial SNPs). After concerns were raised about the Prime Plus portion of the Arbor capture^29^, we repeated the most important samples with Ancient Human DNA Panel Workflow (Twist Bioscience^30^).

Samples that passed the screening sequencing on an Illumina MiSeq system with Reagent Kit v3 (2 × 75 cycles) were later submitted to Novogene Ltd for deep sequencing on NovaSeq 6000 platform, using 2 × 150 cycles.

Criteria for deeper shotgun sequencing were >40% endogenous DNA content and <2% duplicated reads mapped on the genome. Criteria for genomic capture were 4%-40% endogenous DNA detected and <5% duplicated reads at the shallow shotgun screening (Supplementary Table S2).

### Bioinformatic analysis of the NGS data

We used the same in-house pipeline for raw sequence data processing as described in Gerber et al. ^70^ using PAPline (https://github.com/ArchGenIn/papline) (for Supplementary Table S1). The ancient data as BAM and FASTQ files were uploaded to ENA, under the project number PRJEB81975 (https://www.ebi.ac.uk/ena/browser/home). The GRCH37.p13 reference sequence was used to call the pseudohaploid genomes.

We used BREADR package (Biological Relatedness from Ancient DNA in R) to assess kinship among the samples^32^, for results, see (Supplementary Table S8). Haplogroup determination for mtDNA was performed by HaploGrep 2 on FASTA files that were called with a custom R script tailored for archaic DNA^71^. Using the deep-sequenced genomes, we determined the Y-chromosomal haplogroups with Yleaf v1 software^72^.

Contamination levels of the mtDNA were estimated using the ContamMix software^73^ X-chromosomal contamination in male individuals was measured with two software tools: ANGSD (version 0.939-10-g21ed01c, htslib 1.14-9-ge769401) and hapCon (hapROH package version 0.60)^74^. The doCounts for the former method was run on X-chromosomal region from positions 5500000 to 154000000, with base quality 30 and mapping quality 25. The toolkit’s contamination executable was used to process the resulting file. Parameters were set to -b 5500000 -c 154000000 -d 3 -e 100 -p 1. Additionally, the file specified by the -h parameter was included as provided by the software developers. For results, see Supplementary Table S3 and S4.

Runs of homozygosity were calculated with the hapROH program^33^, run with default parameters for all pseudo-haploid genotypes with at least 150k SNP covered from the 1240k SNP panel, and repeated for the merged BAM files including libraries captured via the two above-detailed methods (Supplementary Table S13).

### Population genetic analyses

We performed various population genetic analyses, in which we compared the studied populations to several other ancient and modern-day published datasets. We compiled AADR v54.1^39^ with further relevant studies from Western and Central Asia^14,56,75–82^.

CHG group was created by combining KK1_noUDG.SG and SATP_noUDG.SG^83^ EHG group was created with a combination of I0061, I0211, I0124, Sidelkino_noUDG.SG^28,84^. China_LN was prepared as a combination of the China_Upper_YR_LN and China_YR_LN groups^85^. Levant_preChalcolithic included Jordan_PPNB and Israel_Natufian^13^. Armenia_N was a combination of I1390 and I1391^13^ West_Siberia_N_Botai was a composite group including the individuals BOT14, BOT2016, I1960, I5766^12,19^.

The Principal Component Analysis (PCA) was computed using the EIGENSOFT smartpca software (v16000)^86^ with the Human Origins Panel SNP set^39^ and a reduced set of West-Central Eurasian modern populations for calculation, as listed in Table S9. For every other analysis, the 1240k array SNP set^28^ was used for variant calling. To estimate ancestry composition, first we used supervised ADMIXTURE analysis (K=7) calculated by the ADMIXTURE v1.3.0 software^42^. The seven fixed ADMIXTURE source components were chosen depending on their previously described genetic impacts on the Eurasian continent (details of the sources are in Table S10)^12,19,66,85,87,88^. To present the standard error bars of the components in Figure 3B, the standard errors for each component of the individuals of the ADMIXTURE dataset were calculated using point estimation and bootstrapping, as described in the official ADMIXTURE manual. We then calculated mean component values for the groups and the group standard errors with the formula= (√(SE1²+SE2²+…+SEn²))/n, where “n” represents the number of samples in each analysis group. *f*-statistics and qpAdm were performed using AdmixTools v7.0.1^40^ and ADMIXTOOLS v2^41^.

Data analysis and visualization of the *f*-statistics was performed in Python using pandas (data handling), numpy (numerical operations), matplotlib (visualization), and statsmodels (regression analysis). Weights were calculated as the inverse of the combined variances to account for measurement precision. Weighted Least Squares (WLS) regression was used to model the relationships between *f*_4_ variables, with weights assigned inversely to measurement variance, prioritizing more precise data points. A constant was added to the predictor variable to include an intercept. Slope and intercept were extracted, and residuals were calculated to evaluate prediction accuracy. Outliers were identified on the *f_4_* plots based on a weighted standard deviation of residuals, with a threshold set at twice this standard deviation. Points exceeding this threshold were flagged as significant deviations. Scatter plots with error bars, color-coded by Z-scores, were created to visualise each comparison and highlight outliers.

To elucidate the ancestries on a global and regional bases, we tested deep and proximal ancestry qpAdm models for the newly analysed dataset, where we considered only the individuals with at least 100,000 SNP covered on the 1240K panel. For the distal qpAdm models, the right group set consisted of Mbuti.DG, Papuan.DG, China_LN, Luxembourg_Loschbour.DG, Russia_MA1_HG.SG, EHG, Morocco_Iberomaurusian, Turkey_Epipaleolithic, Levant_preChalcolithic. In the proximal qpAdm models, we used Mbuti.DG, ONG.SG (Onge), Caucasian HG (CHG), EHG, Turkey_N, Jordan_PPNB, Russia_MA1_HG and Serbia_IronGates_Mesolithic as right groups. These populations in the two right group sets, have diverse genetic relationships to the ancient populations of the Iranian Plateau and its surroundings, as well as to the tested left group sets, which are expected to enhance the resolution of qpAdm analyses^89^. We refrained from rotating the left and right group populations, due to severe criticism raised in recent research^90^. For the qpAdm analyses we used the qpadm_multi function of ADMIXTOOLS v2 with the parameter allsnps: NO (as allsnps=FALSE as an argument of the qpadm_multi function). Group-based proximal qpAdm models were run for the historical-period Liarsangbon and Marsin Chal shotgun genomes. In addition, the Iran_North_Historical group was created using these individuals and the historical period Vestemin individual. Our in-house script for qpAdm analyses considered only the results with ≤0.2 (20%) standard error per source component in the model. In addition, results with standard error higher than the source proportion, or those with a negative value for at least one source proportion were deemed unfeasible. Feasible models in distal analyses are presented in Table S13. Unfeasible models in proximal analyses are presented with p-value=0 (no decimals) and NA in source proportion columns (Supplementary Table S14). When evaluating qpAdm models, we accounted for high false-positive rates reported recently^90^, and chose the most likely models based on the PCA, ADMIXTURE and *f*-statistics. Between two-way and three-way models, we preferred the lower-ranking ones.

## Institutional Review Board Statement

Ancient samples were taken with the written consent of the stakeholder museums and the excavator and processor scholars. Human remains were treated with dignity and respect, to ensure minimal destruction of the samples.

## Data Availability Statement

The ancient mitogenome and shotgun genomic data presented in this study will be openly available in the European Nucleotide Archive at (https://www.ebi.ac.uk/ena/browser/search) upon publication, with accession number (PRJEB81975).

## Acknowledgement

We thank Krisztián Oross for his helpful comments on radiocarbon dating. Our sincere appreciation is extended to Dr. Seyed Mansur Seyed Sajjadi, Dr. Arman Shishegar and Dr. Lili Niakan from the Iran Center for Archaeological Research (ICAR), Dr. Ali Mahfroozi and Serollah Ghasemi from the Mazandaran Research Institute for Cultural Heritage and Tourism, Ms. Somayeh Chavoshi from the Noor Human Genetic Research Center and Ms. Beheshteh Nasiri Rad from Ministry of Cultural Heritage, Tourism and Handicrafts for generously providing samples for this research. We express special thanks to Dr. Jebrael Nokandeh, Director of the National Museum of Iran, and Mr. Javad Nasiri for accessing valuable museum collections. Moreover, we thank Mr. Rohollah Zarin Pour from the Falak-ol-Aflak Museum for accessing valuable museum collections. We extend our gratitude to Prof. Kamalodin Niknami and Prof. Hassan Karimian from Tehran University for their expertise in Iranian Archaeological contexts and Mr. Farzad Frouzanfar from the ICAR for his insightful guidance throughout the anthropological endeavor. We extend our appreciation to Ms. Parastoo Erfanmanesh and Mr. Jalil Golshan from the Research Institute of Cultural Heritage & Tourism. Their tireless advocacy and outreach efforts were crucial in raising awareness about this type of research. We also extend our appreciation to Miss Forough Taheri from the Metabolic Disorders Research Centre, Endocrinology and Metabolism Molecular-Cellular Sciences Institute, Tehran University of Medical Sciences, Dr. Raofian and Dr. Danesh from the Legal Medicine Organization of Iran, Mr. Alireza Abolfazli, Director of ARA Electronic Company, Mr. N. Atighech from the Student Affairs Organization of Iran, Ms. Masoumeh Heidarian, Ms. Norasteh Abolfazli and Mr. Hamid Abolfazli from the Cultural Heritage, Ms. Masoumeh Sabri from the Iranian Red Crescent Society, Mr. Sadegh Taheri PhD Student, Department of Animal Science, Ferdowsi University of Mashhad, Dr. Sepideh Maziar from Institute of Archaeological Sciences, Dept. of Near Eastern Archaeology Goethe University, Frankfurt and Mr. Mehdi Amjadi for their support. We are thankful to the Research Institute Of Cultural Heritage and Tourism and the National Archaeological Research Institute for facilitating the acquisition of the National Legal Permits to work on ancient skeletal human remains.

## Author contribution

Conceptualization: MAA, MR and ASN; Methodology: MAA, MR, ASN and BGM; Collecting the samples: MAA, MR, AB, ZS, MHT, PZ, AH, VJ, AMS, and MT; Performed the DNA extractions and library preparation: MAA and MM; Conducted the bioinformatics analyses: MAA, YCO, KJ, ZH, and ASN; Performed the statistical analyses and data interpretation: MAA, YCO, MR, and ASN; Resources: MAA, MR, AB, MHT, HM, PZ, AH, VJ, AMS, BE, MT and ASN; Visualization: MAA and ASN; Writing – original main draft of the paper: MAA, YCO, MR, and ASN; Writing – review & editing: MAA, YCO, MR, and ASN; Funding acquisition: BGM, MR, MT, and ASN; Supervision: MR, MT and ASN.

## Abbreviations

AHG: Andamanese Hunter-Gatherer
ANE: Ancient North Eurasian
ANF: Anatolian Neolithic Farmer
BA: Bronze Age
C: Chalcolithic EN: Early Neolithic
LN: Late Neolithic
H: Historical period
HG: Hunter-Gatherer
IA: Iron Age
mtDNA: mitochondrial DNA
N: Neolithic
PCA: Principal Component Analysis
WSHG: Western Siberian Hunter-Gatherer

