## Supplementary texts and figures for "Ancient DNA indicates 3,000 years of genetic continuity in the Northern Iranian Plateau, from the Copper Age to the Sassanid Empire"

1: Doctoral School of Biology, ELTE-Eötvös Loránd University, Budapest, Hungary
2: Institute of Archaeogenomics, HUN-REN Research Centre for the Humanities, Budapest, Hungary
3: Noor Human Genetic Research Centre, Tehran, Iran
4: Department of Archaeology, University of Tehran, Tehran, Iran
5: Faculty of Management and Financial Science, Department of Management, Khatam University, Tehran, Iran
6: Université Lumière Lyon 2, Laboratoire Archéorient; Maison de l'Orient et de la Méditerranée, Lyon, France
7: Iranian Centre for Archaeological Research (ICAR), Tehran, Iran
8: Department of Biology, Science and Research branch, Islamic Azad University, Tehran, Iran
9: Research Institute for Cultural Heritage and Tourism, Lurestan, Iran
10: Research Institute for Cultural Heritage and Tourism, Gilan, Iran
11: Research Institute for Cultural Heritage and Tourism, Sari, Iran
12: Department of Genetics, ELTE-Eötvös Loránd University, Budapest, Hungary
13: Kawsar Human Genetic Research Centre, Tehran, Iran
14: MTA-BTK Lendület 'Momentum' Bioarchaeology Research Group, Budapest, Hungary

#### Index


An overview of current research related to genetic samples from prehistory to the Sassanid Period 3

Archaeological and anthropological context of the samples 4

- Chapter 1: Gol-Afshan Tepe site: Stratigraphy and Excavations 5
- Chapter 2: Shahr-i Sokhta cemetery 10
- Chapter 3: Gohar Tepe 14
- Chapter 4: Cham Papi Cemetery 18
- Chapter 5: Kalmakareh cave 20
- Chapter 6: Jubaji site 26
- Chapter 7: Marsin-Chal cemetery 34
- Chapter 8: Liār-Sang-Bon site 49
- Chapter 9: Vestemin 61

Supplementary information to the genetic analyses 74

- Chapter 10: Reconciling genetics and archaeology from a uniparental perspective 74
- Chapter 11: Supplementary Figures and notes to the whole-genome based analyses 80

References 89

#### An overview of current research related to genetic samples from prehistory to the Sassanid Period

The Chalcolithic (also known as the Copper Age) of Iran covers the period from the middle of the fifth millennium BCE to the end of the fourth millennium BCE. In general, it encompasses the transition from the simple social structures of the Neolithic period to a phase of increasing social and cultural complexity, marked by changes such as permanent settlement, agriculture, animal husbandry, economic progress, the development of commercial networks, and the emergence of specialised work tasks.

The Bakun culture is one of the most important cultures of the Chalcolithic of Iran (from the fifth to and early fourth millennium BCE), representing an important stage in the social and economic changes of prehistoric societies in the Fars province within the Persian Plateau (Alizadeh, 2006).

Moving to historical times, the Achaemenid/Persian Empire stretched from Anatolia and Egypt across Western Asia to the western side of the Makran mountains and South-Central Asia. This empire, which was founded in 550 BCE by Cyrus II (“the Great''), consisted of a vast territory and a large population. This empire included various ethnic groups consisting of leaders of different Iranian ethnicities such as Persian, Median, Elamite, and Scythian, who lived within the empire, handing over power to the king himself, while the power of local governments was subject to him and in charge of managing both internal and external forces.

After Alexander the Great's attack in 330 BCE, very little change was seen in the Achaemenid government, such that some historians refer to Alexander as the last Achaemenid king. After the death of Alexander in 323 BC, the Achaemenid dynasty officially ended, and the Seleucid dynasty began. During the Seleucid era, a mixture of Greek and Iranian culture was formed in our focus territory. During this period, although the people of the Iranian Plateau were defeated by the Greeks, they kept their former lifestyles and ways of living (Brosius, 2020).

In 247 BCE, the Parni tribe (Aparni) who were in the northeastern region rebelled against the Seleucids and established a separate state. The Parthian state was established under the leadership of Arsaces (Arshak) in the city of Nisa, close to the modern Iran-Turkmenistan border.

They created an empire that lasted for 476 years. It took over a century following the Arsaces uprising for the Seleucids to be completely expelled from Iran. This prolonged conflict was due to the Seleucid government's persistent efforts to maintain its rule over the region, leading to numerous battles with the Parthians. The Parthians gradually advanced from the eastern regions of Iran towards the centre of the country and after a series of wars, they dominated the western regions of Iran, i.e. the mountainous regions of Zagros. Seleucia, the former Seleucid capital, became the seat of the Parthian government around 141 BCE (Ahmadi Vastani & Ebdal Mahmoodabadi, 2020). The Parthians, renowned for their equestrian skills and archery, were often referred to as Pahlavan/Pahlav/Parthua, a term still used today (Sharifi et al., 2023).

The Sasanians, who arose from the Fars province in southern Iran from the land of Persia, in 224 CE, presented themselves as the unrivalled successors of the Achaemenids, and for more than four centuries, the Sassanid government was one of the two great governments of the civilised world at that time in Western Asia. Its borders in the east extended as far as the Indus River Valley and Peshawar, and in the northeast, it was sometimes extended to Kashgar city in southwestern Xinjiang, China. By overthrowing the Parthian monarchies, they established a national dynasty based on the two foundations of unity and centralization of religion and government. Ardeshir defeated the Parthian king and sat in his place. After his death, his son Shapur was crowned in 241 CE. Shapur first turned his attention to the Kushans and succeeded in destroying the Kushan state with a detailed campaign, and then went to the west to recover Mesopotamia and Armenia from the Romans. During the reign of Shapur II, Sasanian Iran reached the peak of its power. The concept of Iran as a geo-political term was accepted by historians during the ruling of Shapur II. The Sassanids ruled Iran for 427 years finally falling in 651 CE (Daryaee, 2023).

#### Archaeological and anthropological context of the samples

In this research, nine sites and cemeteries were studied, which include:

1. Gol-Afshan Tepe, Semirom, Isfahan province: Under the supervision of Dr. Mohammad Hossein Taheri, University of Tehran, Department of Archaeology, Tehran, Iran and Université Lumière Lyon 2, Laboratoire Archéorient; Maison de l'Orient et de la Méditerranée, France.

2. Shahr-i Sokhta, Zabol, Sistan and Baluchistan province: Under the supervision of Dr. Seyyed Mansur Seyed Sajjadi and Dr. Hossein Moradi, Iranian Centre for Archaeological Research (ICAR), Tehran, Iran.

3. Gohar Tepe, Behshahr, Mazandaran province: Under the supervision of Dr. Ali Mahofroozi,

Centre of Research, Office of Cultural Heritage, Handicrafts and Tourism Organization of Mazandaran, Iran and his colleagues Serollah Ghasemi Gorji, PhD student, Faculty of Art and Architecture, Mazandaran University, Babolsar, Iran.

4. Cham Papi, Tangeh Kafari- Rue River, Ilam Province: Under the supervision of Dr. Lili Niakan, Academic Member of the Iranian Centre for Archaeological Research (ICAR).

5. Kalmakareh cave, Pol-e Dokhtar- Rumeshkan- Lorestan province, under the supervision of Dr. Ata Hasanpour, Centre of Research, Office of Cultural Heritage, Handicrafts and Tourism Organization of Lorestan, Iran.

6. Jubaji, Ramhormoz, Khuzestan province: Under the supervision of Dr. Arman Shishegar, Assistant Professor, Iranian Centre for Archaeological Research (ICAR).

7. Marsin-Chal, Talajim, Semnan province: Under the supervision of Dr. Ata Hasanpour, Centre of Research, Office of Cultural Heritage, Handicrafts and Tourism Organization of Lorestan, Iran.

8. Liār-Sang-Bon site, Amlash, Gilan province: Under the supervision of Dr. Vali Jahani, Centre of Research, Office of Cultural Heritage, Handicrafts and Tourism Organization of Gilan, Iran.

9. Vestemin, Kiasar, Mazandaran province: under the supervision of Dr. Abdol Motalleb Sharifi, Centre of Research, Office of Cultural Heritage, Handicrafts and Tourism Organization of Mazandaran, Sari, Iran.

The Research Institute of Cultural Heritage and Tourism of Iran (RICHT) has authorised all excavations conducted at the listed archaeological sites from Iran. The directors of the excavation teams provided the archaeological information for the excavated areas studied in this research. We examined samples representing a wide range of periods and cultures. Archaeological information of only those graves, which have been studied for ancient DNA and dating purposes in this research, is summarised below.

#### Chapter 1: Gol-Afshan Tepe site: Stratigraphy and Excavations

The main information of this chapter was collected by the excavator of Gol-Afshan Tepe, Dr. Mohammad Hossein Taheri.

##### 1.1. Geographic location and general description

Named after the adjoining village, Gol-Afshan Tepe is in the Ghabr-e Keykha plain of the southeastern Semirom County, on the fringes of Isfahan Province, at an altitude of 2344 m above sea level. Its geographical coordinates are 34.5708 °N latitude and 57.5204°E longitude. Previous surveys have reported a high concentration of the distinct black and buff pottery and lithic tools of the Bakun Period (5000-4000 BCE) from the surface of the site.

##### 1.2. Trench E

A total of four trenches were excavated in this area, and excavation continued to virgin soil in trenches A and B, and excavation stopped in trenches D and E at the third settlement phase before reaching virgin soil.

Trench E was completed in the eastern quarter of the mound as a north-south 5 x 5 metre square. The four sides of the trench were conventionally designated as A, B, C and D respectively. It was opened to clarify the stratigraphy of the eastern part of the site.

In Trench E (Supplementary Figure 1.1.), 35 Locus and 3 occupation phases were distinguishable. Phase III, the earliest recognized phase, includes Floors 023, 025 and 026. A metal awl, grinding stones, numerical tokens, and a burial were found in this phase. Phase II was defined by Floor 032 and Walls 010, 017, 018, 007. The three occupation phases identified in this trench were represented by immediately superimposed deposits without any discernible hiatus in-between. The severe disturbance of the mud-brick walls made it impossible to draw complete architectural plans and barring a few scattered walls, the structures were absolutely demolished. A major find made in this trench was a deformed skull belonging to an approximately 30-year-old male, placed in front of the stone-lined fireplace beneath a mudbrick wall.

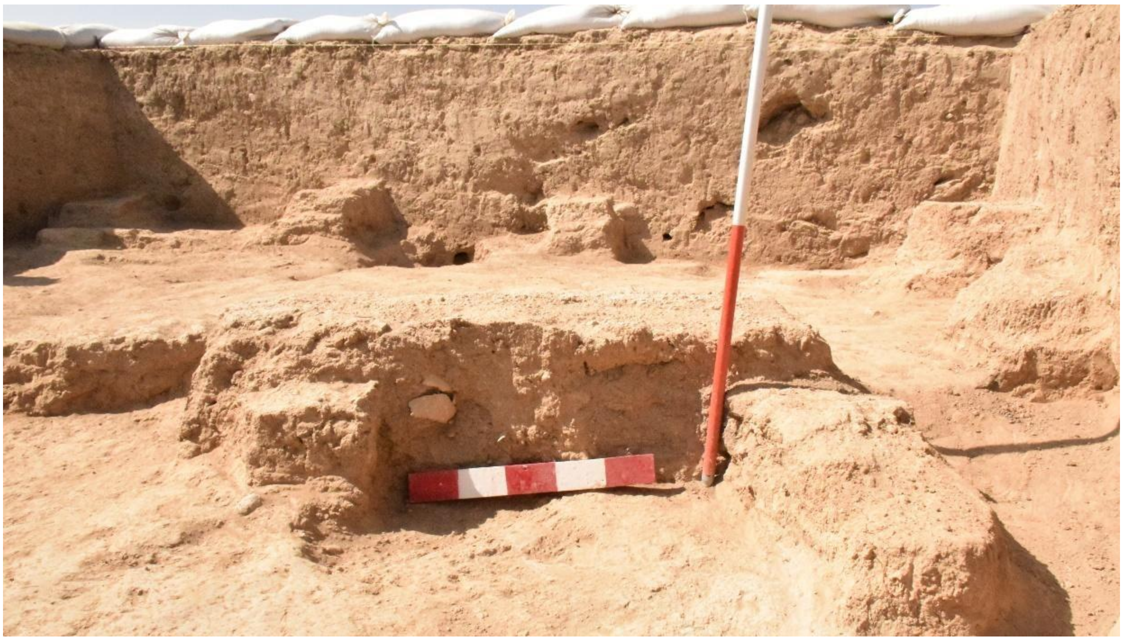

**Figure 1.1. Location of the skull in Trench E**

##### 1.3. Human skeletal remains from Gol Afshan Tepe

From Gol-Afshan Tepe we sampled an artificially deformed cranium of a young male (identified genetically and anthropologically, sample IRN24), aged between 25 and 35 years old. It was recovered in Trench E Locus 031, which was a grave that contained this deformed and incomplete human skull that completely lacks the left mandible and maxilla and is associated with a lumbar vertebra. The direct 14C radiocarbon date which was conducted in the Archéorient laboratory at the Maison de l'Orient et de la Méditerranée à Lyon (France) for phase II of Trench E, confirms the dating of the burial to 4690-4360 cal BCE (Taheri, 2022), corresponding to the Bakun Period.

At the time of the excavation, the skull was positioned face-up, with the orbital bone located at the same layer as the start point of Wall 018, while the cervical vertebra and occipital segments were under Wall 018. The Locus was sealed by Locus 007 and 003 and was adjacent to Locus 030. Probably, sometime after the end of Phase II (related with Floor 032), the residents of Phase I created a burial within the boundaries of Trench E on the southern side of the mudbrick Wall 007, also truncating part of Floor 032. Once the skull was interred in Locus 031, the upper part of Locus 007 was built using mud bricks as an attempt to cover the skull. To bury the skull, the area beneath Wall 018 was cut and the upper layer of Locus 007 was created, immediately following the internment, as the skull’s frontal bone and orbits lay within this layer.

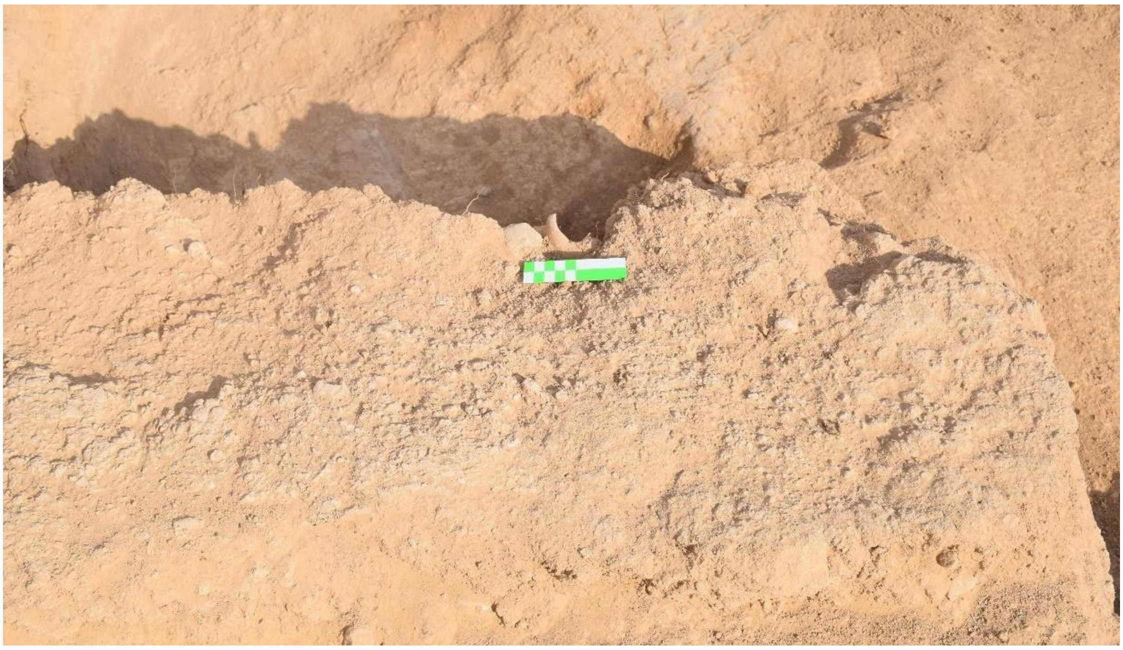

**Figure 1.2. Location of the skull in Trench E, and the position of the skull facing upwards**

##### 1.3.1. Artificial or cultural modification of the cranium of Gol-Afshan Tepe human individual

This section was drafted by Dr. Maryam Ramezani. The anthropological studies on the individual of Gol-Afshan Tepe was also conducted by Dr. Maryam Ramezani.

The term "artificial or cultural modification" refers to practices that alter the cranium's shape during infancy and early childhood. Since the skull is highly malleable during the early years of life, a desired shape can eventually be achieved through manipulation. Most cultural deformational procedures begin within the first few days of life, and the deforming apparatus is used for approximately six months to one year. However, some societies, such as Ecuador and Peru, may employ this process for three to five years (Torres-Rouff 2002). Although both genders can be the subject of cranial modification, the frequencies vary among societies. The methods used to induce cranial modification differ, depending on cultural practices and techniques.

Two methods have been commonly used: 1) Annular form: In this way, the anterior and posterior of the head are extended and lengthened by wrapping bandages or by a tight hat, producing a cylindrical or conical head shape. Such an extension influences the frontal, temporal, parietal, and occipital bones, creating cranial tapering on both sides. 2) Tabular forms: This method involves a more directed pressure resulting from the use of boards or stiff

pads on the anterior and posterior of the skull, resulting in a broader and shorter cranial shape (Torres-Rouff 2002).
Gol-Afshan Tepe skull (Supplementary Figure 1.3) exhibits an Annular form in which these cultural modification practices are the same as observed at Chega Sofla (Supplementary Figure 1.5) and Ali Kosh excavation sites.

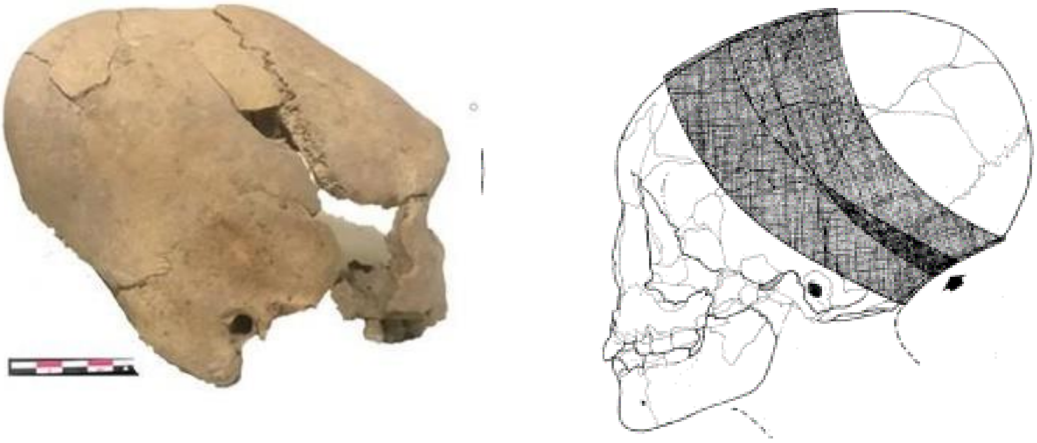

**Figure 1.3. left: Deformed skull from Gol-Afshan Tepe right: The method of deformation of the Byblos skull proposed by Özbek (Özbek, 1974).**

Although some anthropological study back to 2011 shed light that some individuals like the skull of a female from Ali Kosh may not be certainly considered as a deliberately deformed cranium, still this tradition certainly existed in Ali Kosh (Supplementary Figure 1.4) (Niknami et al., 2011).

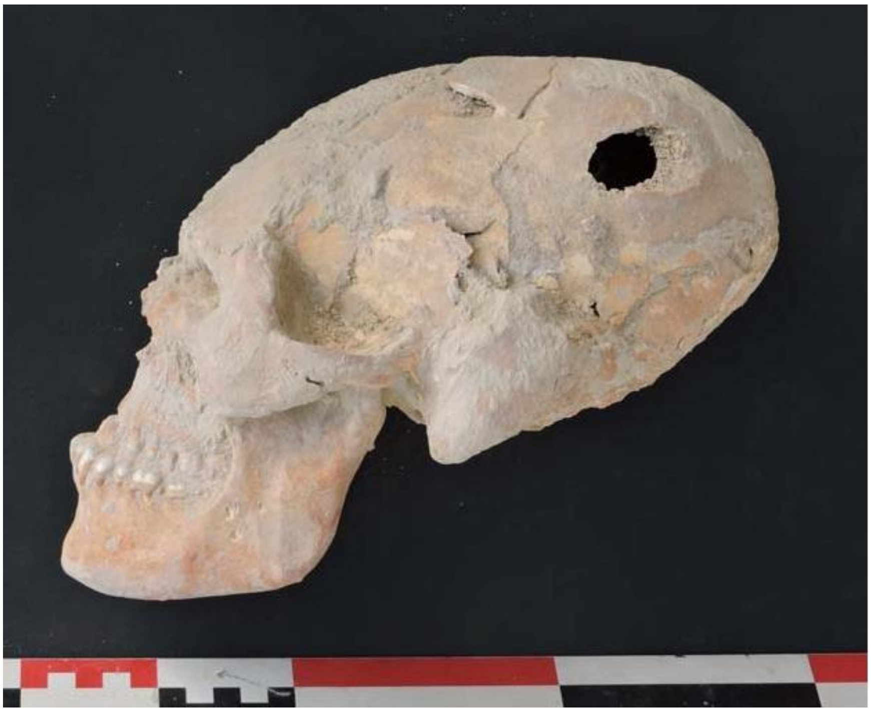

**Figure 1.4. Cranial deformation from Ali Kosh, Ilam, province, Iran (**Sołtysiak & Darabi, 2017).

Overall, there are similarities in the methods and traditions of skull modification between the Zagros and Southwest Asia. It is worth noting that prehistoric skulls modified in Southwest Asia are of the Annular form, in contrast to other parts of the world, such as North or South America, where the skull modification typically follows the Tabular form. According to a published report in Southwest Asia, boards were not used in the modification process (Niknami et al., 2011).

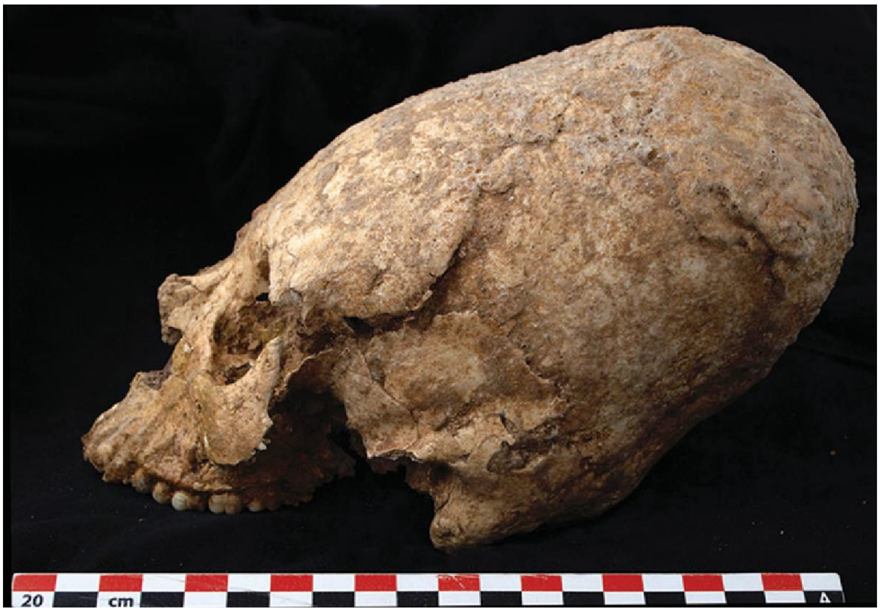


**Figure 1.5. Deformed skull from Chega Sofla, Khuzestan, Iran (Alirezazadeh-Nodehi et al., 2024).**

The oldest known evidence of cranial modification was discovered at the Zhou-Kou-Tien cave in China, and it dates back to between 23,000-18,000 BC of the Upper Paleolithic era (Soto-Heim 1986). In the Near East, the tradition can be traced back to the Neanderthals of the Shanidar cave in Iraqi Kurdistan. The most important Near Eastern cranial modifications are those from Neolithic sites in Jericho (Palestine), Ganj Dareh, Ghenil Tepe, Ali Kosh, Chagha Sefid (Iran), and Khiro-Kitiral (Cyprus) (Niknami et al., 2011). Some well-known Chalcolithic sites at which modified crania have been discovered are She Gabi (Iran), Eridu (Iraq), and Byblos (Lebanon). Furthermore, some of the Bronze Age sites of the Near East have also yielded evidence of modified crania, the most important of which are: Enkomi (Cyprus), Hayas Hoyuk, Adiyaman (Turkey), Lachish (Palestine) (Niknami et al., 2011). The tradition of cranial modification is also found in southern Turkmenistan, Chorasmia, and Tajikistan, extending to the Caspian and Black Seas. Some are later than the samples of the Near Eastern regions and have been dated between the Bronze and the Later Iron Ages. In the Near East, cranial modification has probably continued until recently. Some tribes of Caspian and Black Sea coastal areas still practice it today (Niknami et al., 2011).

From the human skeletal remains from Gol-Afshan, a petrous bone was sampled by Dr. Maryam Ramezani and was sent to the HUN-REN Research Centre for the Humanities, Budapest, Hungary for genetic analyses (IRN24).

#### Chapter 2: Shahr-i Sokhta cemetery

This section has been drafted by Dr. Hossein Moradi.

##### 2.1. Geographical information

Shahr-i Sokhta, 57 km south of Zabol, is located in the Southeastern region of Iran in the southern part of the Helmand River in the Sistan and Baluchistan province. It is one of the most important archaeological sites in Western Asia in the Late Chalcolithic and Early Bronze Age. During the 3rd millennium BCE, Hamoon Lake had surrounded the west and north of the site and was the main source of the drinking and agricultural water for more than a millennium. The first excavations were conducted by an Italian mission between 1968-1978 (Amiet et al., 1978) and after a 20-year gap, were started again in 1997 by Iranian archaeologists from ICAR, excavating the graveyard, the urban centre and industrial zone (Sajjadi & Moradi, 2014).

##### 2.2. Periods and cultures:

The cultural sequence of Shahr-i Sokhta is divided into four main periods and recently to eleven construction phases between 3550 to 2300 BCE (Sajjadi & Moradi, 2024). In the middle of the fourth millennium BCE, the first half of the Period I (about 3550 BCE) the little village of Shahr-i Sokhta was formed on one of the biggest *Kaluts* in Sistan plain with two big depressions that were used for storing water. There is no evidence for human burials in the floor of the buildings. Due to increasing population in the site at the second half of the 4th millennium BCE (Period I phase 10-9, around 3300-3000 BCE) the area of the village covered the region about five hectares and the necropolis in the south was used for burial ceremonies. In this period the small village of Shahr-i Sokhta transformed into a small town. The people who settled at Shahr-i Sokhta in the late 4th millennium BCE came from different regions, mainly from Kerman area of Iran, southern Turkmenistan, and Quetta valley in the centre and north of Pakistan-Baluchistan. The cultural materials such as pottery, figurines, and exchange objects—including seals impressions, tokens, and tablets—demonstrate the presence of different cultures from east and west (Sajjadi & Moradi, 2014). The Nal and Quetta wares from central and north of Baluchistan, the Emir gray were from central Baluchistan (Probably from Bampur) and Makran in Pakistan, the buff wares with solid geometric designs from Namazga III in southern Turkmenistan, and finally the red wares and four lung jars—referred to as pseudo-Jemdet Nasr-type—as well as Proto Elamite tablets and cylinder seal impressions mainly from Kerman area, indicate the development of trans-regional relationships at Shahr-i Sokhta between 3300 to 3000-2900 BCE, covering the entire Period I. This period is contemporary with the Proto Elamite period in the western areas such as Kerman, Fars and Susa plains (Moradi, 2023).

In the beginning of the next millennia, during Period II starting around 2800 BC, the first remarkable development was held in urban constructions as well as socio-economical aspects. During these times the urban centre expanded to 30 hectares both in the Eastern Residential Area and Central parts of the site. Furthermore, some small sites (less than 1 hectare) scattered around the urban centre in a radius of five kilometres (Moradi & Rajabioun, 2022) appeared, formed through interactions with the main core of settlement in Sistan plain. In this period (2800-2500 BCE) the standardisation of the art style of Shahr-i Sokhta was fixed and the city built up a unique ethnicity by mixing different styles and making their own cultural materials. The black on buff wares with linear motifs became the most popular pottery design among the other kinds of the wares. Although the grey and red wares were used both in ritual ceremony (burial costumes) and domestic life. A lot of grey and red pottery were discovered both in the necropolis and residential areas of Shahr-i Sokhta. The abundant usage of the seals witnessed not only in the site but also in the satellite villages around the city, suggests the interaction between the centre and the periphery in the 30 kilometre radius and beyond. Moreover, from the graveyard, the stamp seals came only from the female graves. In the middle of this period (Phase 5 around 2600-2500 BCE) the area of the urban centre reached to about 120 hectares and the spatial separation or specialisation became evident in architectural styles and urban developments in such a way that the big buildings were built on the northern parts of Shahr-i Sokhta and the craftsmen workshops relocated to the western edge of the mound and also to numerous industrial sites in the south. The site was separated by different areas—primarily residential and monumental areas, industrial zones, and the graveyard—based on the socio-economic functions (Mariani, 1992).

In Period III (2500-2350 BCE) with the development of the urban centres all around the southeast of Iran, the area of Shahr-i Sokhta expanded to 200 hectares, reaching its peak, becoming one of the largest sites in the entire Iranian Plateau. Period III is the continuation of the previous period in art style and architecture with some changes in the details. However, some modifications in architectural techniques are evident, including the application of thick plaster on the walls and the presence of public structures (Moradi, 2020). For example, the building no. 20 as a temple or the building no. 26 as a bazaar were built for public use (Sajjadi & Moradi, 2014). Such examples, not only in the urban area but also in the cemetery of Shahr-i Sokhta suggest some changes in the lifestyle of the people. With the amplification of transregional relationships, possible accumulation of wealth and forming an elite class, Shahr-i Sokhta entered the new era of development. More evidence for such changes came from the excavation at the graveyard. Due to the increasing number of catacomb graves (Moradi & Rajabioun, 2022) as well as the high number of goods in such kinds of graves (up to 226 goods) we can infer that the catacomb graves are connected to the khanates or elite classification (Sajjadi, 2015). Existence of such a class is reflected not only in the structure of the graves but also, in the urban area with evidence of large buildings with many storage rooms such as building no. 1(Sajjadi & Moradi, 2014).

In comparison with previous periods, few evidences of Period IV (2350-2000 BCE) were collected only from the main areas of Shahr-i Sokhta and the majority are obtained from the various sites around Shahr-i Sokhta, with more in the Rud-i Biaban area in a radius between

5 to 50 kilometres in the south and southeast region. Due to the majority of the evidence being concentrated near the branches of Rud-i Biaban, a seasonal river that flowed in the southern area of Shahr-i Sokhta, we named the Rud-i Biaban Phase using a kind of material dated back to the last centuries of 3$^{rd}$ millennium BCE (Moradi & Rajabioun, 2022). The materials here are in contrast with a series of sites around Shahr-i Sokhta, such as at Tepe Taleb Khan, Tepe Gratziani, Tepe Pir Zal. The pottery of this phase at Shahr-i Sokhta, was collected only from the upper layers and marginal structures from Burnt Building, building no.26 and building no.20 (Moradi & Rajabioun, 2022). Graves which are connected to this phase are rare at the necropolis and all are related to the earliest part of this phase (2300 BCE or maybe earlier). The similarities between the pottery of the Rud-i Biaban Phase and some BMAC[1] wares in the early phase are questionable and encourage us to mention that the Rud-i Biaban Phase has a great role in the formation of BMAC culture in the northern parts of Sistan (Moradi, 2023). Moreover, evidence for the relationship with southern Turkmenistan came from the catacomb and memorial graves (Sajjadi, 2015), a kind of grave without any burial remains but with the goods. In contrast to Period I (early phase) when the evidence directly came from southern Turkmenistan, at the end of Shahr-i Sokhta sequence, some movements in the Khorasan, Afghanistan, and Turkmenistan area should have originated from the Sistan area. Moreover in this period, the cultural relationship with the Indus valley is ambiguous because of lack of evidence of Harappan material culture in Shahr-i Sokhta (Jarrige et al., 2011). Some scholars believe that when the Indus civilization arose in the middle of the third millennium B.C, Shahr-i Sokhta was under the pressure of some social and environmental crisis that led to the end of occupation of the site in the 2400~2300 B.C. They mention that there are no obvious evidences of Harappan culture among the Shahr-i Sokhta cultural materials for the last Phase i.e. Period IV (Jarrige et.al 2011). Based on the vast excavation of the necropolis, it has been estimated that the area of the graveyard of Shahr-i Sokhta covers a region about 25 hectares in the south side of the site with more than 40,000 graves and 60,000 skeletons (Sajjadi, 2015). It is worth noting that only 1,100 graves have been revealed up to now from the 5,500 m2 excavation held by Italian and Iranian missions (Sajjadi, 2023).The graves include 10 types of structures in which simple, bipartite and catacomb graves are more common than the others. The graves have zero, one or more burials, sometimes also including animal remains, such as goats and dogs. In addition to a *Macaca fuscata* skeleton recently excavated in a simple grave with an unpainted pear beaker shaped near the body (Minniti & Sajjadi, 2019)

Sixteen percent of the total number of graves, or 160 graves in total, were found without any grave goods, likely due to the low social class of the burials or the young age of the deceased. In the others, there were between one and 226 goods in each, including pottery, bronze objects, stone beads, seals, bones and clay objects. The catacomb graves represent the richest on the site. In addition, there are some graves without any skeletal remains but containing some goods. The reason for the construction of such empty graves remains

[1] BMAC stands for Bactriana Margiana Archaeological Complex

unclear and cannot be firmly attributed to the commemoration of lost individuals. This type of grave is also related to the BMAC funerary culture in the past millennia between 2300-1500 BCE (Moradi & Rajabioun, 2022).

**2.3. Information on the skeletal samples**

The main information of this section was collected by the excavators of Shahr-i Sokhta, Dr. Seyyed Mansur Seyed Sajjadi and Dr. Hossein Moradi.

The four human skeletal samples, dating back to Periods I and II and unearthed over the course of three excavation seasons (1997, 2002 and 2003) led by Dr. Seyyed Mansour Seyyed Sajjadi, were analysed for this study.

A brief summary of the information related to these graves is provided below:

1. The excavations of 1997: First Season:

The first of five excavation seasons at the Shahr-i Sokhta cemetery, spanning from autumn 1997 to 2001, involved the exploration of nineteen 10 x 10 metre trenches, cumulatively covering 880 square metres. The initial season alone led to the discovery of 137 burials. One notable grave, number 1520 in trench IUL, situated in a high-density area of the cemetery, is characterised by its flat, pebbled surface. This 10x10 metre trench yielded 22 skeletons, with graves varying from two-part pits and simple pits to crypt-like structures and a unique circular grave with a sealed entrance.

2. The excavations of 2002: Sixth Season:

In 2002, during the sixth excavation season, 12 trenches encompassing 729 square metres were explored, revealing 78 graves. The majority were either simple or two-section graves, totalling 38 and 39 respectively. A single crypt-like grave was also identified. From these 78 graves, 84 skeletons were excavated, with three graves notably lacking skeletal remains.

3. The excavations of 2003: Seventh Season:

In 2003, excavations were carried out in 11 trenches, covering a total area of 526 square metres, leading to the discovery of 66 graves. These included 21 simple pit burials, 43 two-section burials and 2 crypt-like burials. Notably, 8 of these burials yielded both human and animal remains. A total of 72 human skeletons were unearthed and classified as follows: 23 children, 21 women and 15 men. The sex of 13 skeletons remained undetermined.

**2.4. Graves analysed in this study**

1. Trench IUL, Grave 1520 (1997): An irregular polygonal, two-section pit. Artefacts and a skeleton were placed in the northern section, with the skeleton buried supine, facing northeast. The individual (IRN44), estimated to be a female aged 35-40, was discovered with her hands on her chest. The dating of this grave, based on the archaeological context, is estimated to 3350–2900 BCE, corresponding to Period I.

2. Trench MIC, Grave 4800 (2002): This grave is a two-section type, located at a depth of 100 centimetres within the trench and situated beneath a sandy layer. It contained a human skeleton (IRN30) positioned in a curved form, lying on its left side in a north-south orientation. Due to the significant deterioration of the bone remains, it was not possible to ascertain the sex and age of the individual with certainty. A marble bowl was also found beside the skeleton, enhancing the grave's archaeological significance. The dating of this grave, based on the archaeological context, is estimated to 3350-2900 BCE, corresponding to the first occupation period.

3. Trench IPV, Grave 3402 (2002): A two-part burial adjacent to the western trench wall, containing a female skeleton (IRN27) estimated to be 30-35 years old, buried in a curved position on the left side. Seven grave goods were found, including pottery bowls and a stone bead. The dating of this grave, based on the archaeological context, is estimated to 2800-2600 BCE, corresponding to the second occupation period.

4. Trench HYM, Grave 5100 (2003): A simple pit grave housing a male skeleton (IRN43) estimated to be 30-35 years old, buried in a right-side, north-south orientation, with the face almost looking upwards. The dating of this grave, based on the archaeological context, is estimated to 2800-2600 BCE, corresponding to the second period of the site.

#### Chapter 3: Gohar Tepe

This section was drafted by Motahareh Amjadi and Ms. Arezoo Bibak and Mr. Serollah Ghasemi Gorji. The anthropological studies were conducted by Dr. Maryam Ramezani and Ms. Zeinab Salehi.

##### 3.1. Site description

Gohar Tepe, a significant prehistoric archaeological site in northern Iran, is located in Mazandaran Province, Behshahr County. This site, about 2 kilometres northwest of Rostamkola and 30 kilometres along the Sari to Behshahr road, covers approximately fifty hectares and consists of a series of interconnected hillocks. The highest point, at coordinates 36.40814°E and 53.24082°N, stands 32 metres above sea level. Located roughly 800 metres south of the central Alborz Mountains' foothills and 20 kilometres from the Caspian Sea to the north, Gohar Tepe's strategic position provided easy access to a wealth of natural resources.

This ancient site, which served both residential and burial purposes, was utilised from the Chalcolithic to Iron Age. From 2003 to 2012, Dr Ali Mahforoozi and his team led approximately ten seasons of archaeological excavations at Gohar Tepe. These excavations uncovered significant findings from the Bronze and Iron Ages, including skeletal remains, 226 human burials and two animal burials (Ghasemi Gorji, 2017).

Of these discoveries, four human skeletal samples were specifically selected for our research.

A summary of the graves associated with these remains is provided in the following section.

##### 3.2. Human skeletal samples

1.Trench AJ2XX, burial 2:

This context includes a human burial, positioned in a contracted position on the right within the burial pit. The burial is oriented from north (head) to south (feet), with the face directed eastwards. To the west of the skeleton, seven varied ceramic vessels were found, including a pitcher, a cup, a bowl and a teapot (characterised by a spout or beak-like pourer). This burial (IRN71) was identified as a male aged between 40-50 years and excavated during the archaeological projects conducted in 2007 and 2008. The dating of this grave (Supplementary Figure 3.1), based on the archaeological context, is between 3100 to 2300 BCE (Piller & Mahfroozi, 2009).

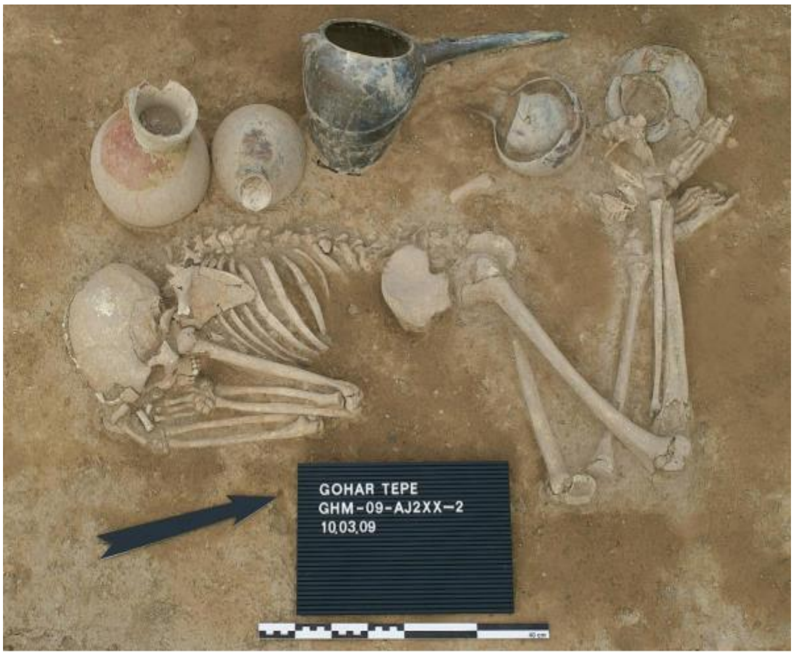


**Figure 3.1. The structure of the grave, the condition of the burial, and the placement of the grave goods within T-AJ2XX-2 (Piller & Mahfroozi, 2009).**

2. Tench AJ2XX, burial 8:

This context involves a human burial placed in a foetal position on its left shoulder within the burial pit. The burial's orientation is from east (head) to west (feet), with the face turned southward. Near the forehead area (south), a relatively small grey ceramic vessel was discovered. This burial (IRN72) was identified as probably male between 25-30 years of age and excavated as part of the archaeological projects conducted in 2007 and 2008. The dating of this grave (Supplementary Figure 3.2), based on the archaeological context, is between 3100 to 2300 BCE (Piller & Mahfroozi, 2009).

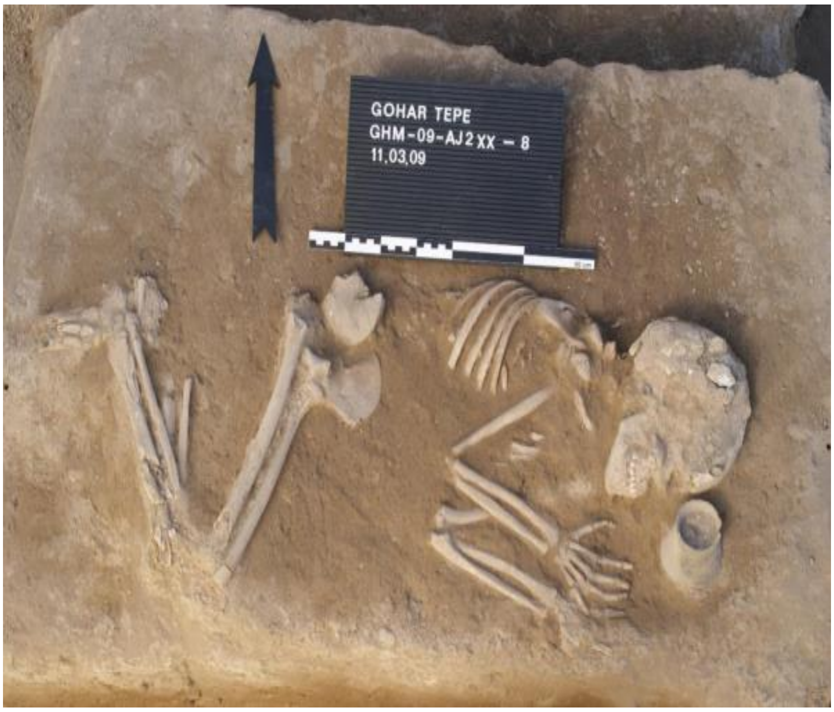


**Figure 3.2. The structure of the grave, the condition of the burial, and the placement of the grave goods within T-AJ2XX-8 (Piller & Mahfroozi, 2009).**

3. Trench AG2IV, burial 96:

This context involves a human burial situated in a foetal position, resting on the left shoulder within the burial pit. The burial is oriented northwest (head) to southeast (feet), with the face directed east. Around the head area, three ceramic vessels were found and notably around the waist and near the hands, a considerable number of decorative items made of bronze were discovered. This burial (IRN70) was identified as probably-male aged between 25-30 years and excavated as part of an archaeological project in the year 2008. This grave, based on the archaeological context, is dated between 2600 to 2300 BCE (Supplementary Figure 3.3) (Piller & Mahfroozi, 2009).

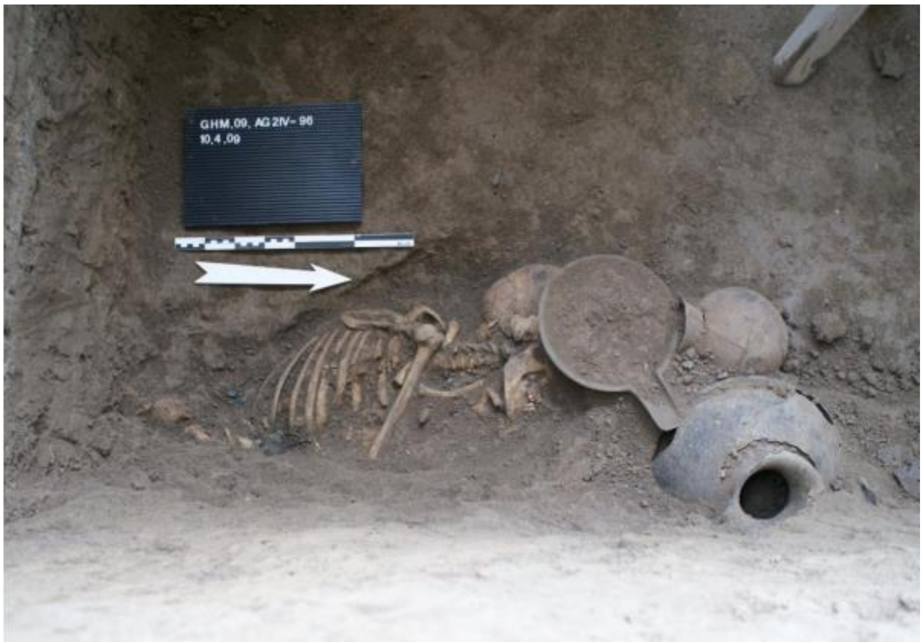


**Figure 3.3. The condition of the burial and the placement of the grave goods within T- AG2IV -96 (Piller & Mahfroozi, 2009).**

4. Trench AI2XX, burial 3:

This burial ranks as one of the most significant and richly appointed discoveries at Gohar Tepe.

The burial was oriented southwest to northeast and positioned in a foetal position on the left shoulder. Due to the skeleton's poor state of preservation, sex determination was not possible. From this burial, 11 pottery vessels of various forms and a stamp seal with a geometric design were recovered. The pottery collection features a spectrum of colours, including light and dark grey, red and brown. This burial (IRN80) was identified and excavated during archaeological projects in the years 2006 and 2007. The direct $^{14}$C radiocarbon date (1625-1505 cal BCE) (Supplementary Table S1) confirms the dating of the IRN80 (Supplementary Figure 3.4) to the Iron Age (Ghasemi Gorji, 2017).

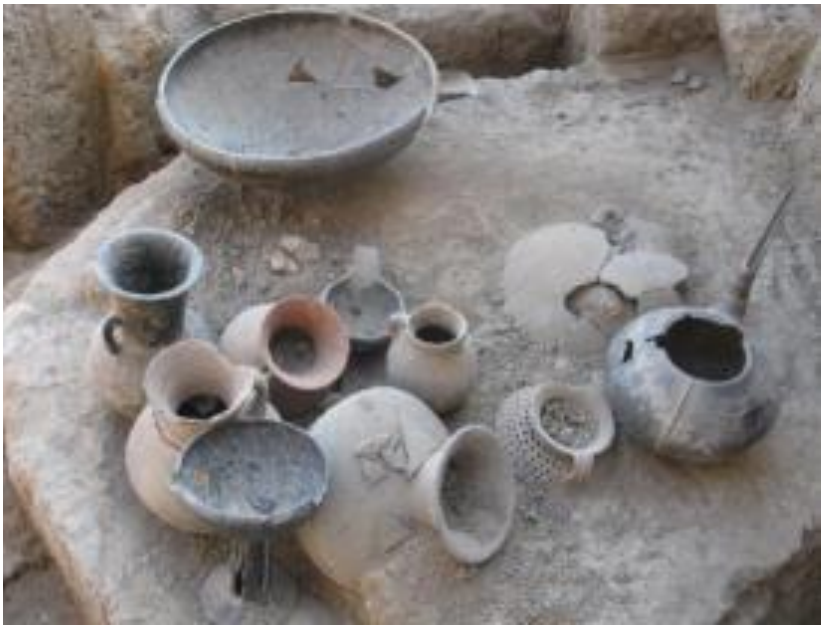

**Figure 3.4. The placement of the grave goods within T- AI2XX (Ghasemi Gorji, 2017).**

#### Chapter 4: Cham Papi Cemetery

This section was drafted by Motahareh Amjadi and Dr. Lili Niakan. The anthropological studies and sampling for this research was conducted by Dr. Maryam Ramezani.

##### 4.1. Cemetery description

Located on the bank of the Rue River within the Kaferi Strait, the Cham Papi Cemetery covers an area of one hectare and is elevated 12 metres above the surrounding land. The Rue River, originating from the Kolm and Jaber Rivers' confluence, enhances the site's geographical significance. Positioned at coordinates 33.2235 °N latitude and 46.5947 °E longitude and with an altitude ranging between 630 to 730 metres above sea level, the cemetery is set upon a geologically significant cliff from the third geological period. This ancient site, lying in the western expanse of the Kaferi Strait and at a distance of 3.5 kilometres from the primary Ilam to Badreh thoroughfare, indicates that the site dates back to the Middle Bronze Age through surface discoveries, including pottery fragments and remnants of stone wall architecture (Niakan, 2018).

The cemetery harbours a rich array of historical remnants, with tombs and graves dating back from the third millennium BCE to the early first millennium BCE. Dr. Lili Niakan and her team observed extensive damage across various site areas, resulting from unauthorised excavations and activities by cultural profiteers and looters. The site reveals a mix of tombs and individual graves, constructed at differing levels on a rocky slope, seamlessly integrated into the natural mountain bed.

The preservation of the skeletal remains has been significantly impacted by environmental factors, including rainfall and water seepage, which have led to the deterioration of most skeletons and have obscured their original burial positions. Notably, many graves, in keeping with an ancient tradition of offering food to the deceased, contained remnants of vessels

with animal meat, wheat, lentils, oak seeds, and wild pistachios. Furthermore, the graves also contained burial gifts such as pottery, stone objects, and bronze tools (Niakan, 2018).
In this study, human skeletal remains from a grave in which two individuals were buried were examined.

**4.2. Grave's information**

1. Trench C, Grave 24:

This square-shaped grave, located in the site's eastern part on a north-south oriented rocky slope, measures 70 centimetres in length, 70 centimetres in width and 137 centimetres in height, was built atop a surface elevated 158 centimetres above ground level. The interred body was positioned in a squatting pose, with the grave likely serving as a secondary burial. The bones, especially the skull and face, were found in a completely fragmented state, and the skeleton was oriented southwards. Beside the skeleton lay a ceramic bowl and a metal pin. The grave's walls, composed of river and nearby cliff stones, were structured with two layers of stones and a mud mortar layer. Preliminary anthropological analyses suggested that the remains in this grave likely belonged to two individuals. The grave goods, including pottery vessels, resemble pottery types discovered at Tepe Guran and Godin Tepe, dating back to 1800-1600 BCE (Niakan, 2018).
Among the skeletal remains, the first individual belonged to an adult, likely female, as indicated by the shape of the mastoid process. Dental examination suggested the individual (IRN29) was a young woman aged between 18-20 years. Next to the skull, a conical-shaped pin with a narrow tip and a decorated round bottom featuring small circular patterns was found. Limb erosion precluded height estimation, but dental examination revealed most teeth and sockets were intact, with mineral coverage more intense on deciduous teeth.
The second skeletal remains (IRN33) in the same grave belonged to a child. Due to unclear dental conditions and incomplete limb development, estimating the child's age was not possible (Niakan, 2018).

#### Chapter 5: Kalmakareh cave

This section was drafted by Ms. Arezoo Bibak and Motahareh Amjadi.

##### 5.1. The geographical area of Kalmakareh Cave

Kalmakareh Cave is situated in Kohdasht County in the southwestern part of Lorestan Province, 98 kilometres northwest of the central region of Pol-e-Dokhtarand is perched at an altitude of 448 metres above the plains. The cave is positioned on the eastern side of one of the final rocky valleys of the limestone Malleh Mountain. In terms of geographic coordinates, the cave is located at 33.13315°N latitude and 36.47380°E longitude (Khosravi, 2013), with an elevation of 1,270 metres above sea level. Mal-e Mountain, a significant limestone mountain in the western Lorestan Province, begins to the west of Pol-e-Dokhtar City and rises to a height of 2004 metres. This mountain, known for its numerous caves and crevices, is covered with vegetation, including oak trees, wild almonds, figs, and pistachio trees (Motamedi, 1993). The unique natural characteristics of Kalmakareh Cave have always made access to it quite challenging.

##### 5.2. The location of Kalmakareh Cave

The Kalmakareh Cave is located on the eastern side of Malleh Mountain, along the smooth and flat walls that stretch from the summit to the base of a valley, at the junction where the valley walls converge on the northern side of the mountain. The entrance to the cave is situated thirty metres below the mountain's peak, with the front of the cave protruding one and a half metres from the lower section of the entrance. The front area of the cave has been segmented into three entrances due to the presence of several large rocks (Motamedi, 1993).

The entrance of the Kalmakareh Cave faces westward and penetrates eastward into the mountain. The cumulative area of the spaces mapped out within the cave amounts to roughly 4300 square metres, with the combined length of the interconnected chambers along the present route totalling approximately 670 m. Not only is the journey to the cave's entrance arduous, but navigating through the cave's interior proves equally demanding. The cave encompasses four large chambers and the corridors that connect these chambers, as well as some of the cave's entrances, vary in diameter from a narrow 80 centimetres to a spacious 15 m (Ghazanfari, 1987).

In his studies conducted in 1993, Moatamedi highlighted the environmental conditions of the region surrounding the Kalmakareh Cave. He pointed out the scarcity of water and springs in the eastern parts of Malleh Mountain, noting their unsuitability for drinking or agricultural use. He also mentioned the lack of accessible food sources in the vicinity of the cave. Moatamedi's observations include the presence of water throughout the length of the cave, high heat and humidity during the warmer seasons, and the enclosed nature of the cave's interior. He noted that the cave's internal air circulation is facilitated only by night-time storms.

Considering the absolute darkness, that dominates the cave and the existing climatic conditions, Moatamedi deemed the possibility of maintaining a long-lasting fire for lighting, warmth during cold seasons, and cooking, as almost impossible within the Kalmakareh Cave. He interpreted these factors as indicators of the impracticality of sustained human habitation in the cave. Furthermore, Moatamedi cites the absence of hearths and thin pottery vessels for daily use as additional major signs of the lack of prolonged human presence in the cave (Motamedi, 1993).

These observations collectively suggest that the cave was not a site of continuous human settlement, but rather, it may have been used occasionally or for specific purposes.

**5.3. Research history of Kalmakareh Cave**

In December 1989, a local hunter accidentally discovered the treasure of the Kalmakareh Cave. This chance discovery, likely due to the collapse of walls that had concealed the treasure from sight, became the hunter's fortune. Unfortunately, before the cultural heritage authorities of Lorestan Province in Iran could be informed, looters plundered a significant portion of this treasure.

Hossein Ghazanfari's report in the autumn of 1987 on the Kalmakareh Cave, published in the journal Asar, was the first and most significant report concerning the geographical location and description of the cave's interior in Iranian publications (Kalhor, 2018). Following this, Mir Abedin Kaboli in his article on the study of pre-Islamic artefacts, published in the Cultural Heritage journal in the autumn of 1991, provided an analysis and description of some of the artefacts from this cave (Kalhor, 2018).

Ahmad Parviz, in a paper presented at the International Conference on Iranian Archaeology held in Kermanshah Province in 2006, focused on the gilded silver vessels from Kalmakareh (Kalhor, 2018). Eventually, an archaeological team was dispatched to Lorestan in July 1992 to conduct excavations and studies (Motamedi, 1993).

Further excavation in the Kalmakareh Cave proved futile, as no remnants of the treasure remained, with a substantial portion of the artefacts having been smuggled out of Iran over the course of four years. Most of these artefacts were trafficked to neighbouring countries. Dondaz published information about the artefacts smuggled to Turkey, yet several others remain in the country with no available information. The first news of Kalmakareh artefacts outside Iran emerged in 1993 in the Independent and other Western newspapers, and even in media outlets, under the title "The Western Cave Treasure." The renowned antiquities dealer Houshang Mahboubian in London purchased a significant portion of the Kalmakareh artefacts. The inscriptions on these objects were deciphered by Lambert, identifying 22 Elamite names. François Vallat also conducted a study and research on the inscription of a silver rhyton (Vallat, 2017). The inscriptions on the artefacts still in Iran were read and published by Rasoul Bashash (Khosravi, 2013).

Today, approximately 90 artefacts from the discoveries in the Kalmakareh Cave are preserved in Iran. In 1989, Leila Khosravi conducted a chronological study of these items in her master's thesis at the University of Tehran (Pirmomen, 2013). Ultimately, in her doctoral dissertation, with the research objective of proving the artefacts' authenticity and their association with the culture of ancient Iran, she conducted a comprehensive study on their technique, decorations, and the influence of the Kalmakareh Cave artefacts on the metalworking art of subsequent periods. Following this, the osteological data obtained from the first hall of the cave were examined by Venus Pirmomen in her master's thesis at the University of Science and Research. This scholarly work contributed significantly to the understanding and appreciation of the historical and cultural value of the artefacts and remains from the Kalmakareh Cave (Pirmomen, 2013).

The artefacts attributed to the Kalmakareh Cave have been categorised into groups: 1) Plain and undecorated vessels, 2) Cups and saucers, 3) Goblets and drinking vessels (situlae), 4) Tubular vessels, 5) Amphorae, 6) Rhytons, 7) Human and animal figurines, 8) Zoomorphic statuary vessels, 9) Jewellery, 10) Ceremonial tools, 11) Decorative and symbolic objects. A significant characteristic of the metal artefacts from Kalmakareh is their incorporation of various animal forms in their design (Khosravi, 2013).

##### 5.4. Samaturra or Samati

The Elamite civilization, with its territorial expanse covering modern-day Khuzestan, the back hills of Lorestan and the Bakhtiari mountains, was historically managed in a federal manner including major cities. Independent governments with autonomy emerged around these cities in the 7th century BCE. However, this autonomy weakened as central power strengthened, eventually unifying these independent governments under a single command. One such independent local government, revealed through inscriptions on the artefacts from the Kalmakareh Cave treasure, was the land of Samatura, belonging to the Late Elamite Period and located in southern Lorestan (Khosravi, 2010). The names of five generations of rulers from this realm were identified on these objects, with the majority of the names belonging to the second generation of this dynasty, namely Ampirish. The presence of inscriptions in Late Elamite script on the objects and the use of names with Elamite linguistic roots clearly indicate a connection to Elamite culture. Furthermore, names with Indo-Iranian roots suggest a fusion of traditions between these cultures. Apart from two inscriptions in Aramaic and Neo-Assyrian script, the rest are inscribed in Late Elamite script.

The inscriptions on the artefacts from the Kalmakareh Cave treasure, purchased by the renowned collector Houshang Mahboubian, were studied and deciphered by François Vallat (Vallat, 2017). Based on Lambert's research, Vallat acknowledged that among the 22 Elamite names identified by Lambert, only four were titled as Lugal (rulers) of Samaturra, which are as follows (Vallat, 2017):

Ampiriš

Anni- šilhak

Unzi- kilik

Unsak

**5.5. The Inscriptions of the Kalmakareh Cave and their Interpretations**

Out of the 90 artefacts found in Iran from Kalmakareh Cave, six have inscriptions. Four of these have been deciphered by Rasoul Bashash, as described below:

1. On a silver rhyton, which is supported by three horned goats at its base (Supplementary Figure 5.1), there is an inscription in Aramaic script and language dating back to the 7th century BCE. The inscription reads:

"The cup of Kuban's governor[2] with the goat vessel for the health of Madyes king[3] or "The cup of Kuban's governor Azlerspand (Astiyag)[4] for the health of Madyes king (Bashash Kanzaq, 1999).

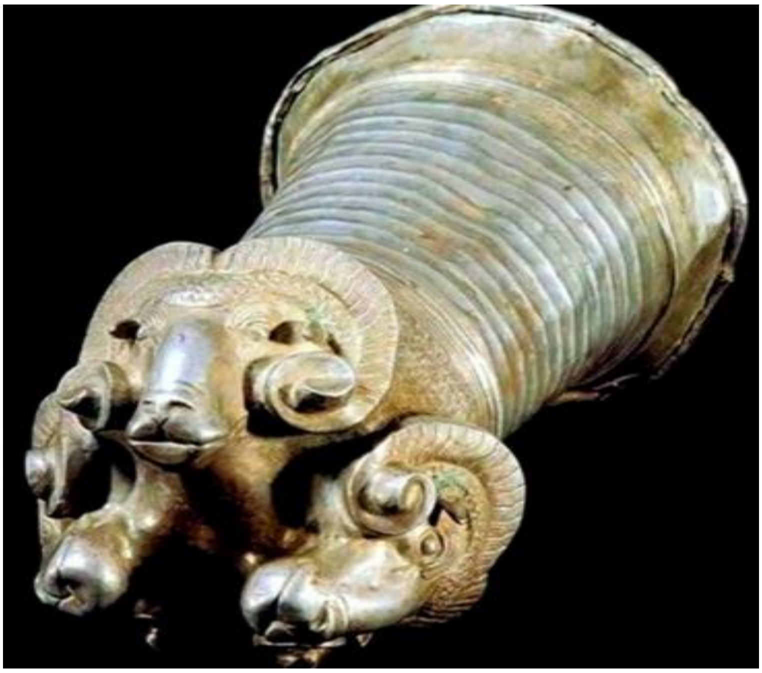

**Fig 5.1. Rhyton silver of Kalmakareh (Khosravi, 2010)**

[2]The Kuban region is a historical and geographical area located in the southern part of Russia, within the North Caucasus.

[3]During the 7th century BCE, Madyes, a monarch of the Scythians, governed in the period of their expansion into Western Asia.

[4]The Last king of Medes Empire

2. A silver-handled vessel shaped like a bucket, on which there is a cuneiform inscription in the Assyrian language, written across two lines:

Offered to the god 'ADAD'[5], the god of wind and storm, united with the city of Gilzanu

Asarhaddon, King of Assyria, son of Senakhrib king of the land of Assyria, may he be the one to judge (Supplementary Figure 5.2) (Khosravi, 2010).

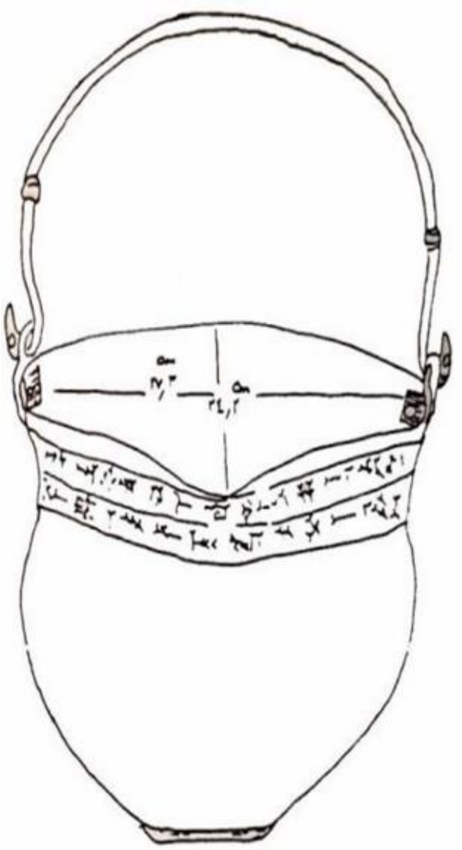

**Fig 5.2. Silver-handled bucket (Khosravi, 2010)**

3. Other inscriptions, both engraved on a cup and its silver coaster from Kalmakareh. These inscriptions share a common theme, which is why they are presented together here. The script on both items is cuneiform and the orthography aligns with Neo-Elamite, a style still in use during the Achaemenid period. The inscriptions on these vessels read as follows:

“Amprish the king of Samaturra/Samataoro, The son of the Tabal”

[5]The god ADAD is a prominent deity among the Arameans and the Hittites, known as the great god of weather. The most famous temples dedicated to this weather god are located in Northern Syria and the Taurus region. The city of GILZANU is also one of the well-known cities from the period of the Mannaeans (Khosravi, 2010).

Samaturra/Samataoro was the centre of governance for the rulers of the Samatu (the owners of the Kalmakareh artefacts). The exact location of this region remains unidentified to date. One hypothesis is that the Samaturra were actually the Cimmerians (Khosravi, 2010).

##### 5.6. Human remains from Kalmakareh Cave

The osteological materials of this cave were analysed by Venus Pirmomen (Pirmomen, 2013).

<u>1. IRN85:</u>

The skull of a 15 to 17-year-old female (genetically and anthropologically), which had the highest number of teeth among these three skeletons, with 6 teeth in the maxilla. Traces of fractures are visible on the cranium, which are related to environmental pressures after death[31].

<u>2. IRN84:</u>

Belonging to a mature male, aged between 49 to 51 years, this skull has only two molars remaining in the maxilla. It was determined that this individual suffered from acute cerebral meningitis during his lifetime and passed away due to this condition (Pirmomen, 2013).

<u>3. IRN83:</u>

This is the complete skeleton of a young male aged between 24 to 27 years. Measurements taken indicate that he was healthy and robust. It is the only complete skeleton remaining from Kalmakareh Cave. The cranial fracture occurred post-mortem due to environmental pressures. From this skull, one upper lateral incisor and one lower first molar remained (Pirmomen, 2013).

##### 5.7. The dating results of Kalmakareh cave

For radiocarbon dating on the bone remains from Kalmakareh Cave, Arezoo Bibak, after obtaining the necessary permissions from Dr J. Nokandeh, the Director-General of the National Museum of Iran, sampled the remains in the Falak-ol-Aflak Museum's repository in Lorestan Province. The radiocarbon dating was performed in the Debrecen HUN-REN ATOMKI laboratory in Hungary. The calibrated radiocarbon date was 1215-1270 cal CE (95.4% CI, Supplementary Table S1) corresponded to the mediaeval period, which does not match the chronology of the Kalmakareh treasures (7$^{th}$ century BCE).

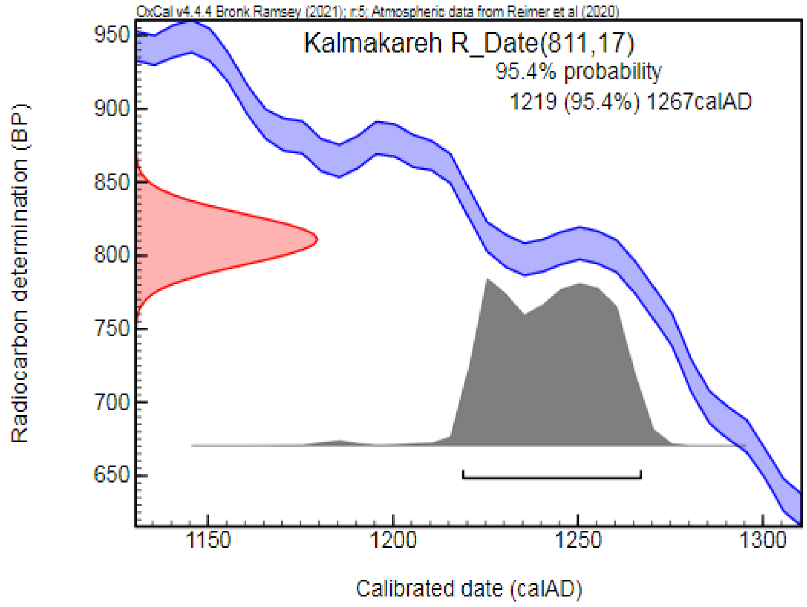


**Figure 5.3. The calibrated radiocarbon curve of the skeleton (IRN85) from Kalmakareh Cave (HUN-REN ATOMKI AMS $^{14}C$ report, Debrecen Hungary)**

##### 5.8. Theories regarding the individuals buried in Kalmakareh Cave

The most prominent and famous theory regarding those human remains from the Kalmakareh Cave suggests that they were the guardians of these ancient artefacts. Other theories suggest they might have been the owners of the treasure or kings/princes of Samaturra or local rulers during the Late Elamite period. A third theory posits that these individuals were neither the guardians of the treasure nor their owners, but they were concealers who had no temporal connection to the treasure (Motamedi, 1993). The dating results given in section 5.7 are completely consistent with the third theory and show that they were concealers of the treasure in the 12th century, that is, a time gap of 1900 years since the Kalmakareh treasures were originally deposited (7$^{th}$ century BCE).

#### <u>Chapter 6: Jubaji site</u>

This section was drafted by Ms. Arezoo Bibak and Motahareh Amjadi.

##### 6.1. Geographical Location of the Jubaji site

The Jubaji archaeological site, located in the Jubaji village, central district, Ramhormoz County, Khuzestan Province, is situated approximately 7 kilometres southeast of Ramhormoz. Its geographical coordinates are 31.1431 °N latitude and 49.4020 °E longitude. The site is at an average elevation of 212 metres above sea level. The Ramhormoz River originates from Mount Lirab, situated 50 kilometres southeast of Izeh. It begins its course under the name of the 'Upper River' towards the village of Chubel. After passing through Ramhormoz and north of the villages of Jubaji and Deh Yor, it joins the Jarahi River and eventually flows into the Arvand Rud (Shihegar, 2014).

In 1948, McCown, on behalf of the Oriental Institute of the University of Chicago, conducted exploratory excavations at the Tel Geser site in the Northwest of Ramhormoz. He uncovered an Elamite fortress dated back to the second millennium BCE, artefacts from the early Elamite period and a cemetery from the first millennium BCE where the artefacts of the New Elamite period (1100-539 BCE) were found. In 2006, Lili Niakan from the Iranian Archaeology Research Institute and Abbas Alizadeh from the Oriental Institute of the University of Chicago surveyed the Ramhormoz plain to understand land use and settlement organisation in the area (Shihegar, 2014).

**6.2. Findings from the Jubaji archaeological site**

**Jubaji Tomb**

In 2007, the exploration in the eastern area of this region began with the discovery of the tomb of two Elamite women from the royal Shutur-Nahunte family. This tomb was hidden within an ancient hill, located one kilometre north of the Jubaji village. The hill, spanning an area of about forty hectares, was excavated by Mrs. Arman Shishegar.

The tomb discovered at this archaeological site is a rectangular stone structure, oriented east-west, measuring 450 centimetres in length and approximately 230 centimetres in width. Due to the frequent floods and surges of the Ala River, as well as the sandy bed and stone structure of this tomb, the extent of damage to the tomb is quite severe—so much so that the entrance of the tomb is no longer recognisable (Shihegar, 2014).

**Bronze Coffins of Jubaji**

In the Jubaji tomb, two bronze coffins were discovered. Based on their positions within the tomb, they have been designated as the 'Eastern Coffin' and the 'Western Coffin'.

The coffins unearthed from Jubaji are unique in their design, resembling bathtubs or having a U-shape and are made of bronze. These coffins were placed parallel to each other but in opposite orientations (the Eastern Coffin facing north and the Western Coffin facing south), with a distance of about one and a half metres between them (Supplementary Figure 6.1) (Shihegar, 2014).

The decorations on these coffins are unparalleled. One of the motifs found is the rosette, or Eastern lotus, which is commonly seen in the civilizations of Mesopotamia and Egypt. Besides Egypt and Mesopotamia, especially during the Assyrian period, this motif was also highly regarded and used in Iran and other Near Eastern countries. For instance, among the collection attributed to the Kalmakareh cave in Lorestan, Iran, which Ms. Khosravi interprets as belonging to the Samaturian or Semite tribes of Indo-Iranian origin; this motif is evident on three circular silver sheets (Khosravi, 2010).

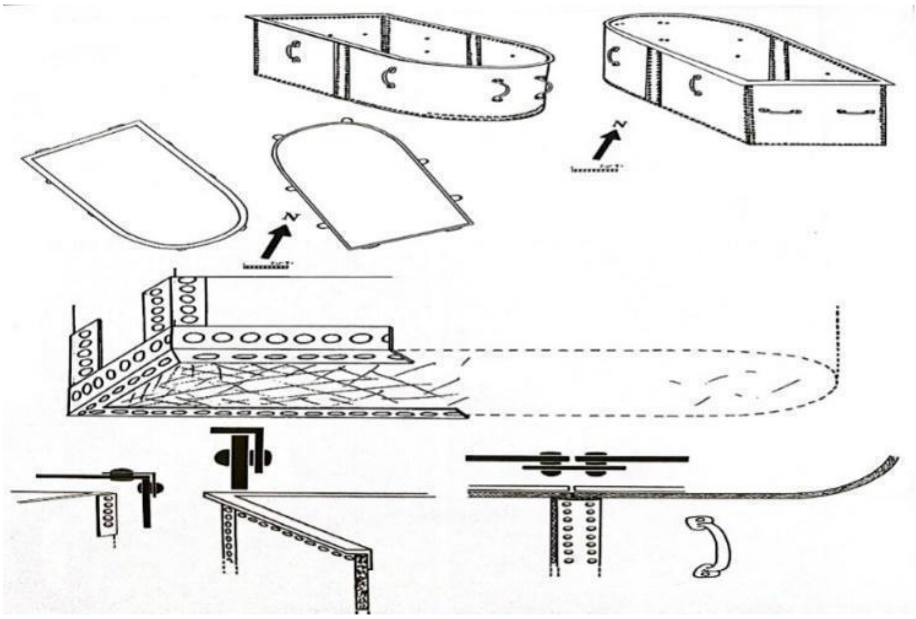

**Figure 6.1. Designs or reconstructions of the coffins, illustrating the method used to rivet the pieces together (Shihegar, 2014).**

This tribe settled around the 6th and 7th centuries BCE in southern Lorestan, near the Seymareh River and founded the local princes of Samatureh (Shihegar, 2014).

#### Metal objects

In the Jubaji tomb, metal objects made of gold, silver, bronze, and iron were found inside the coffins and in their vicinity. The golden objects are the most significant finds, having been placed inside the coffins alongside the buried individuals. Many of the Jubaji artefacts are older than the tomb itself, suggesting a range of different dates. This can be attributed to the fact that the jewellery found in the coffins and the tomb belonged to two Elamite princesses. These jewels, considered royal treasures, were passed down within the family and were ultimately buried with them (Shihegar, 2014).

In the Jubaji treasure trove, a large number of silver objects were found. These include jewellery adorned with precious stones, a number of silver vessels, a bowl, a goblet, a dagger with an inlaid handle, a silver kettle, and a ceremonial vessel shaped like a goddess (Darabi, 2021).

These artefacts are highly comparable to the finds from Haft Tepe related to the Middle Elamite period (around 1500-1300 BCE), as well as those from Susa and Mesopotamia (Supplementary Figure 6.2) (Darabi, 2021).

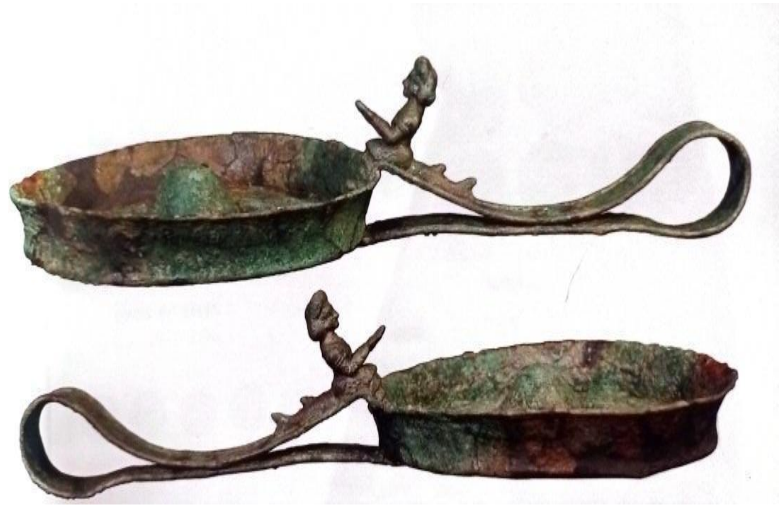

**Figure 6.2. The ceremonial vessel with the figure of a fish Goddess (Shihegar, 2014).**

#### New- Elamite Pottery in the Jubaji site

In the first excavation season, which took place in 2007, a total of 2175 distinctive pots and pottery fragments were found. In the survey conducted to determine the scope and boundaries of the Jubaji-Deh Yur site complex, 293 notable pottery pieces were identified. During the second excavation season in 2014, 500 pottery pieces were collected, of which 115 were categorised and typified (Khosravi, 2010).

The pottery is classified into two groups: glazed and unglazed.

1. Glazed pottery (Supplementary Figure 6.3) includes bowls and jugs, rims and necks of small bottles, bottles with plain bodies, bottles with parallel circular ridges on the body, ornate jugs and bottles, and cubic rectangular vessels with three, two and single compartments, each with a handled lid (Shihegar, 2014).

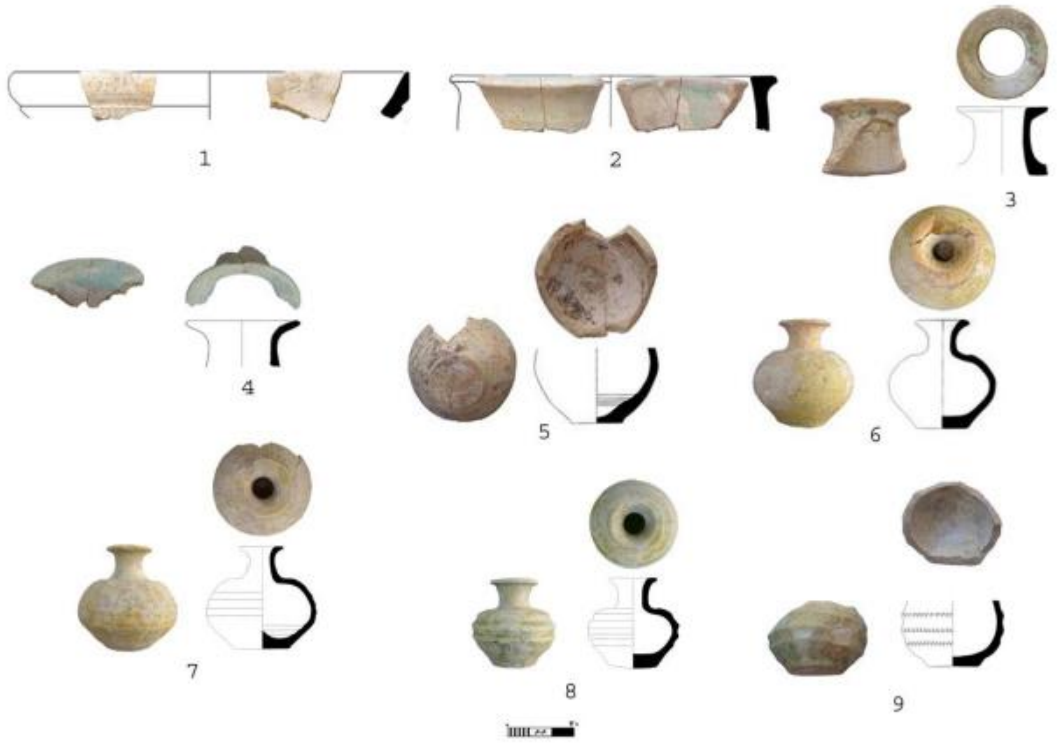


**Figure 6.3. Examples of glazed pottery from the Jubaji site (Shihegar, 2014).**

2. Unglazed pottery (Supplementary Figure 6.4.) includes: shoulder-jar with stepped rims, elongated-bodied jars with rounded base or handle-less amphorae, small elongated-bodied jars with flat bases, small elongated- bodied jars with slightly convex flat base sand cups (Shihegar, 2014).

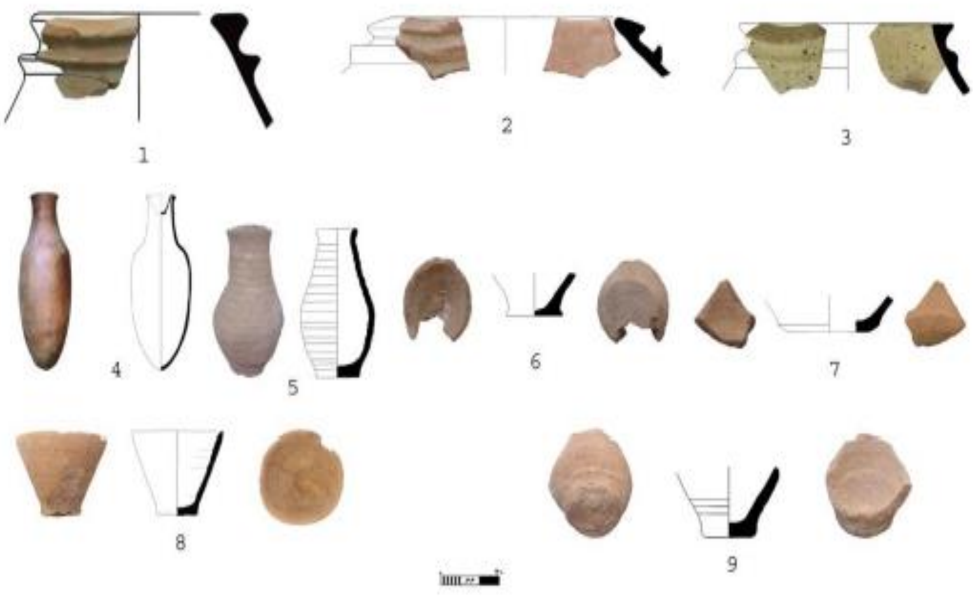


**Figure 6.4. Unglazed pottery from the Jubaji site (Shihegar, 2014).**

#### 6.3. The inscriptions of Jubaji

The inscriptions of Jubaji consist of four tablets, which have been deciphered by Mr. Hamid Rezaei Sadr, a senior expert in ancient languages and ancient texts at the Linguistics Research Centre.

1. The inscription that was found on one of the pages of a golden ring, written in the Elamite language and cuneiform script, has been deciphered by Mr. Hamid Rezaei Sadr. It reads:
"**v.cu-tur $^{d}$.nahunte DUMU in-da-da-na,"** which translates to "Shutur Nahunte[6], son of Indada" [35].

2. On the inner part of a golden armlet, in the Elamite language and cuneiform script, a feminine Elamite name (presumably) is engraved, which reads:
**"La-ar-na**" (Shihegar, 2014).

[6] Shutruk-Nahhunte II, also known as Shutur-Nahhunte, was one of the kings of Neo-Elamite

3. Engraved on the onyx agate seal of Babaghuri[7], belonging to the woman in the eastern coffin, in two lines, each containing two syllables, is the name of an Elamite lady, as follows: **"nu-ma/ku "** (Shihegar, 2014).

4. On both sides of a circular agate stone, which was the gemstone of a golden brooch, two inscriptions in cuneiform script and Sumerian language have been engraved.

Side A: "$^{d}$.en-lil lugal [dingir-mec] "
Lugal-a-ni-ir- v.ku-ir-gal-zu x…zu

Side B: " Ir?-mu-car-tum e?
$^{d}$.en-lil
mu-du-gan?-e
The above statement means that:
Kurigelzu [8]for Soroush (the deity), Enlil[9] King (of the gods)"(Shihegar, 2014).

[7] A type of opal precious stone
[8] Kassities king- 1332 to 1308 BCE
[9] Enlil, Mesopotamian god of the atmosphere and a member of the triad of gods completed by Anu (Sumerian: An) and Ea (Enki)

#### 6.4. Human Remains from the Jubaji site

Collections of osteological samples at the excavation site were meticulously examined and recorded on-site by Mr. Farzad Forouzanfar during the excavation.

Skeletons were found in both the Eastern and Western coffins. In the Eastern Coffin, the remains of a 30-year-old female were discovered, while in the Western Coffin, there were the remains of a 17-year-old female. Out of an estimated total of 60 teeth between these two individuals, 51 teeth were recovered, all of which were heavily damaged (Shihegar, 2014). In total, from 152 potential phalanges, metatarsals, and metacarpals belonging to these two individuals, 70 were retrieved and many others likely perished under the blades of bulldozers. Based on the direct radiocarbon test, the human remains from Jubaji are dated back to ca. 900 to 800 cal BCE (Shihegar, 2014).

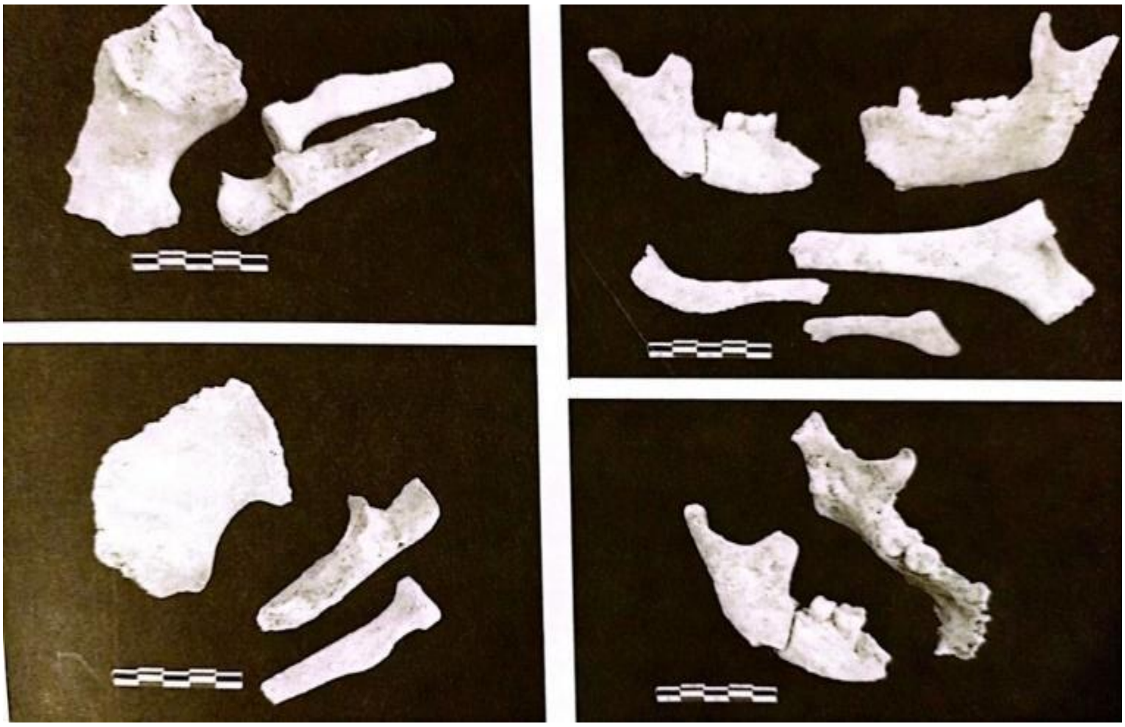

**Figure 6.5 On the right side, remains of the mandible, ulna, clavicle, and the first and second molars from the Eastern Coffin and on the left side, the proximal epiphyses of the radius and ulna and the sciatic notch from the Western Coffin (Shihegar, 2014).**

#### 6.5. Sampling of the human remains

All samples and objects, including archaeological artefacts and skeletal materials from this site, have been transferred to the repository of the National Museum of Iran located in Tehran. Ms. Arezoo Bibak, in coordination with the director of the National Museum of Iran,

Dr. J. Nokandeh, and under the auspices of Dr. Arman Shishegar, the excavator of the site, visited the museum's repository located on Si-e Tir Street and took samples from the two individuals in the Eastern and Western coffins. Considering the remains that were preserved, the sampling was conducted as follows:
Two teeth (IRN81 and IRN82) were taken from the remains of the woman in the Western Coffin and a portion of the distal femur (IRN55) and ulna (IRN56) from the remains of the woman in the Eastern Coffin.
After the sampling, the samples were directly transferred to the HUN-REN Research Centre for the Humanities.

#### Chapter 7: Marsin-Chal cemetery

The information of this chapter was collected by the excavator of Marsin-Chal cemetery, Dr. Ata Hasanpour. Ms. Arezoo Bibak helped to sample the human skeletons, the archaeological materials of this excavation area, and contributed this text part. This section was drafted by Ms. Arezoo Bibak and Motahareh Amjadi.

##### 7.1. Geographical information

The Marsin-Chal cemetery, located within the catchment area of the Finsk Dam in Mehdi Shahr County, Semnan Province, is situated at the geographical coordinates of 36.04578°N latitude and 53.42766°E longitude.

The Marsin-Chal cemetery is approximately 200 metres south of the village of Talajim, located on a natural bed overlooking the Espe-Rud River. The site extends 120 metres in the west-east direction and spans 50 metres in the north-south direction. The site is bordered by Taq Dari mountain and Sertala heights to the south, the Heliashane valley to the west and the site of the Finsk dam to the east. The studies discussed in this section of the article pertain to the third season of excavation at this site.

The third excavation season of the Marsin-Chal cemetery (in 2021), involved two archaeological teams simultaneously excavating an area of 600 square metres. This operation led to the identification and excavation of 130 graves across 6 interconnected trenches. Moreover, a 350 square metre section of the Marsin-Chal cemetery was explored, involving three trenches each measuring 10 x 10 metres and one excavation site measuring 10 x 5 metres. From this area, 73 graves were identified, with 68 of these graves excavated under the supervision of Dr. Ata Hasanpour.

The graves in the Marsin-Chal cemetery consist of hand-dug, rectangular pits, with no evidence of stone lining on the walls, essentially classifying them as pit graves. At the top and bottom of these grave pits, two vertical stones mark the graves. The graves are oriented west-east, with the deceased's head pointing west and their feet east.

Among the graves excavated in this season, only 9 had no grave goods. The rest contained various items made of bronze, iron, decorative beads from precious stones, plaster, and pottery vessels. Generally, the grave goods can be categorised into several types: Out of the 68 excavated graves, 45 contained jewellery. Notably, none of the graves revealed decorations related to horse trappings and tools. The most abundant grave goods were found in Grave 9, which contained 29 items. Furthermore, 33% of the graves in this cemetery contained iron weaponry, but none of these were made of bronze.

Considering the cultural objects and materials retrieved from this cemetery, as well as the results of dating conducted on human samples obtained during the third season of excavation, this cemetery can be dated to the Achaemenid and Parthian periods. In the second season of excavation of this cemetery in August 2014, the information published by Dr. M. Malekzadeh indicated graves pertaining specifically to the Achaemenid period (Malekzadeh et al., 2023).

The skeletal sampling was carried out by Ms. Arezoo Bibak. Anthropological studies were conducted by Mr. Farzad Forouzanfar, Dr. Maryam Ramezani, Mrs. Arezoo Bibak and Ms. Zeinab Salehi.

For this research, a total of thirteen human skeletons were made available.

##### 7.2. The archaeological and anthropological characteristics of the graves

###### 7.2.1. Trench F-12, Grave 63

Grave description:
Grave 63 is located in the western part of the Marsin-Chal cemetery, situated between the southern half of trench F 10 and the northern half of trench F 11. The depth of the grave from a fixed point of measurement from the present day surface of the excavation areas is -378 centimetres.

Objects in the grave:

Within this burial site, a collection of eight distinct grave goods was unearthed. This assemblage encompasses an iron earring, several beads for a necklace, an iron brooch, a set of iron bracelets, a diminutive pottery jug, a ceramic bead intended for an armband, and an iron ring designed for a finger.

Anatomical and chronological assessments: According to the anatomical assessment, this individual (IRN57) was a female (genetically and anthropologically) and estimated to have died in the age range of 25 to 40 years. The direct $^{14}C$ radiocarbon date (405-230 cal BCE) (Supplementary Table S1) confirms the dating of the burial to the Achaemenid-Seleucid Periods.

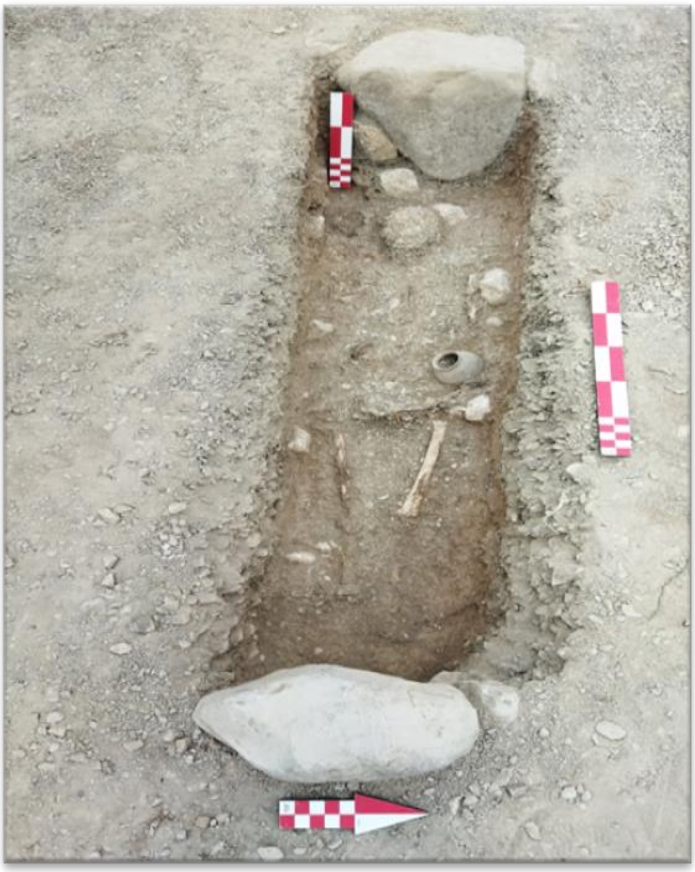

**Figure 7.1. The structure of the grave, the burial situation and the placement of the grave goods in Grave 63 (Photo by Ata Hasanpour)**

##### 7.2.2. Trench F-1, Grave 52

Grave description:
Grave 52 is located in the western part of the Marsin-Chal cemetery, positioned in the northern and western half of Trench F 11. The depth of the grave's surface from the fixed measurement point is -417 centimetres.

Objects in the grave:

Accompanying the burial in Grave 52, 14 grave goods were placed inside. These include a small, closed-mouth pot, silver earrings, necklace beads, crafting tools such as probable pottery spindle whorls, iron bracelets, a perforated bronze bell, a bronze rod, anklets, and an iron ring. Spindle whorls were also common in later periods such as in Parthian graves belonging to Liār-Sang-Bon cemetery.

Anatomical and chronological assessments:

This individual (IRN58) was identified as a female who died between 18 to 22 years old. This individual, based on the archaeological context, dates back to 355 BCE - 280 BCE, corresponding to the Achaemenid-Seleucid Period.

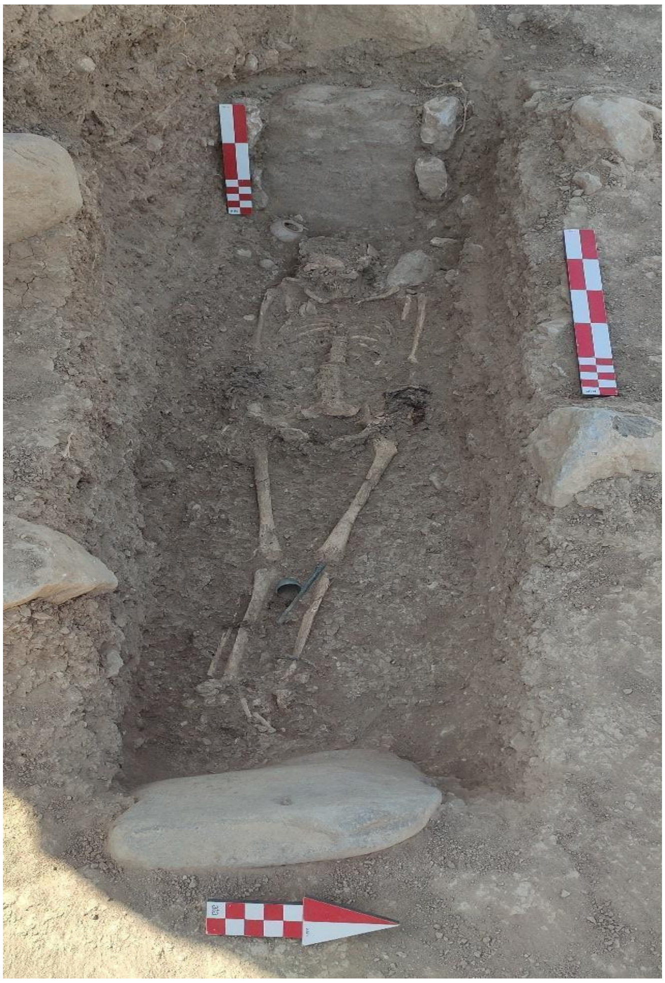

**Figure 7.2. The structure of the grave, the condition of the burial, and the placement of the grave goods within Grave 52 (Photo by Ata Hasanpour).**

##### 7.2.3. Trench F-17, Grave 47

Grave description:

Grave 47 is located in the western part of the Marsin-Chal cemetery, situated in the northern and eastern half of Trench E 12, at a depth of -323 centimetres from the fixed point of measurement from the present-day surface of the excavation.

Objects in the grave:

Along with the burial, 10 grave goods were placed inside the grave. These items include a pottery vessel, a small jug, a silver earring with a grape-shaped pendant, necklace beads, a bronze bead, iron bracelets, a bronze bell, and an iron anklet.

Anatomical and chronological assessments:

Anatomical evaluations indicate that the skeleton (IRN59) discovered in this burial belongs to a female, estimated to have died in the age range of 50 to 60 years. This individual, based on the archaeological context, dates back to 355 BCE - 280 BCE, corresponding to the Achaemenid- Seleucid Period.

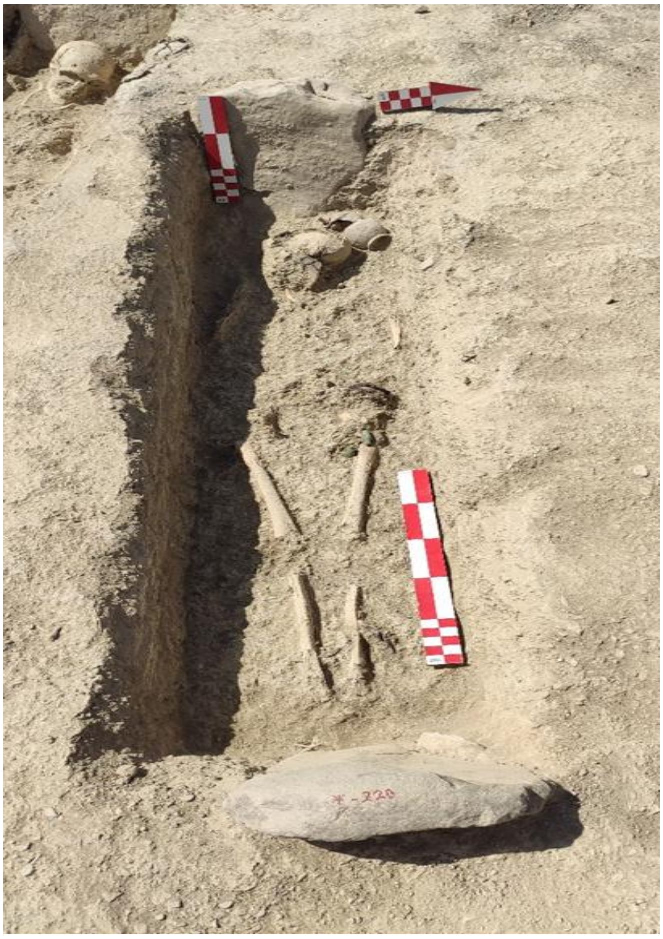

**Figure 7.3. The structure of the grave, its burial condition, and the arrangement of the grave goods within Grave 47 (Photo by Ata Hasanpour).**

##### 7.2.4. Trench F-11, Grave 41

Grave description:

Grave 41 is located in the western part of the Marsin-Chal cemetery, situated in the southern half of Trench E 12, at a depth of -43 centimetres from the fixed point of measurement from the present-day surface of the excavation areas.

Objects in the grave:

Alongside the interment, a total of 28 grave items were laid within Grave 41. These items consist of bronze earrings, various beads for a necklace, ceramic beads (likely used as part of an armband), iron bracelets and rings, another ceramic bead, an iron item identified as a spearhead, a stone bead, a bronze animal statuette, a bronze bell, additional stone beads, an iron anklet, a mixture of iron and bronze rings, and a small, orange pottery bowl and jug.

Anatomical and chronological assessment:

This individual (IRN60) was identified as female (genetically and anthropologically) and is estimated to have died between 25 to 40 years old. This individual, based on the archaeological context, dates back to 355 BCE - 280 BCE, corresponding to the Achaemenid-Seleucid Period.

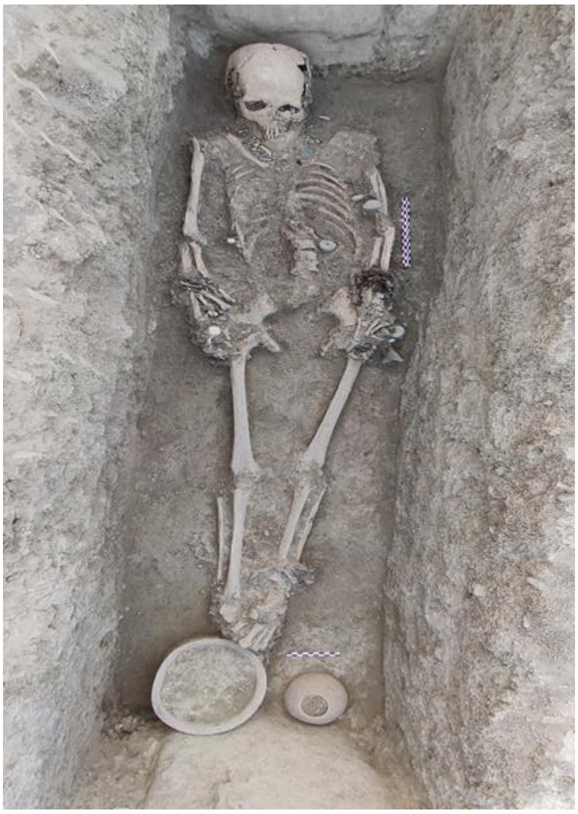

**Figure 7.4. The structure of the grave and the arrangement of the grave goods within the Grave 41 (Photo by Dr Ata Hasanpour).**

#### 7.2.5. Trench F-4, Grave 55

Grave description:

Grave 55 is located in the western part of the Marsin-Chal cemetery, situated in the southern and western half of Trench F 10, at a depth of -341 centimetres from the fixed point of measurement from the present day surface of the excavation area.

Objects in the grave:

This burial had no grave goods.

Anatomical and chronological assessments:

This individual (IRN61) is identified as male and is estimated to have died between 50 to 60 years of age. This individual, based on the archaeological context, dates back to 355 BCE- 280 BCE, corresponding to the Achaemenid-Seleucid Period.

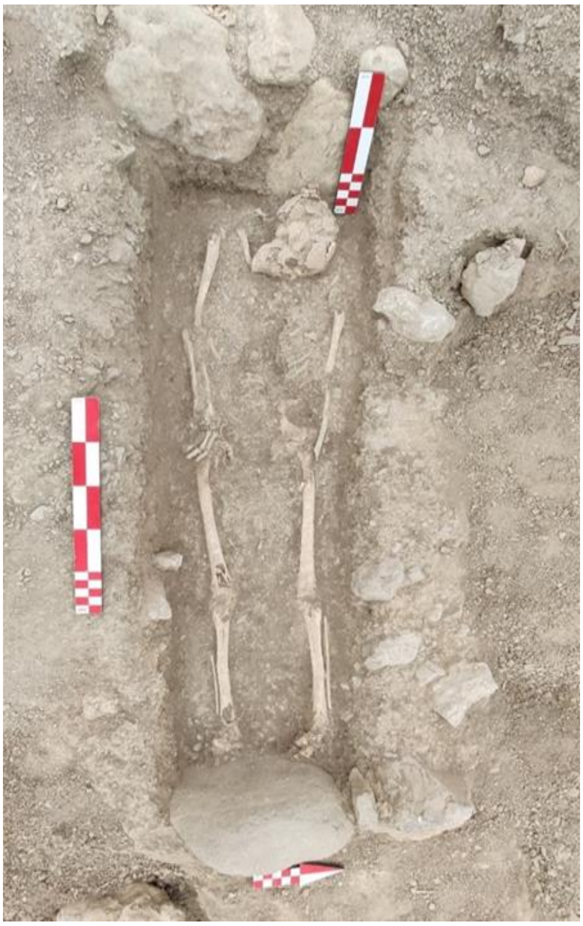

**Figure 7.5. The structure of Grave 55 and the specifics of its burial arrangement (Photo by Ata Hasanpour).**

##### 7.2.6. Trench F-8, Grave 59

Grave description:

Grave 59 is situated in the southern half of trench F11 and in the western part of the Marsin-Chal cemetery. It is worth noting that the depth of this grave is 331 centimetres below the fixed measurement point.

Objects in the grave:

In this burial, three grave goods were discovered. These items included an iron arrowhead, a small clay pot, and an iron dagger.

Anatomical and chronological assessment:

This individual (IRN62) is identified as female and is estimated to have died between 20 to 25 years of age. This individual, based on the archaeological context, dates back to 355 BCE - 280 BCE, corresponding to the Achaemenid-Seleucid Period.

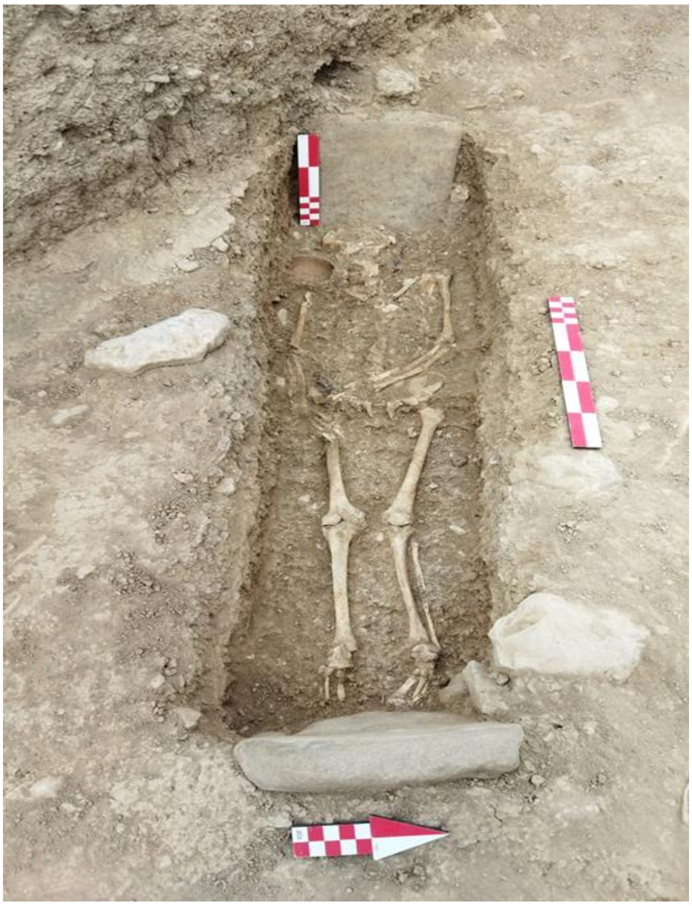

**Figure 7.6. The structure of the grave and the burial arrangement in Grave 59 (Photo by Ata Hasanpour).**

**7. 2.7. Trench F-9, Grave 30**

Grave description:

Grave 30 is located in the northern half of trench D 13 at a depth of 60 centimetres below the fixed point of measurement from the present-day surface of the excavation areas.

Objects in the grave:

Along with the burial in Grave 30, three items were placed inside as burial offerings. These include an iron arrowhead, a bronze ring, and an orange-coloured earthenware vessel.

Anatomical and chronological assessment:

This individual (IRN63) is identified as male and is estimated to have died between 20 to 25 years of age. This individual, based on the archaeological context, dates back to 355 BCE - 280 BCE, corresponding to the Achaemenid- Seleucid Period.

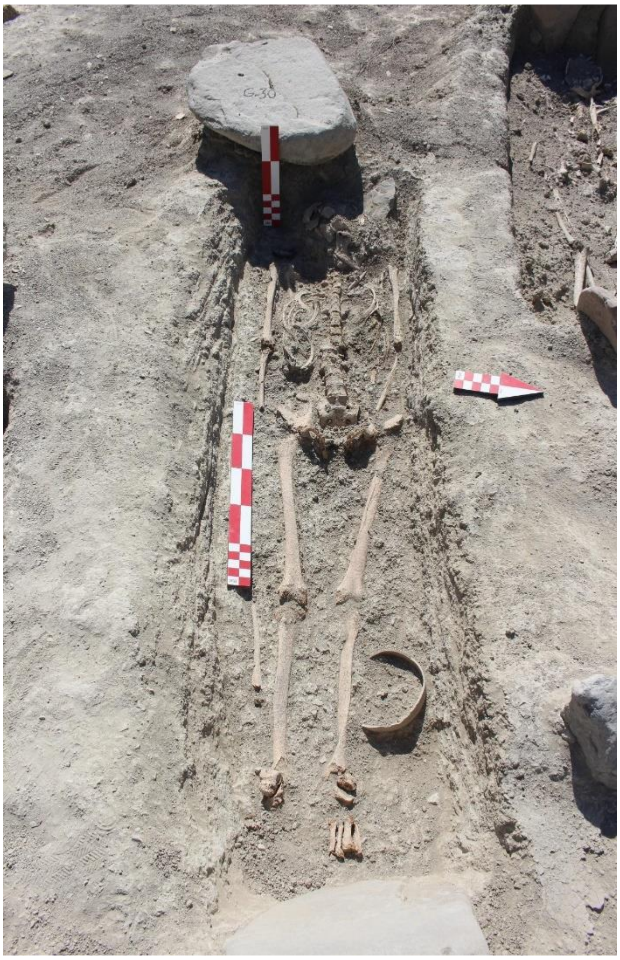

**Figure 7.7. The structure of the grave, the burial arrangement, and the positioning of the grave goods in Grave 30 (Photo by Ata Hasanpour).**

##### 7.8.2. Trench F-13, Grave 43

Grave description:

Grave 43 is located in the southern and eastern part of trench E 12, positioned at a depth of 210 centimetres below the fixed point of measurement from the present-day surface of the excavation areas.

Objects in the grave:

Accompanying the burial in Grave 43, three items were placed as burial offerings. These items include a clay bowl, a bronze earring, and an iron dagger.

Anatomical and chronological assessment:

This individual (IRN64) is identified as female and is estimated to have died between 18 to 35 years of age. This individual, based on the archaeological context, dates back to 355 BCE -280 BCE, corresponding to the Achaemenid-Seleucid Period.

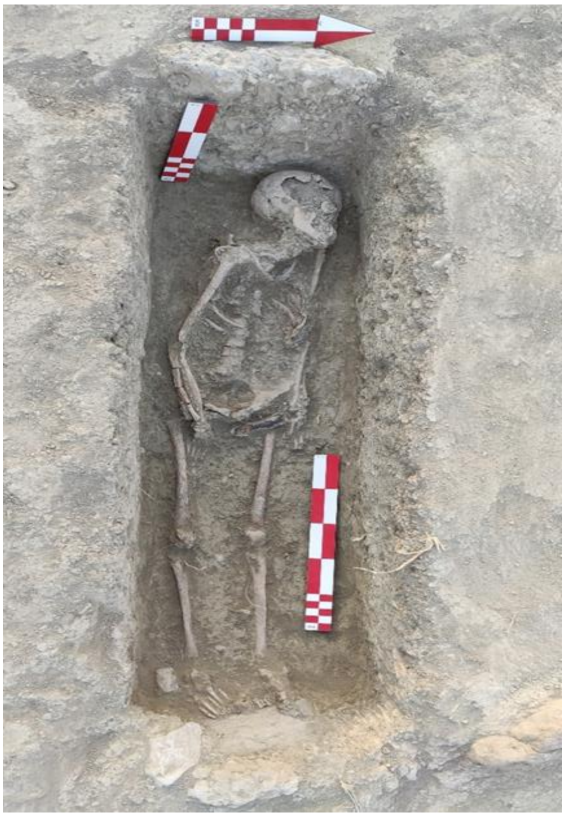

**Figure 7.8. The structure of the grave, the burial arrangement and the positioning of the grave goods in Grave 43 (Photo by Ata Hasanpour).**

**7.2.9. Trench F-3, Grave 54**

Grave description:

Grave 54 is situated in the northern and western portion of trench F 11, at a depth of 326 centimetres below the fixed point of measurement from the present-day surface of the excavation areas.

Objects in the grave:

There was an iron sword placed above the head of the deceased.

Anatomical and chronological assessment:

According to the anatomical assessment, this individual (IRN65) is male (genetically and anthropologically) and estimated to be in the age range of 40 to 45 years. This individual, based on the archaeological context, dates back to 355 BCE- 280 BCE, corresponding to the Achaemenid- Seleucid Period.

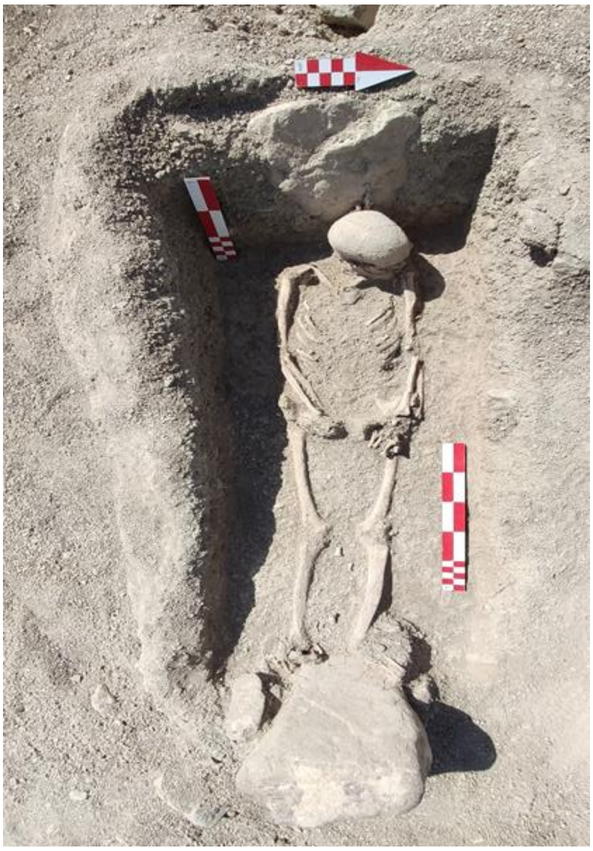

**Figure 7.9. The grave structure, burial arrangement and positioning of grave goods in Grave 54 (Photo by Ata Hasanpour).**

##### 7.2.10. Trench F-19, Grave 49

Grave description:

Grave 49 is located in the southern and eastern half of trench E 12, at a depth of 168 centimetres below the fixed point of measurement from the present-day surface of the excavation areas.

Objects in the grave:

There was only a single iron spearhead within this grave.

Anatomical and chronological assessment:

According to the anatomical assessment, this individual (IRN66) is likely male and estimated to have died between 70 to 75 years old. This individual, based on the archaeological context, dates back to 355 BCE -280 BCE, corresponding to the Achaemenid- Seleucid Period.

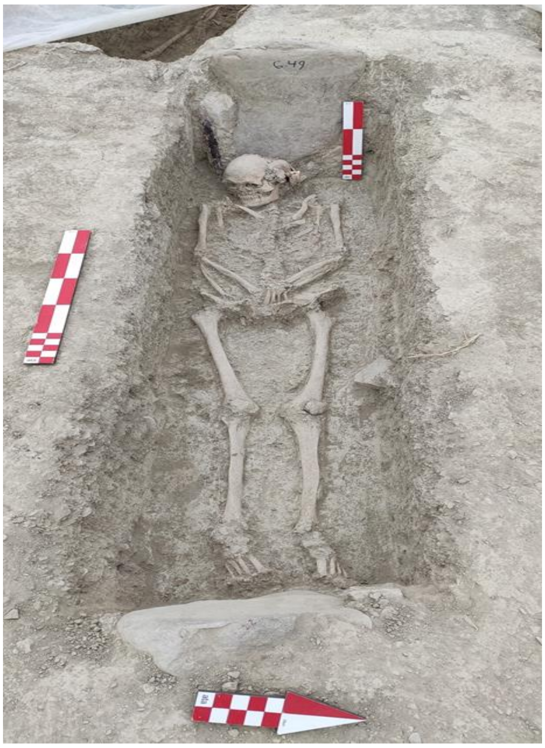

**Figure 7.10. The grave structure, burial arrangement, and positioning of grave goods in Grave 49 (Photo by Ata Hasanpour).**

##### 7.2.11. Trench F-3, Grave 7

Grave description:

Grave 7 is located in the northern half of trench D 12.

Objects in the grave:

In this burial, two grave goods were placed. These items included a clay bowl and an iron sword.

Anatomical and chronological assessment:

According to the anatomical assessment, this individual (IRN67) is likely male and estimated to have died between 45 and 50 years of age. This individual, based on the archaeological context, dates back to 355 BCE - 280 BCE, corresponding to the Achaemenid- Seleucid Period.

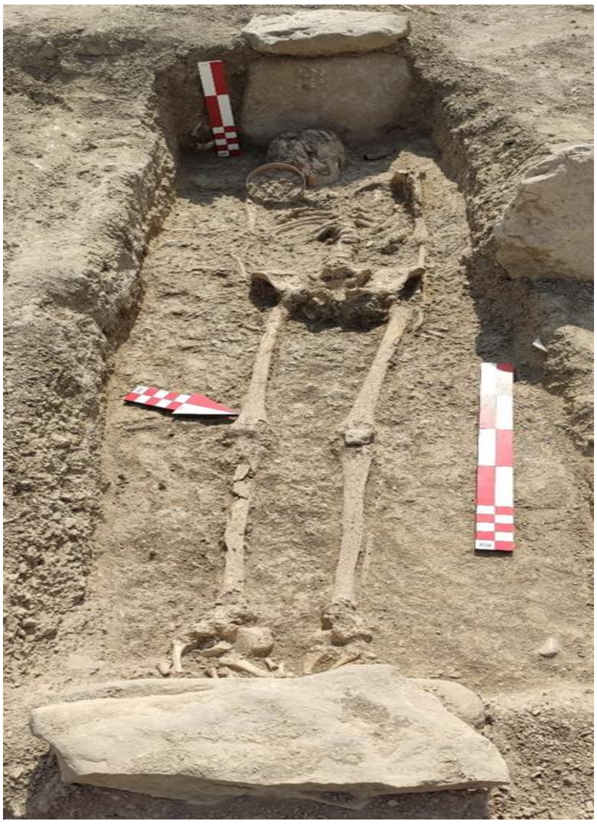

**Figure 7.11. The burial arrangement and positioning of grave goods in Grave 7 (Photo by Ata Hasanpour).**

##### 7.2.12. Trench F-8, Grave 28

Grave description:

Grave 28 is situated in the southern half of trench D 13.

Objects in the grave:

Alongside the burial, three grave offerings were placed inside the grave. These grave goods consisted of a tripod ceramic ritual vessel placed on the western side of the grave, a ceramic cup, and an iron dagger blade.

Anatomical and chronological assessment:

According to the anatomical assessment, this individual (IRN68) is likely male and estimated to have died between 18 to 20 years of age. This individual, based on the archaeological context, dates back to 355 BCE- 280 BCE, corresponding to the Achaemenid- Seleucid Period.

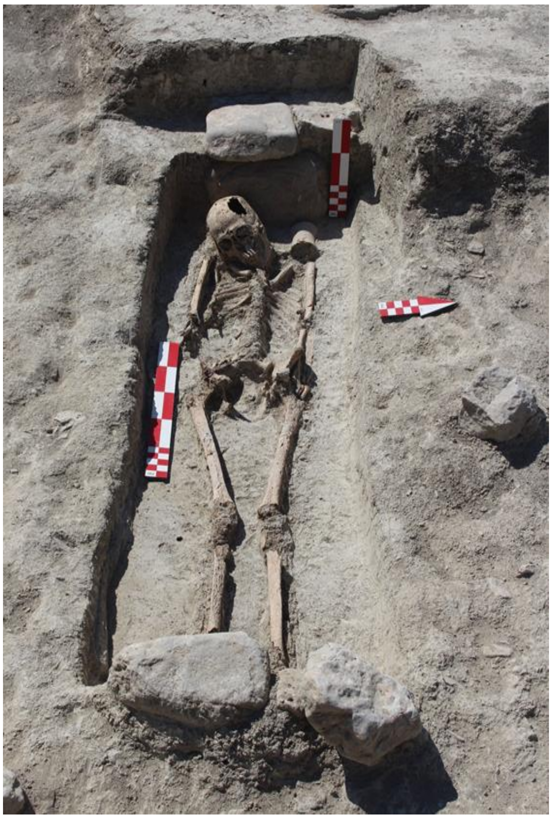

**Figure 7.12. The grave structure, burial arrangement, and positioning of grave goods in Grave 28 (Photo by Ata Hasanpour).**

##### 7.2.13. Trench F-8, Grave 28

Grave description:

Grave 46 is situated in the western part of the Marsin-Chal cemetery.

Objects in the grave:

Accompanying the burial, four grave offerings were placed inside the grave. These grave goods included a tripod ceramic bowl placed at the foot of the grave, an iron sword placed above the head, an iron dagger, and an iron spearhead positioned between the feet and under the pelvic bone.

Anatomical and chronological assessment:

According to the anatomical assessment, this individual (IRN69) is male (genetically and anthropologically) and estimated to be in the age range of 40 to 45 years. The direct $^{14}C$ radiocarbon date (405-230 cal BCE) (Supplementary Table S1) confirms the dating of the burial to the Achaemenid- Seleucid period.

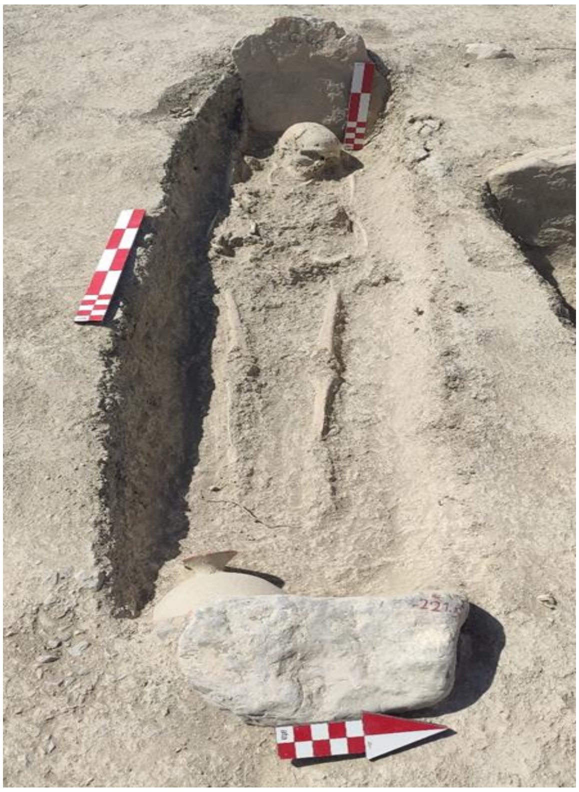

**Figure 7.13. The grave structure, burial arrangement, and positioning of grave goods in Grave 46 (Photo by Ata Hasanpour).**

#### Chapter 8: Liār-Sang-Bon site

The information of this chapter was collected by the excavator of Liār-Sang-Bon cemetery, Dr. Vali Jahani and his colleague Ms. Mana Rouhani. This section was drafted by Motahareh Amjadi and Ms. Arezoo Bibak.

##### 8.1. Site description

The ancient site of Liār-Sang-Bon, located in the Sumam district of Rankuh county in Amlash, Gilan province, is an extensive area with geographic coordinates of 36.581948°N latitude and 50.53396°E longitude. Situated between the villages of Shirchak, Shihe, and Siyakuhand Marbo, the site spans approximately 27 hectares. It consists of a cemetery section on the eastern slope of the valley and a residential area on the western side of the valley, along the southern ridge of the rock outcrop known as Liār-Sang-Bon. Additionally, scattered evidence of settlements from the historical period to the later Islamic centuries has also been identified in the northern, northeastern, and southern parts of the site.

The topography of the region has played a fundamental role in the distribution of the aforementioned archaeological sites. The altitude of this area ranges from 1750 to 1900 metres above sea level and its distance from Amlash city is 75 kilometres via the Garmabdasht - Rahimabad area and 57 kilometres via the Blourdekan Amlash area.

This archaeological complex was first identified in the year 2012 where field studies were carried out over three seasons in 2014, 2016, and 2017, 2018, 2019, and 2020 under the supervision of Dr. Vali Jahani and his colleagues, which yielded a total of 130 catacombs, pit, and jar types of burials (Jahani et al.,2023. Jahani et al., 2024). Radiocarbon dates obtained from dentine collagen extracted from two tombs discovered in the 2016 and 2017 excavation seasons fall within the ranges of 38 cal BC-123 cal AD and 45 cal BC-80 cal AD, aligning the cemetery with the Parthian era (Sołtysiak and Jahani, 2019). Excavations at Liār-Sang-Bon have yielded a large assemblage of burial goods, including various pottery types, iron weaponry, personal accessories, and an array of personal ornaments. The latter consists of various kinds of beads, pendants, rings, finger rings, bracelets, earrings, pins, and medallions, made of various materials such as frit, glass, bronze, silver, gold, and bitumen (Jahani et al.,2023 and Jahani et al., 2024).

For this research, a total of eight human skeletons from the excavations conducted in 2017 were provided to us.

##### 8.2. The archaeological and anthropological characteristics of the Catacombs

###### 8.2.1. Trench 9603, Catacomb 96303

Catacomb description:

This grave, categorised as a catacomb type, is situated in the southeastern quarter of the excavation site. The skeleton recovered from inside this grave was buried on its left side with the legs bent. The bed of the grave, which slopes gently from east to west, measures approximately -278 centimetres from a fixed point of measurement from the present-day surface of the excavation areas.

Objects in the Catacomb:

In this burial, a total of 43 grave goods were discovered. These items included a variety of ceramic vessels and diverse decorative items, such as beads made from agate and gold, along with various types of filigree and pendants.

Anatomical and chronological assessments:

Anatomical evaluations indicate that the skeleton (IRN21) discovered in this burial belongs to a female, estimated to have died between 25 to 40 years of age. The dating of this grave, based on archaeological context, is between 200 BC to 100 CE, corresponding to the Parthian Period.

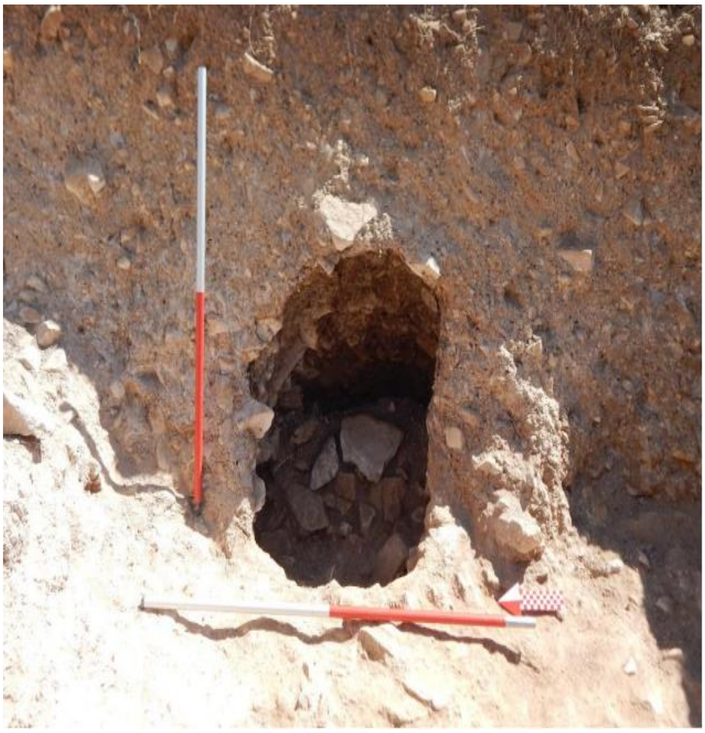

**Figure 8.1. The entrance of the catacomb 96303 (Photo by Vali Jahani).**

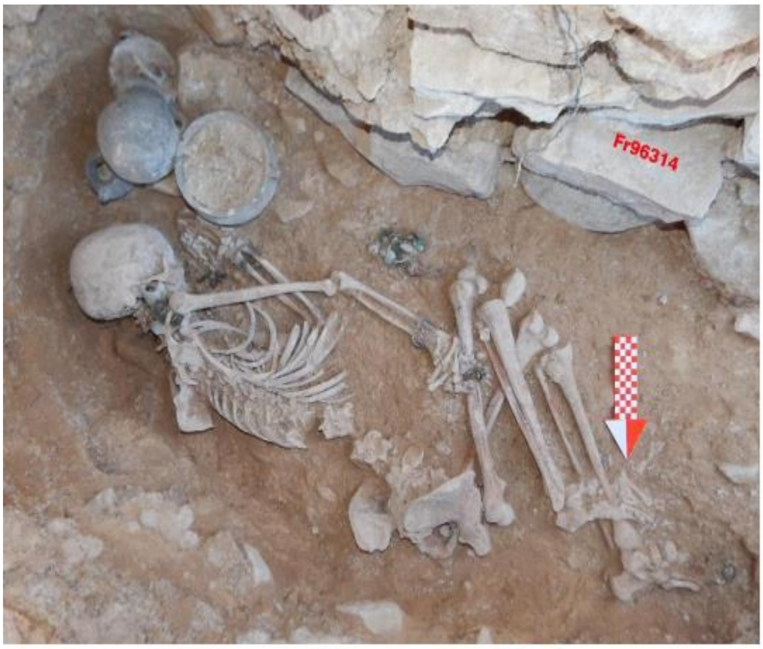


**Figure 8.2. Burial condition and the arrangement of grave goods in the Catacomb 96303 (Photo by Vali Jahani).**

##### 8.2.2. Trench 9603, Catacomb 96317

Catacomb description:

This grave, characterised as a pit-type burial, is located in the north-eastern quarter of the excavation area and extends into the northern section of the site. The burial recovered from this grave (Supplementary Figures 8.3 and 8.4) was interred in a manner similar to a supine position. The depth of the grave from a fixed point of measurement from the present-day surface of the excavation areas is -195 centimetres.

Objects in the Catacomb:

In this burial, a total of 15 grave goods were discovered. These items included a jug, a jar, a tripod bowl, an iron knife and needle, decorative beads, earrings, and a bronze pendant.

Anatomical and chronological assessments:

According to the anatomical assessment, this individual (IRN22) is female (genetically and anthropologically) and estimated to have died in the age range of 19 to 35 years. The direct $^{14}C$ radiocarbon date (165-1 cal BCE) (Supplementary Table S1) confirms the assignment of the burial to the Parthian Period.

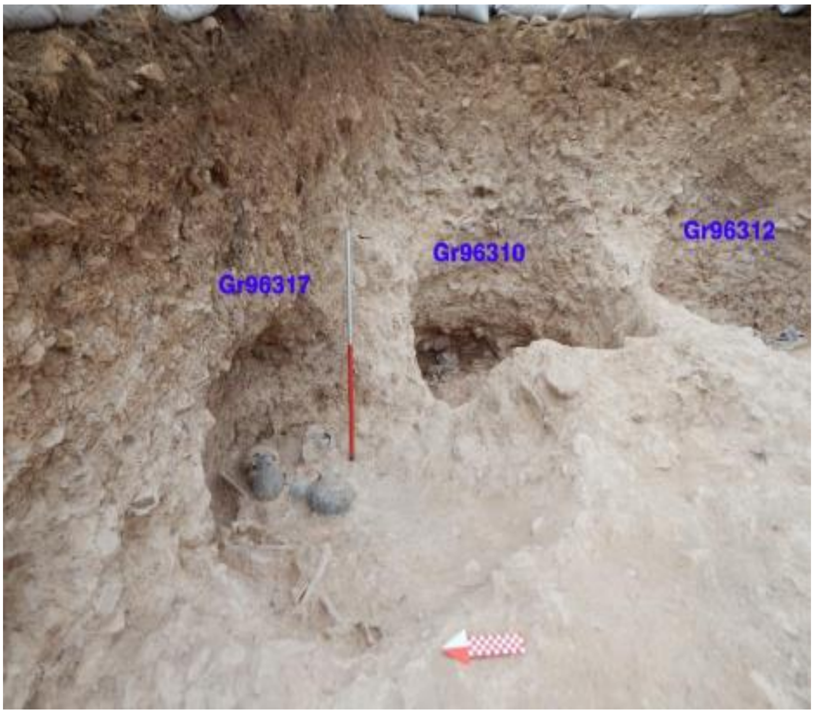


**Figure 8.3. The location of Catacomb 96317 within the excavation site (Photo by Vali Jahani).**

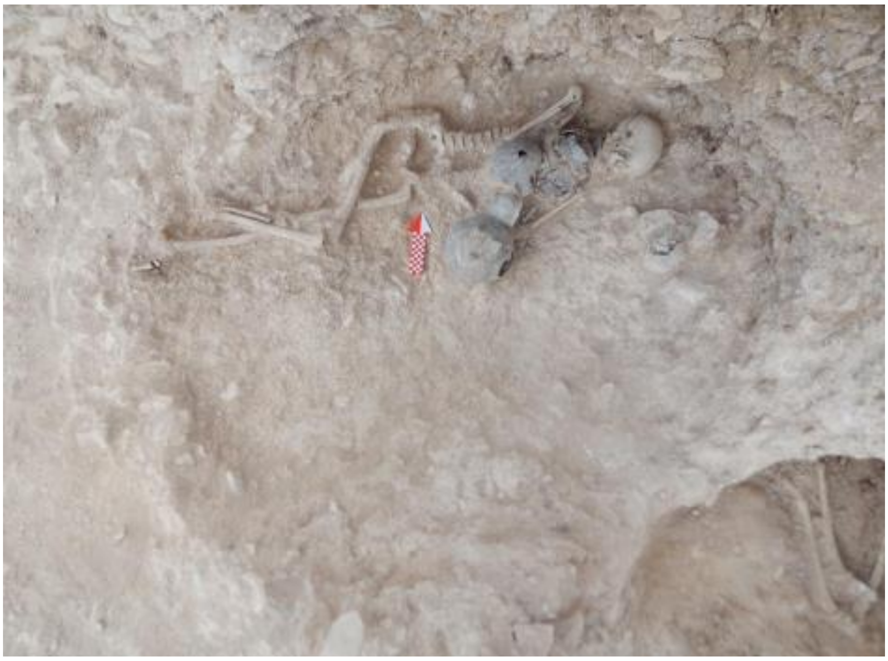

**Fig 8.4. The burial condition of Catacomb 96317 (Photo by Vali Jahani).**

#### 8.2.3. Trench 9603, Catacomb 96301

Catacomb description:

This pit grave is situated in the northeastern quadrant of the Trench 9603 and lies at a depth of -198 centimetres from a fixed point of measurement from the present-day surface of the excavation areas. The individual buried within it was laid on their left side, with the legs positioned relatively straight.

Objects in the Catacomb:

In this burial, four grave goods were discovered. These items included a spearhead, an iron knife, a jug, and a ceramic bowl.

Anatomical and chronological assessments:

This individual (IRN23) was identified as male (genetically and anthropologically) and is estimated to have died between the ages of 45 to 60 years. The dating of this grave based on archaeological context is between 200 BC and 100 CE, corresponding to the Parthian Period. We assessed kinship relatedness among samples, and we determined that IRN23 had twin/identical level allele sharing, and identical maternal and paternal lineages with IRN25. This observation corresponded to anthropological reports which implied that these two individuals were probably twins.

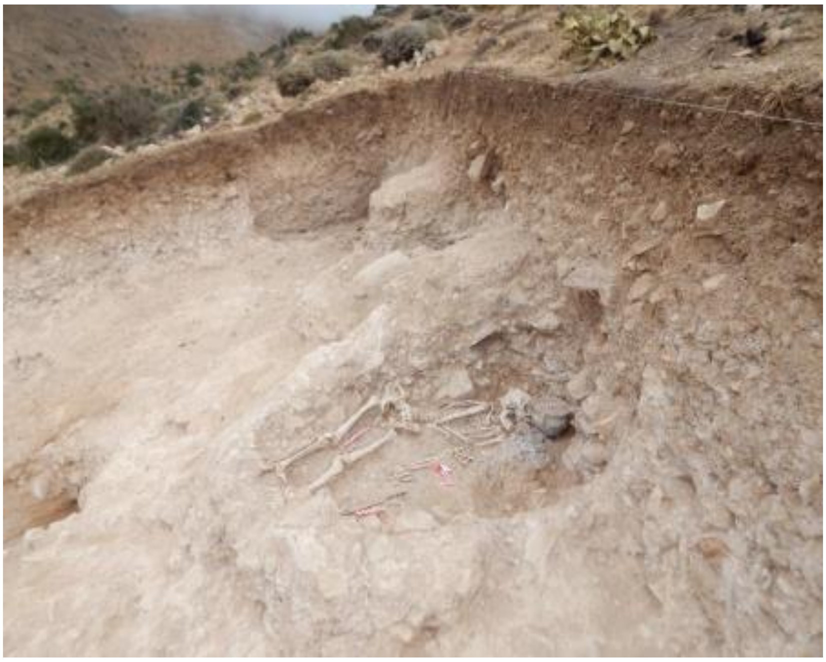

**Figure 8.5. The location of the Catacomb 96301 (Photo by Vali Jahani).**

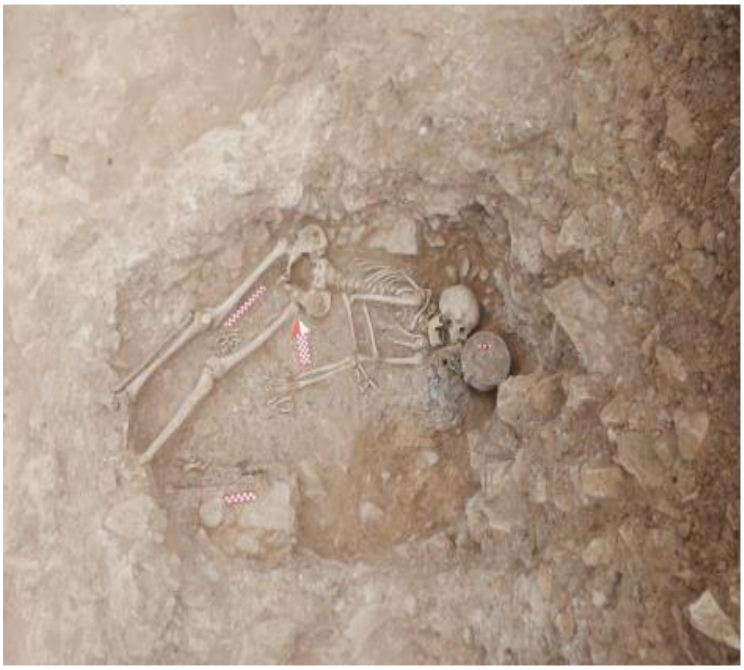

**Figure 8.6. The burial condition of Catacomb 96301 (Photo by Vali Jahani).**

##### 8.2.4. Trench 9602, Catacomb 96202

Catacomb description:

This grave is of the pit-type, located in the trench 9602. The burial recovered from this grave was laid on the left side with legs drawn up. The depth of the grave from a fixed point of measurement from the present-day surface of the excavation areas is -197 centimetres.

Objects in the Catacomb:

In total, four items were discovered from this burial, comprising a jug, a small bowl, an iron knife, and a trapezoidal tripod bowl.

Anatomical and chronological assessments:

This individual (IRN28) is identified as probably female and estimated to be between 11 to 16 years old. The dating of this individual based on archaeological context, dates back to 200 BC to 100 CE, corresponding to the Parthian Period.

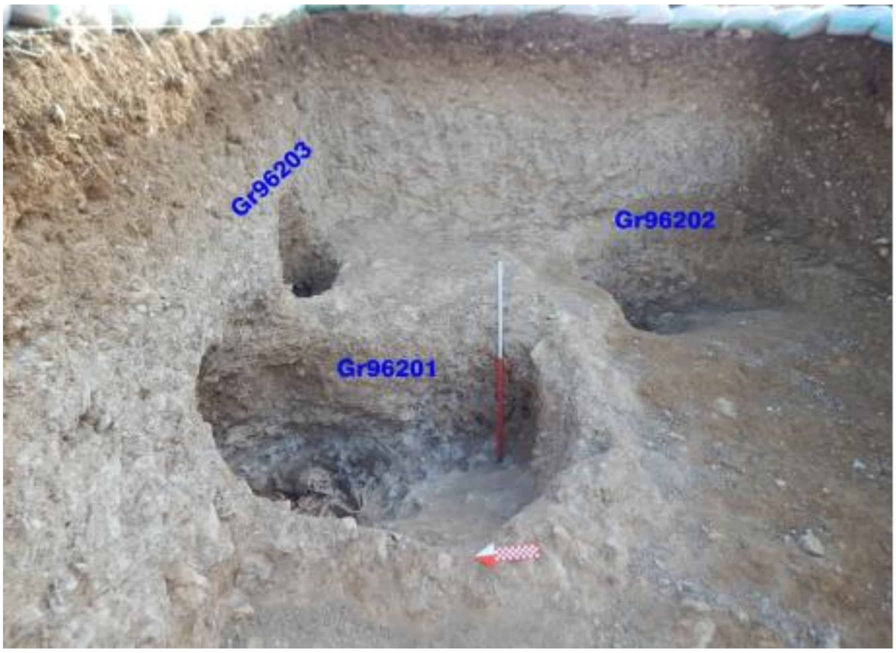


**Fig 8.7. Location and positioning of Catacomb 96202 (Photo by Vali Jahani).**

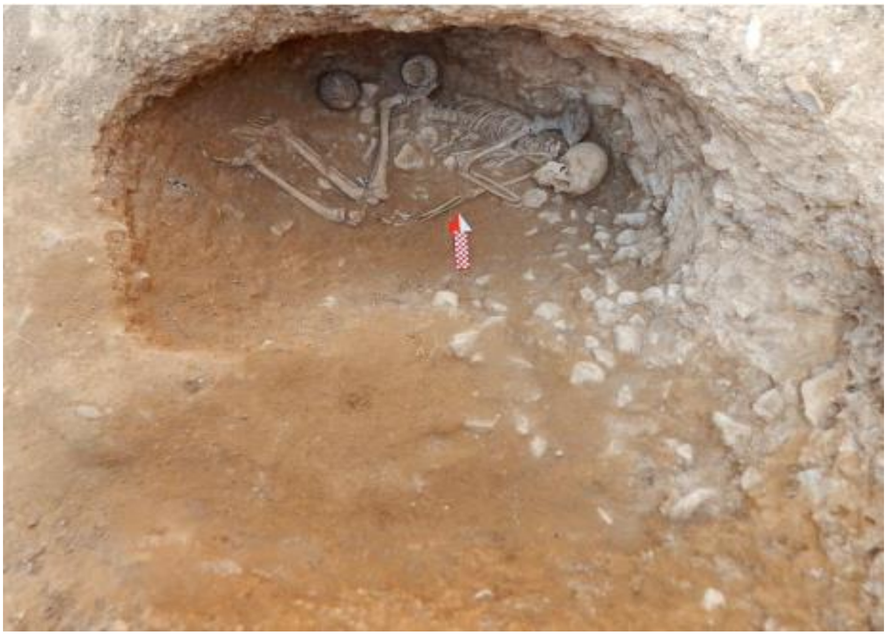

**Fig 8.8. Grave structure and burial condition of Catacomb 96202 (Photo by Vali Jahani).**

##### 8.2.5. Trench 9602, Catacomb 96203

Catacomb description:

This grave, identified as a catacomb type, is situated in the northern section of trench 9602. The interred individual was found in a contracted position, lying on the left side. The grave's depth from a fixed point of measurement from the present-day surface of the excavation area's is recorded at -110 centimetres.

Objects in the Catacomb:

In total, five grave goods were discovered from this burial, including a tripod bowl, a dagger, earrings, a spearhead, and a knife.

Anatomical and chronological assessments:

This individual (IRN31) was identified as male (genetically and anthropologically) and estimated to be between 20 to 35 years old. The direct 14C radiocarbon date (calibrated 50 BCE-65 CE) (Supplementary Table S1) confirms the allocation of the burial to the Parthian Period.

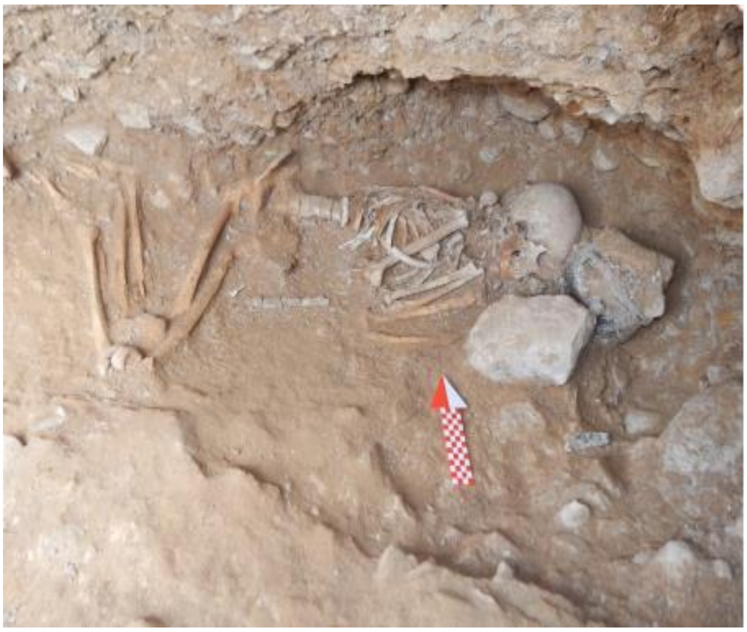

**Figure 8.9. The burial condition and grave goods within Catacomb 96203 (Photo by Vali Jahani).**

##### 8.2.6. Trench 9601, Catacomb 96101:

Catacomb description:

This grave, identified as a pit-type, was located in the northeastern corner of trench 9601. The individual interred within this grave was positioned on their left side with the legs drawn up. The depth of the grave surface, as measured from a fixed point of measurement from the present-day surface of the excavation area, is -183 centimetres.

Objects in the Catacomb:

In total, four objects were discovered from this burial, which included a spearhead, a bowl, and a pair of earrings.

Anatomical and chronological assessments:

This individual (IRN34) is identified as male (genetically and anthropologically) and estimated to have died between 25 to 30 years of age. The dating of this grave, based on archaeological context, is from 200 BC to 100 CE, corresponding to the Parthian Period.

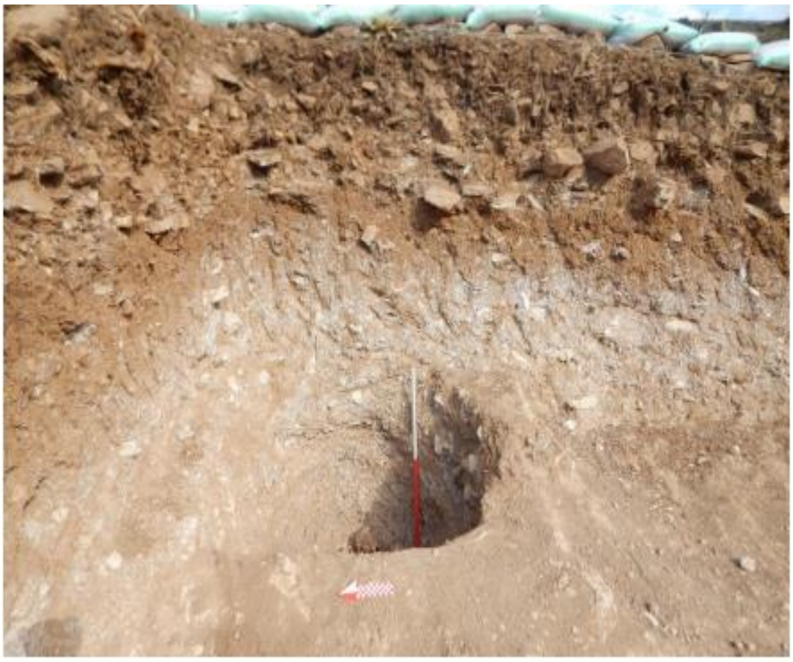

**Figure 8.10. Location of Catacomb 96101 in the northeastern quadrant of the excavation site (Photo by Vali Jahani).**

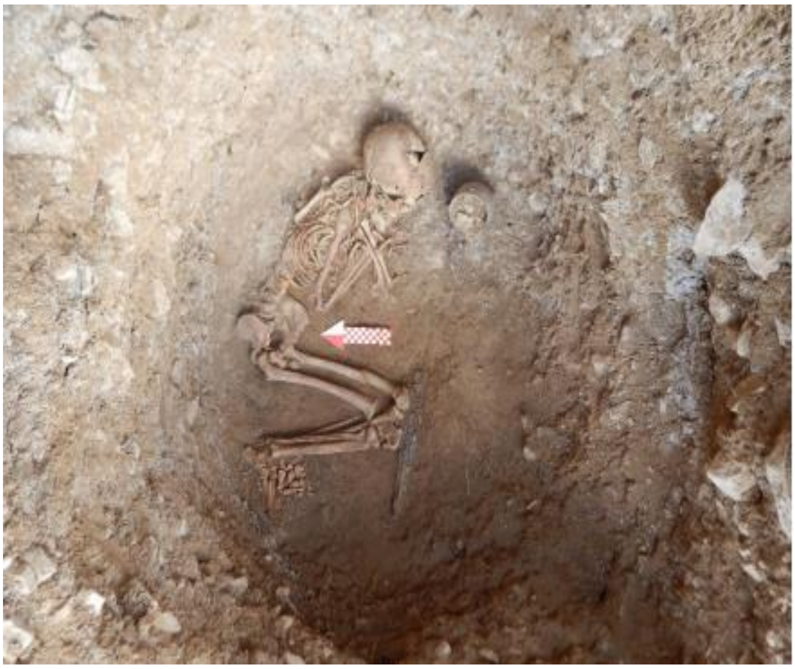

**Figure 8.11. The burial condition and grave goods within Catacomb 96101 (Photo by Vali Jahani).**

##### 8.2.7. Trench 9602, Catacomb 96201

Catacomb description:

This grave, of the pit-type variety, was situated in the middle of the northern side of trench 9602. The burial recovered from this grave was positioned on its back with legs drawn up. The depth of the grave bed from a fixed measurement point from the present-day surface of the excavation areas is -350 centimetres.

Objects in the Catacomb:

In total, seven grave goods were discovered from this burial, including a spearhead, a dagger, an arrowhead, beads, and an iron knife.

Anatomical and chronological assessments:

This individual (IRN26) is identified as probably male and estimated to have died between 25 to 40 years of age. The dating of this grave, based on archaeological context, is from 200 BCE to 100 CE, corresponding to the Parthian Period.

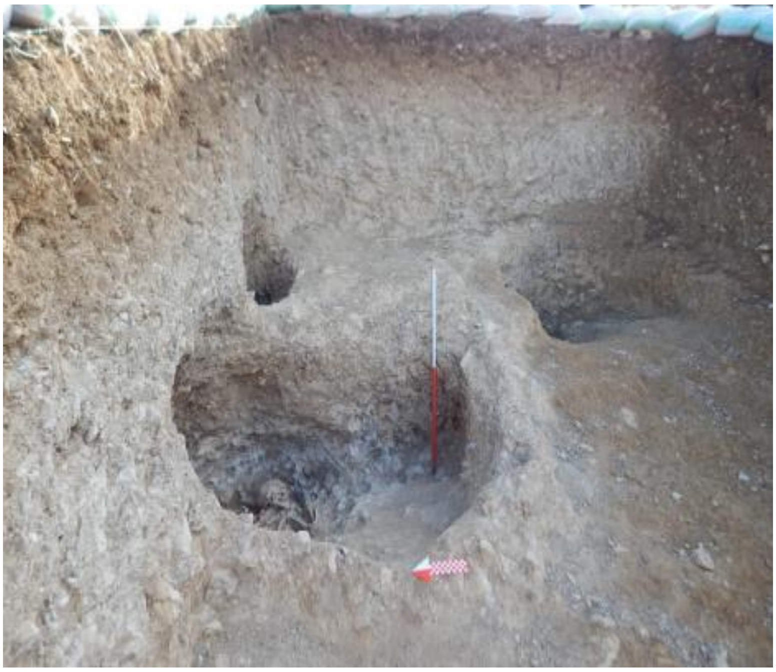

**Figure 8.12. Location of Catacomb 96201 within the excavation site (Photo by Vali Jahani).**

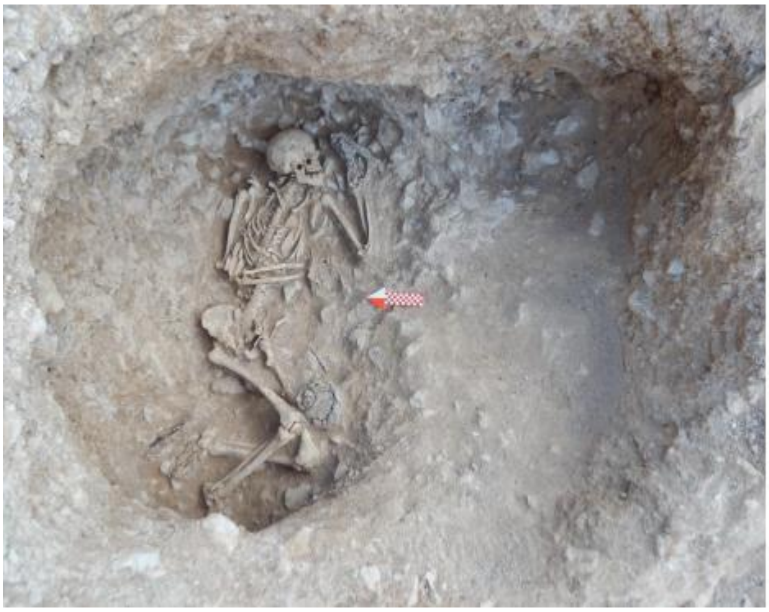

**Figure 8.13. The burial condition and structural aspects within Catacomb 96201 (Photo by Vali Jahani).**

##### 8.2.8. Trench 9603, Catacomb 96309

Catacomb description:

This grave, identified as a catacomb type, is located in the southeastern quadrant of the trench 9603. The skeleton recovered from within this grave was buried on its left side with legs drawn up. The depth of the grave from a fixed point of measurement from the present-day surface of the excavation areas is -225 centimetres (Supplementary Figures 8.15 and 8.16).

Objects in the Catacomb:

From this burial, a total of eight items were discovered, including a ceramic jug, a one-handled jar, a tripod bowl, a bronze ring, a strand of glass paste beads, a ceramic spout, a needle, and an iron knife.

Anatomical and chronological assessments:

This individual (IRN25), is identified as male (genetically and anthropologically) and estimated to be between 45 to 60 years of age. The dating of this grave, based on archaeological context, is from 200 BC to 100 CE, corresponding to the Parthian period. We assessed kinship relatedness among samples, and we determined that IRN25 had twin/identical level allele sharing, with identical maternal and paternal lineages with IRN23. This observation corresponded to anthropological reports which implied that these two individuals were probably twins. The archeological material remains from these twins are exactly double those of each other, which is an interesting finding in relation to the biological and anthropological evidence.

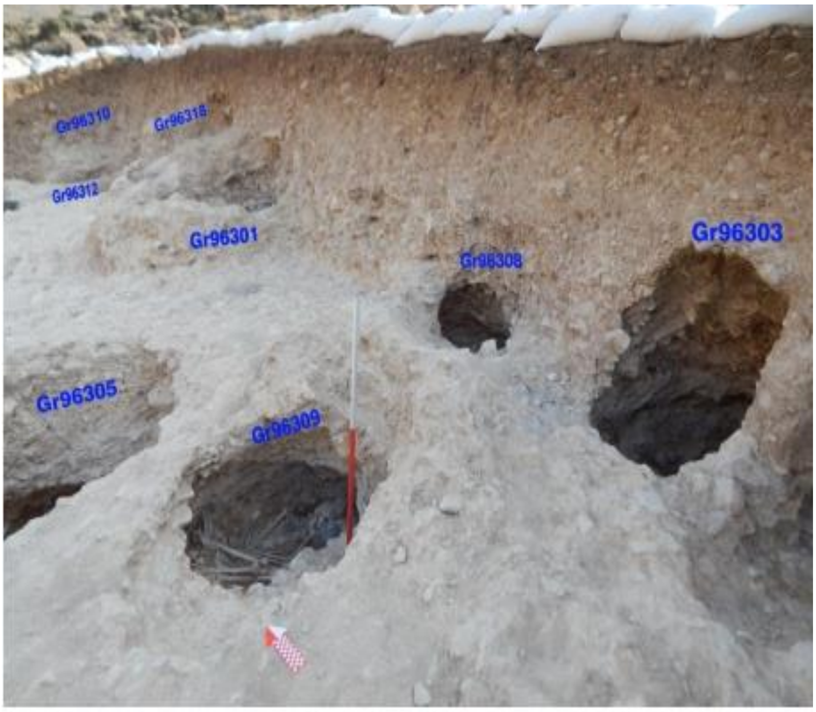


**Fig 8.15. The location of Catacomb 96309 within the excavation site (Photo by Vali Jahani).**

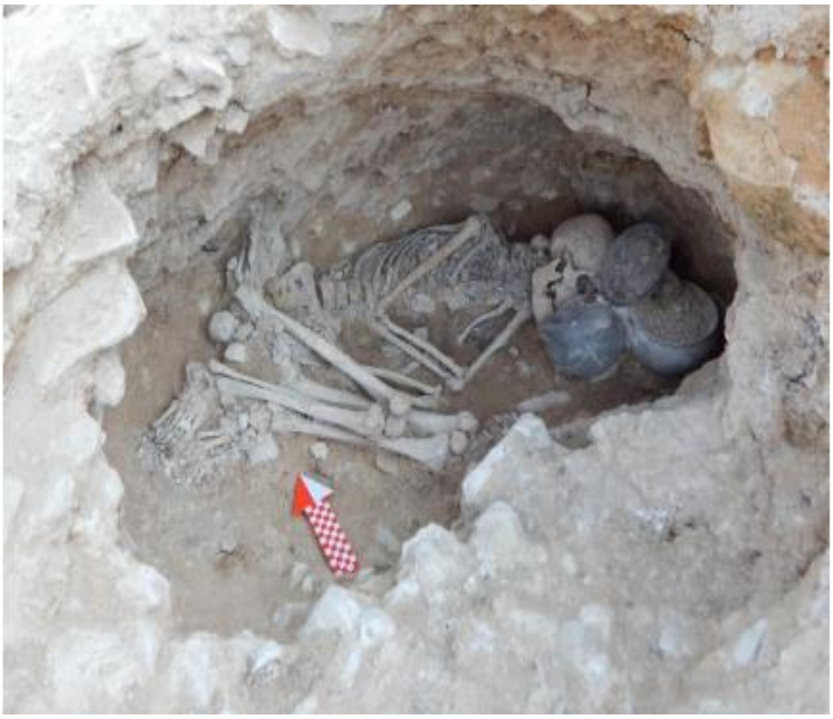

**Fig 8.16 The burial condition of Catacomb 96309 (Photo by Vali Jahani).**

#### Chapter 9: Vestemin

The information of this chapter was collected by the excavator of the Vestemin cemetery, Dr. Abdolmotalleb Sharifi.

##### 9.1. The geographical location of the Let-Sar site in the Vestemin village

The ancient site of Let-Sar, situated near the village of Vestemin, is a remarkable archaeological location in Mazandaran province, Iran. Positioned approximately 9 kilometres southeast of Kiasar, the central town of the Chahardangeh district, and 3 kilometres north of Vestemin village, Let-Sar stands as a testament to the region's historical richness. Often referred to by the name of the nearby village, Vestemin, this site includes a settlement area, two cemeteries (eastern and western) and a fort, all set against a mountainous backdrop. Geographically, Vestemin village is nestled in a mountainous terrain, bordered by the village of Terkam to the northeast, the Kiasar-to-cement-factory road to the northeast, the Sari-to-Semnan Road to the south and the region's dense forests to the west. The Let-Sar site itself is located on the slope of a gently rising hill, extending east-west over a span of 300 metres before descending into a valley to the west. To the south, a spring by the same name, Let-Sar, feeds into a shallow valley, while the northern part opens into a deeper valley where the Babr-Cheshmeh spring is found. The altitude of Let-Sar is 1389 metres above sea level, with geographical coordinates at 53.53408°N latitude and 36.24373°E longitude.
Under the direction of Dr. Abdolmotalleb Sharifi, the Vestemin site underwent extensive explorations in 2015, 2017, and 2018. These investigations encompassed both the eastern and western cemeteries, residential areas, a castle, and an Islamic cemetery. The site is notable for its continuous use from the Parthian era through to Islamic times, as evidenced by the variety of human burials discovered here.

The strategic location of the site, near the Parthian capital of Sad Darvazeh and the Silk Road, is significant. It lies approximately 51 kilometres (air distance) from Sad Darvazeh and 80 kilometers from Sari, the capital of Farashvadgar Province, along an ancient road connecting the two.

The Vestemin cemetery yielded approximately 100 burials, including three single-horse burials. A notable discovery was the presence of armaments in 42 of these burials, such as swords, daggers, arrowheads, trefoil arrowheads, and armour. Trefoil arrowheads were particularly frequent, while shields and armour were less common. Interestingly, the distribution of armaments suggests a disregard for gender distinctions in burial rites, as evidenced by the presence of a sword in a female burial and numerous daggers in other female burials.

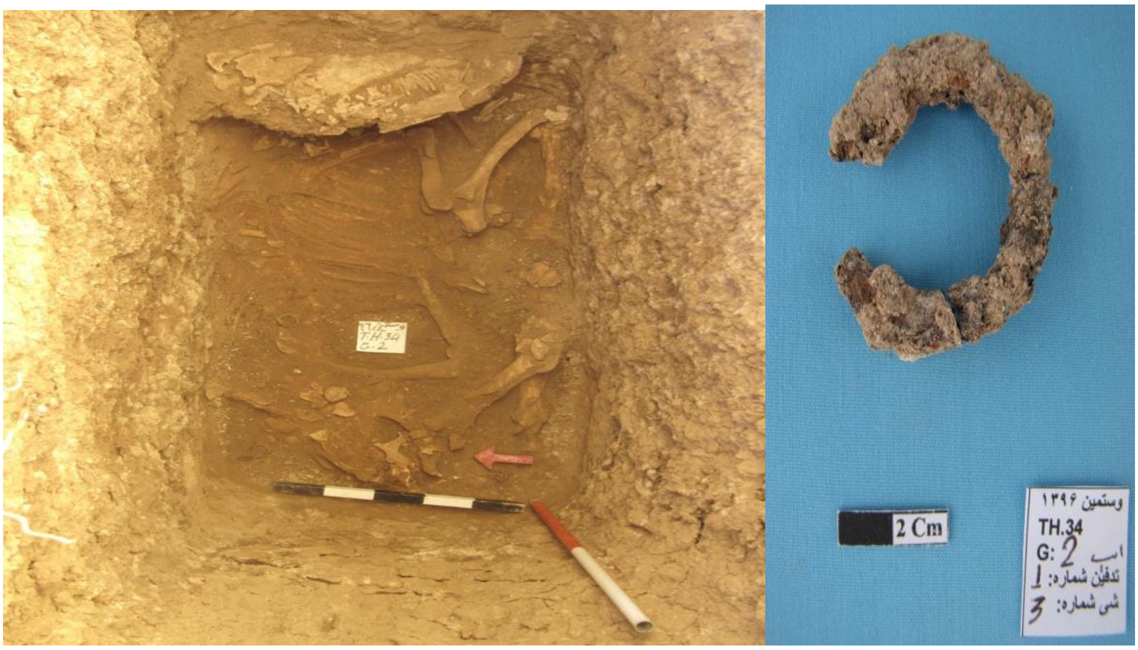


**Figure 9.1. A: Horse burial belonged to Trench H34, Grave 2, B: Iron horseshoe belonged to Trench H34, Grave 2 (Photo by Abdolmotalleb Sharifi).**

The unearthed armaments, primarily cavalry weapons, combined with the discovery of horse burials in the western cemetery, reinforce the hypothesis that the catacombs were intended for Parthian cavalrymen or their families, or at least individuals influenced by a martial culture. This finding underscores the importance of cavalry during the Parthian period. Moreover, the disparities in burial practices and armament types between the eastern and western cemeteries suggest the possibility of two distinct Parthian phases with differing military strategies (Sharifi et al., 2023).

What sets the Vestemin site apart, particularly its western cemetery, are the family catacombs, notable for their unique architectural style. These catacombs are divided into three distinct sections: a rectangular corridor, an entrance gateway connecting the corridor to the crypt room, and the crypt room itself. Inside, multiple burials have been found, with the number of individuals per grave ranging from one to five. Artefacts such as swords, daggers, arrowheads, triangular arrowheads, armour, and other related tools were unearthed predominantly in the eastern cemetery and the western catacombs.

Considering the finds, cultural characteristics, and the radiocarbon dates conducted on human samples, this cemetery was in use from the Parthian period through to the Islamic era.

During the three excavation seasons led by Dr. Sharifi, a total of 10 skeletons were examined and sampled by Zeinab Salehi and Motaharteh Amjadi for this study. A detailed account of the graves associated with these remains are summarised as below.

##### 9.2. Human samples

###### 9.2.1. Trench F-34. Catacomb 3

Grave 3 in trench F34 is marked by a narrow entrance that leads to a rectangular chamber, which is approximately 3 metres deep. Inside, the skeletal remains of a young female (genetically and anthropologically), estimated to be between 18-20-years-old, were found on the southern side (IRN02). The direct radiocarbon date (255-410 cal CE) (Supplementary Table S1) confirms the dating of the burial to the end of Parthian to the middle of the Sassanid Period. Adjacent to her, on the western side (IRN01), lay the disturbed remains of another female, approximately 20 to 22 years old, identified by the slightly more worn condition of her teeth, with visible third molars.

Interestingly, a silver earring associated with the second burial was found at the northern end of the grave. Additionally, the central area of the crypt contained a large, irregular pit with bone fragments and seven varied pottery vessels were located on the northern side. The dating of IRN01 is approximately 200 CE to 460 CE based on the archaeological context, corresponding to the end of Parthian to the middle of the Sassanid Period.

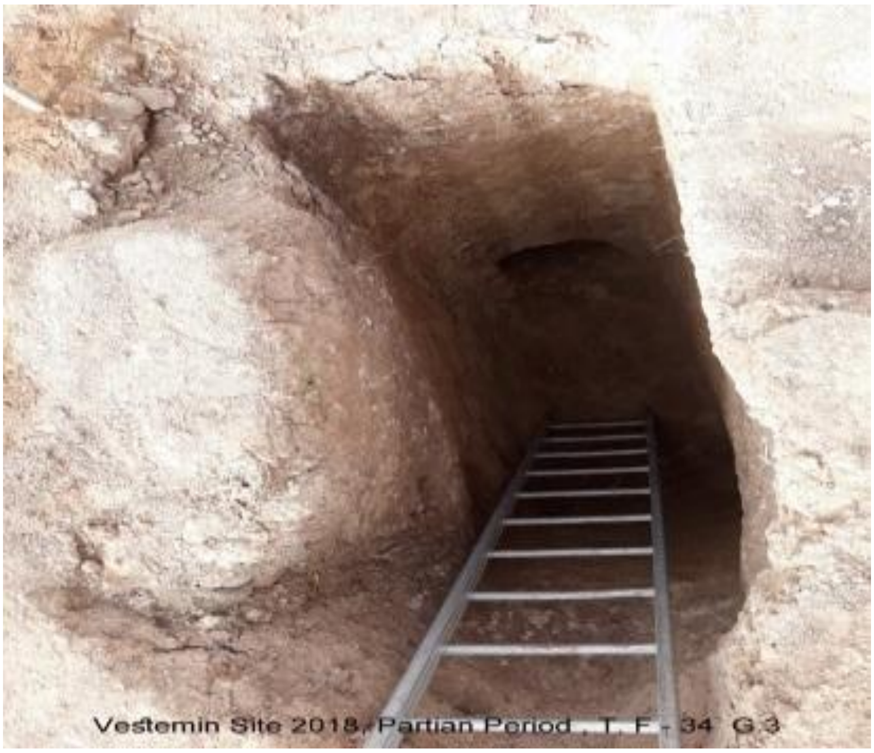


**Figure 9.2. The entrance of the Catacomb 3 (Photo by Abdolmotalleb Sharifi).**

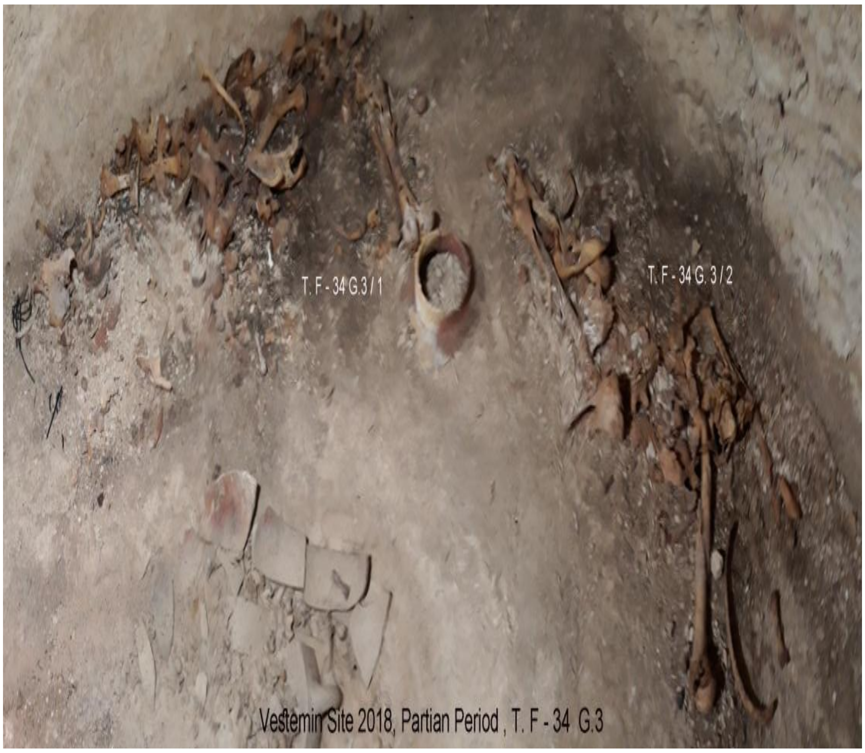


**Figure 9.3. The arrangement of the human skeletal fragments and grave goods within the interior space of Catacomb 3 (Photo by Abdolmotalleb Sharifi).**

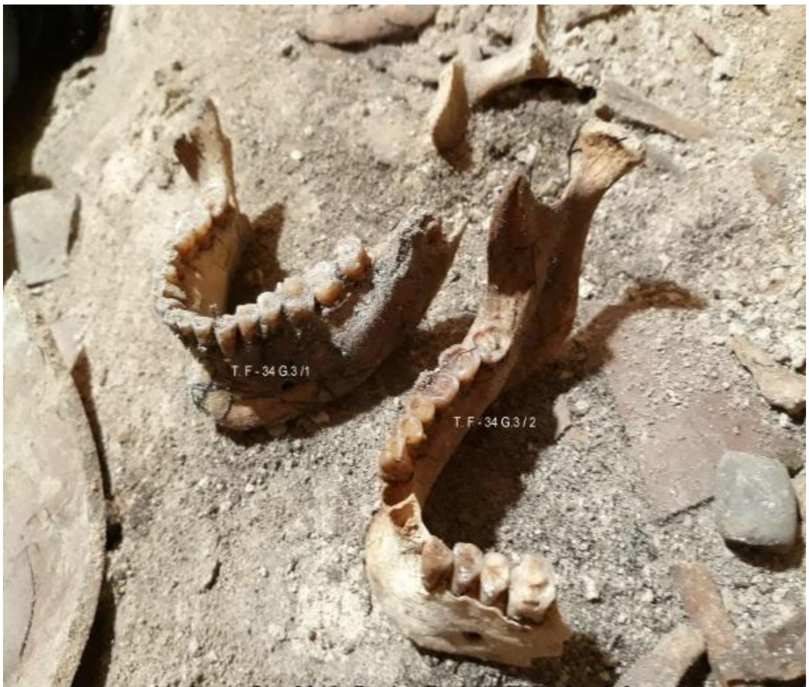


**Figure 9.4. Remains of mandibles of the skeletons associated with IRN01 and IRN02 (Photo by Abdolmotalleb Sharifi).**

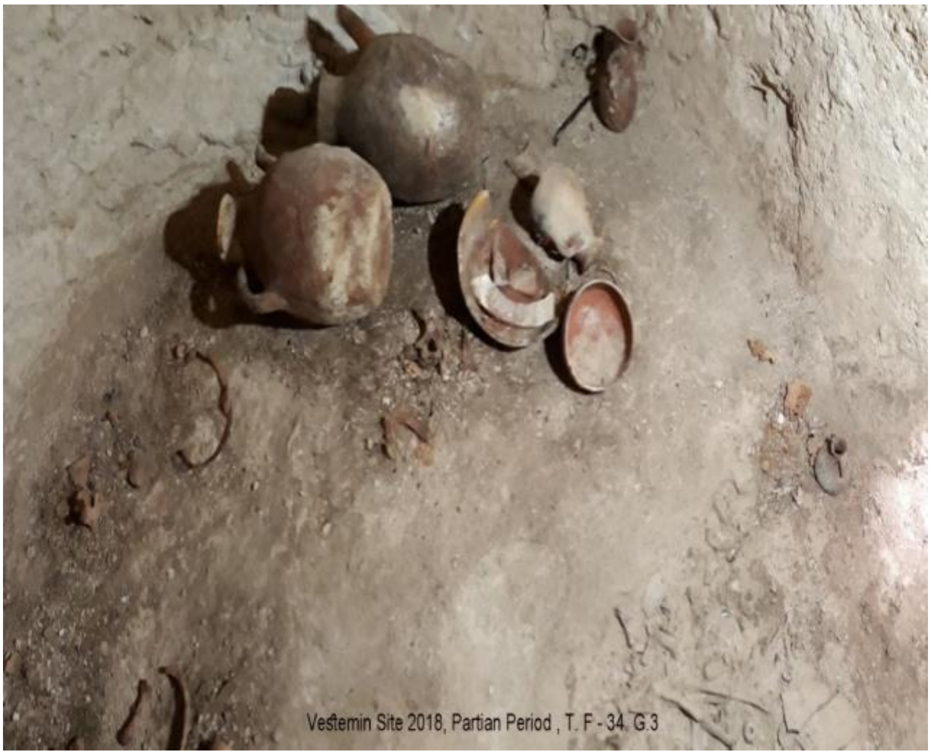


**Figure 9.5. The seven varied pottery vessels were located on the northern side of TF34G3 (Photo by Abdolmotalleb Sharifi).**

##### 9.2.2. Trench F-34 Catacomb 4

Grave 4 in trench F34 presents a different configuration, being over 3 metres deep with a narrow corridor leading to a circular chamber. In this grave, the remains of two individuals were discovered. The first (IRN03), a young female aged between 20 and 25, was found on the northern side. Her skull was destroyed, however, her orientation suggests a west-east position, lying on her right shoulder facing south. Decorative beads, including agate and lapis lazuli, were found around her hands and chest. The dating of this grave, based on archaeological context, is estimated to be between 200 CE and 460 CE, corresponding to the end of Parthian to the middle of the Sassanid Period.

The second individual, positioned on the southern side, was notably more delicate, with a skull so fragmented that it was unrecognisable. Above this individual, a fine miniature pottery vessel—possibly for medicinal oils or fragrances—was discovered, along with a bronze earring and two small rings near the facial area. This individual's hands displayed traces of four bracelets of varying sizes and various beads were scattered around and in front of the hands. Based on anatomical assessments, and the presence of deciduous teeth, the estimated age of this child (IRN15) is between 6 and 7 years. The dating of this grave based on the archaeological context is between 200 CE and 460 CE, corresponding to the end of Parthian to the middle of the Sassanid Period.

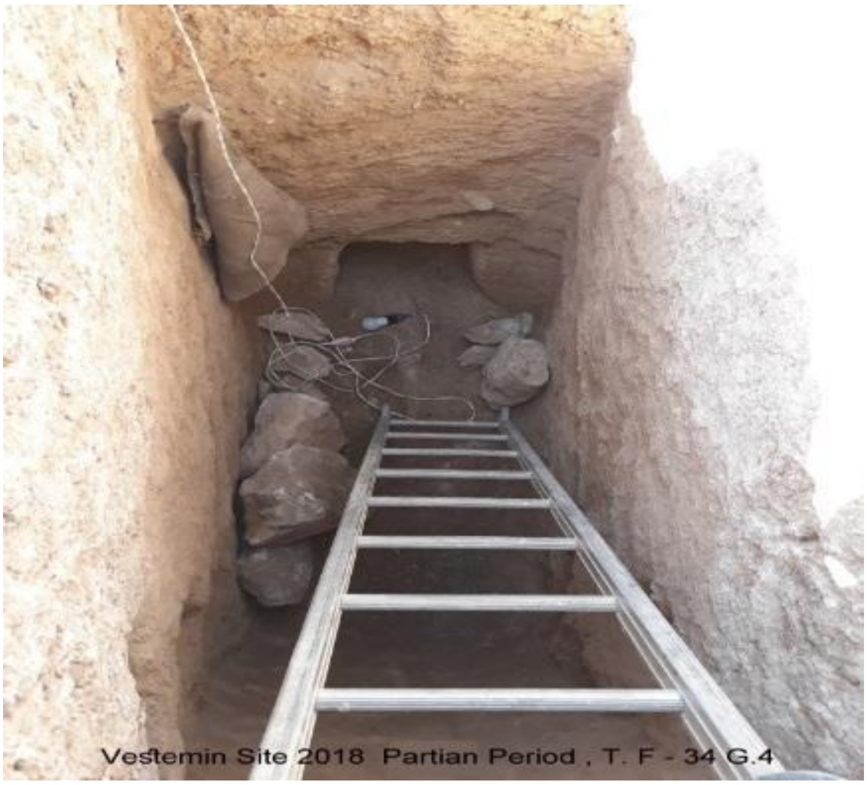


**Figure 9.6. The entrance of the Catacomb 4 (Photo by Abdolmotalleb Sharifi).**

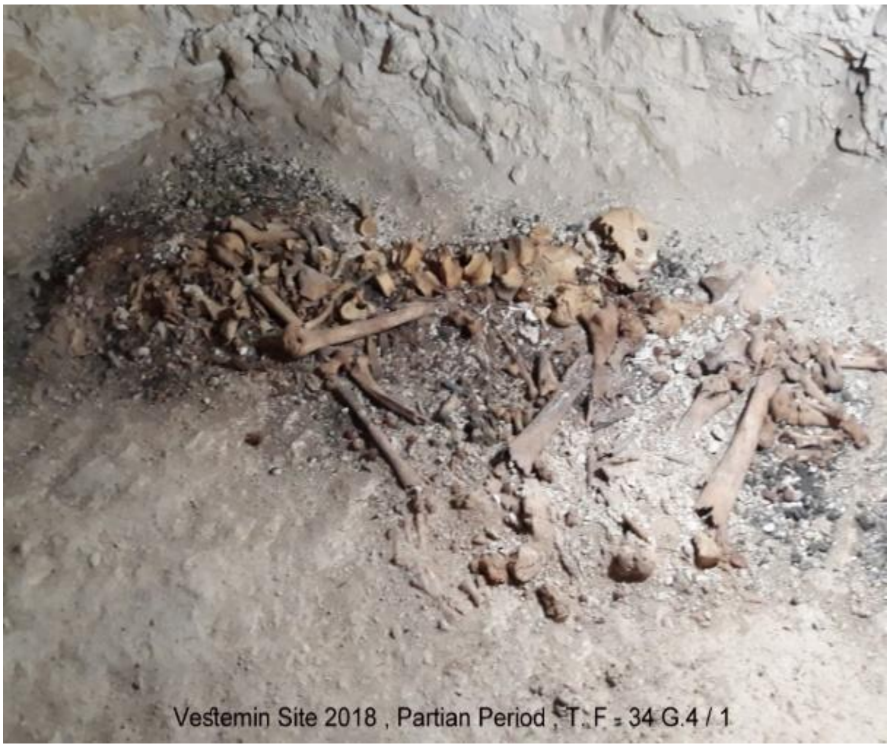


**Figure 9.7. The disturbed remains of two individuals within the interior space of Catacomb 4 (Photo by Abdolmotalleb Sharifi).**

Interestingly, between these two burials in grave 4, five pottery vessels were placed as offerings. A unique discovery for this historical period in northern Iran was the arrangement of 11 intact eggs alongside these vessels and on the ground. This find offers a rare glimpse into the burial customs and beliefs of the time.

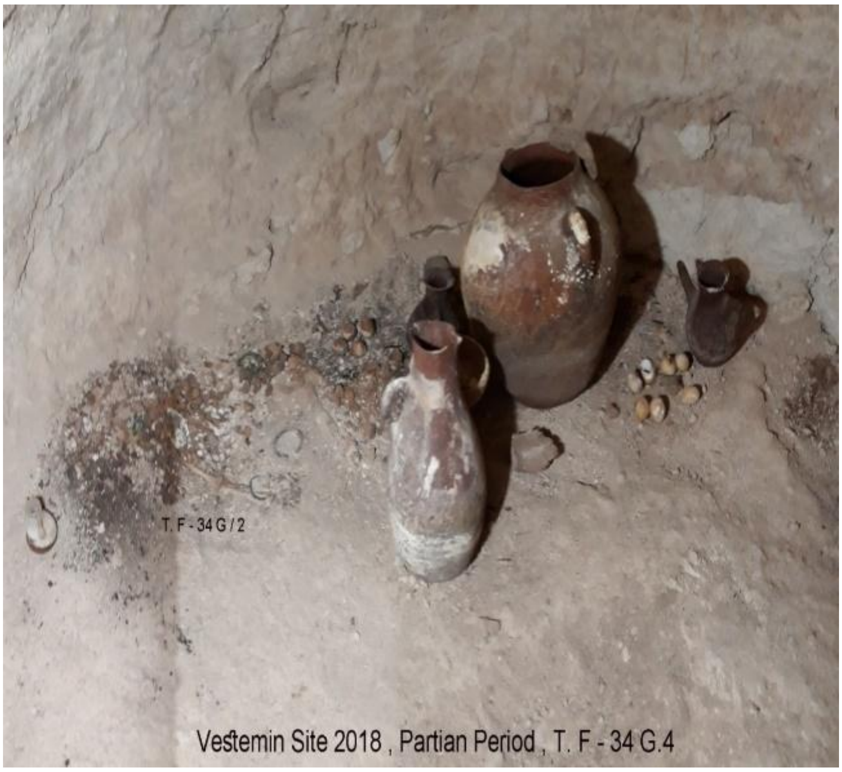

**Figure 9.8. The arrangement of 11 intact eggs alongside these vessels and on the ground within the interior area of Catacomb 4 (Photo by Abdolmotalleb Sharifi).**

##### 9.2.3. Trench I-39 Catacomb 2

In another grave, identified as T-I 39 G.2, a deep crypt with a collapsed roof and an entrance facing almost directly east was found. The corridor housed 10 stone pieces related to the entrance door. On the northern wall of the corridor, a niche revealed a small vessel alongside scant remains of a very young child based on the presence of deciduous teeth. Inside the crypt, on the southern side, lay an adult burial with a delicate skeleton—oriented east-west, legs bent and a relatively intact skull. Near the skull area, a delicate bronze object, possibly an earring, was discovered. This individual (IRN77) was identified as female and estimated to be between 20 and 25 years old. On the northern side of the catacomb, the skeleton of a middle-aged man was positioned southwest-northeast, lying on his right shoulder and facing southeast. In front of the face, a large iron dagger was found. The dating of this grave based on the archaeological context is between 200 CE and 460 CE, corresponding to the end of Parthian to the middle of the Sassanid Period.

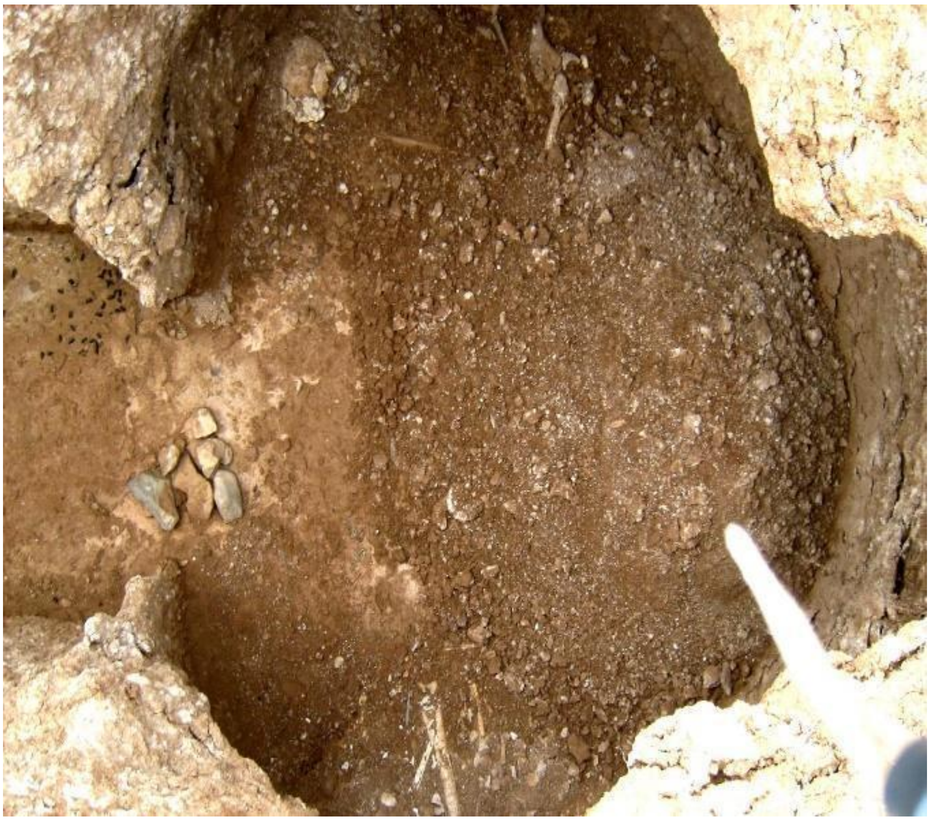

**Figure 9.10. Crypt grave in Trench I-39, Catacomb 2 - traces of two burials within the crypt and one burial in a niche of the entrance corridor (Photo by Abdolmotalleb Sharifi).**

##### 9.2.4. Trench H34, Catacomb 6

The grave identified as T-H 34 G.6 presents a unique arrangement within its entrance corridor. The corridor, oriented southeast at approximately 105 degrees, revealed two distinct skeletal remains forming a bone mass, positioned opposite the entrance door and intertwined with a large stone mass. Interestingly, one-third of the eastern corridor was completely empty. Upon examination, the first individual (IRN 78), a female aged between 20 and 25 years, was discovered with a dolichocephalic skull and signs of cranial swelling. Alongside her were two arrowheads and two small pots. This grave is dated based on the archaeological context between 200 CE and 460 CE, corresponding to the end of Parthian to the middle of the Sassanid Period.

Beneath this mass, a second individual—not included in further genetic examination—was a male estimated to have died between 25 and 30 years old. Both sets of remains, based on the arrangement and evidence, are considered secondary burials.

Furthermore, an intriguing aspect of T-H 34 G.6 is the positioning of the stone mass related to the entrance door, indicating a significant alteration to the burial structure over time. This discovery suggests a complex burial practice, possibly involving multiple stages of rituals.

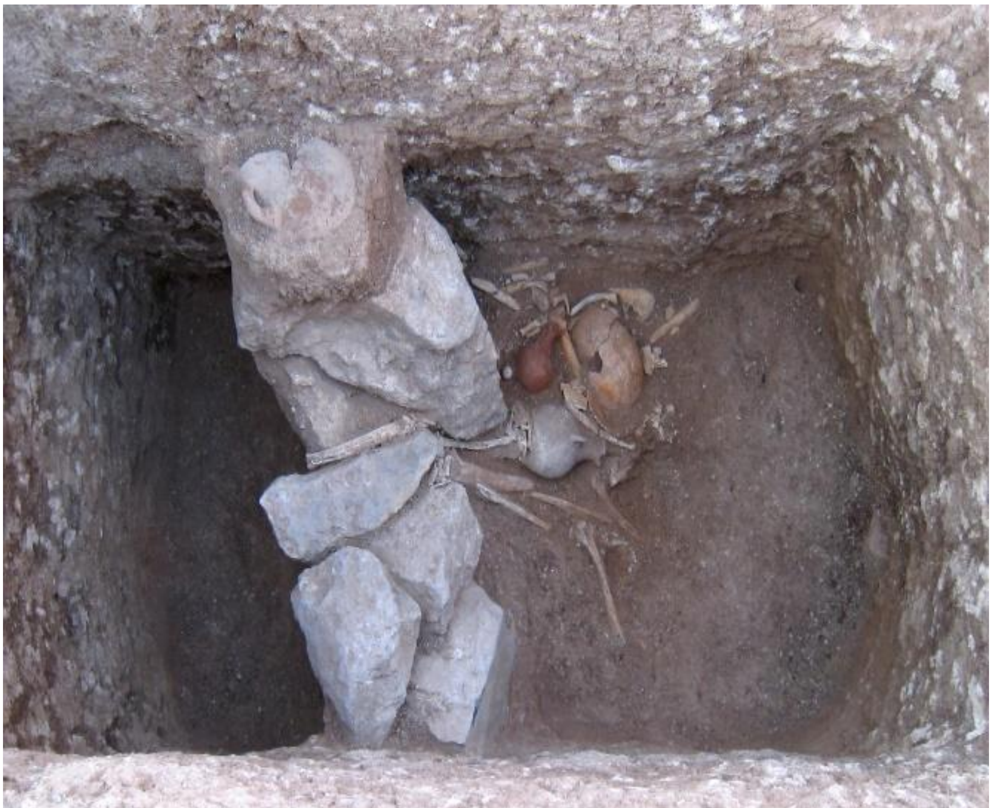

**Figure 9.11. The remains of two secondary burials in the entrance corridor of Catacomb 6 (Photo by Abdolmotalleb Sharifi).**

##### 9.2.5. Trench H-34, Catacomb 4 (TH34G4)

In another area of the catacomb, the disturbed remains of three adults were discovered. These individuals were positioned on different sides of the grave – south, west (in front of the entrance) and north. Analysis of the skeletal features, such as the mastoid process and sciatic notch, suggests the first individual (IRN74), a female (genetically and anthropologically) aged between 65 and 70 years, was positioned on the south side. The second (IRN76), likely a female aged between 20 and 22 years, was found on the west side, with evidence of a dolichocephalic skull. The direct radiocarbon date (255-415 cal CE) (Supplementary Table S1) confirms the dating of the IRN 74 to the end of Parthian till middle Sassanid period. The dating of IRN76, based on archaeological context, is estimated to be between 200 CE and 460 CE, corresponding to the late Parthian to the middle Sassanid period.

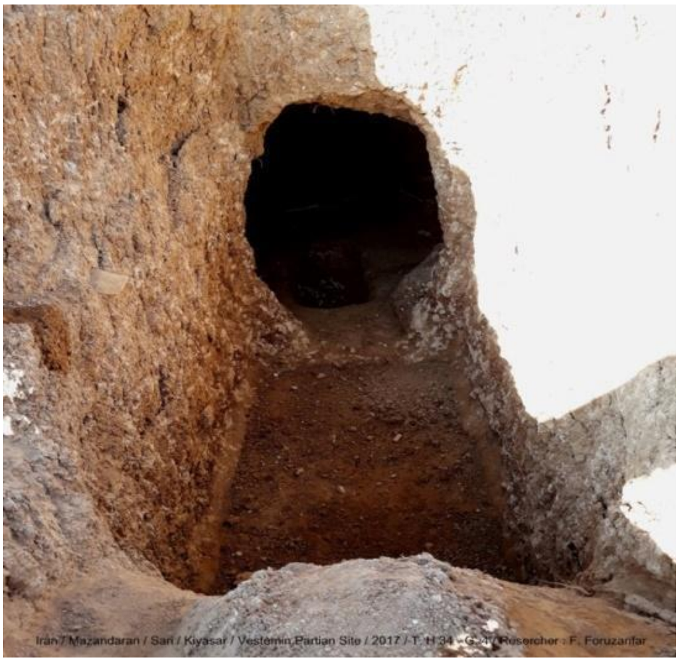


**Figure 9.12. The entrance to the catacomb and the rectangular pit in front of it (Photo by Abdolmotalleb Sharifi).**

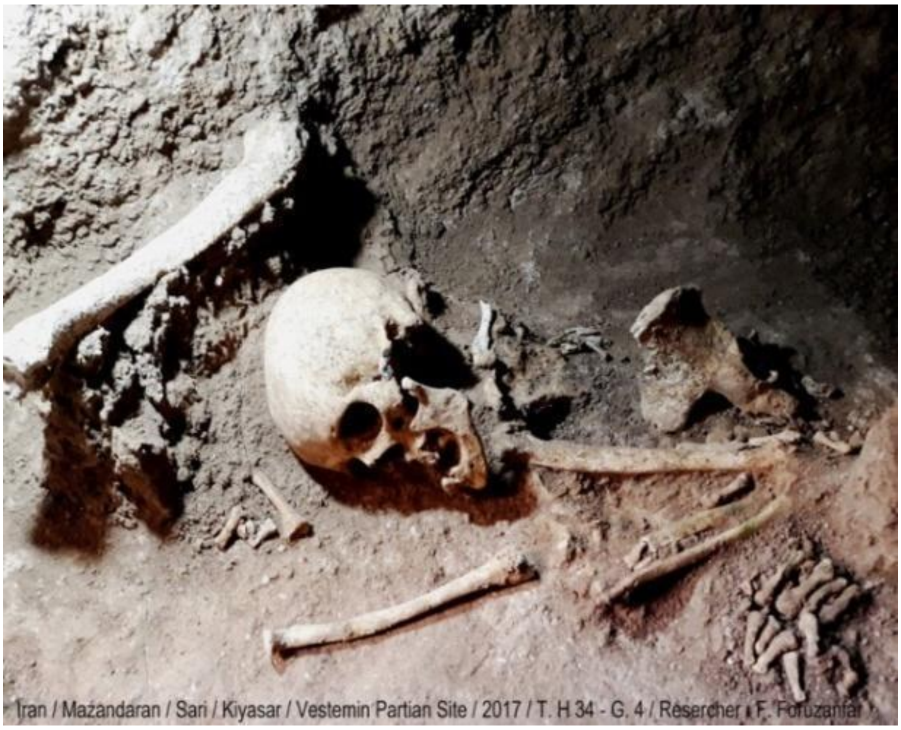


**Figure 9.13. The human skeletal remains of IRN74 in the south position within Catacomb 4 (Photo by Abdolmotalleb Sharifi).**

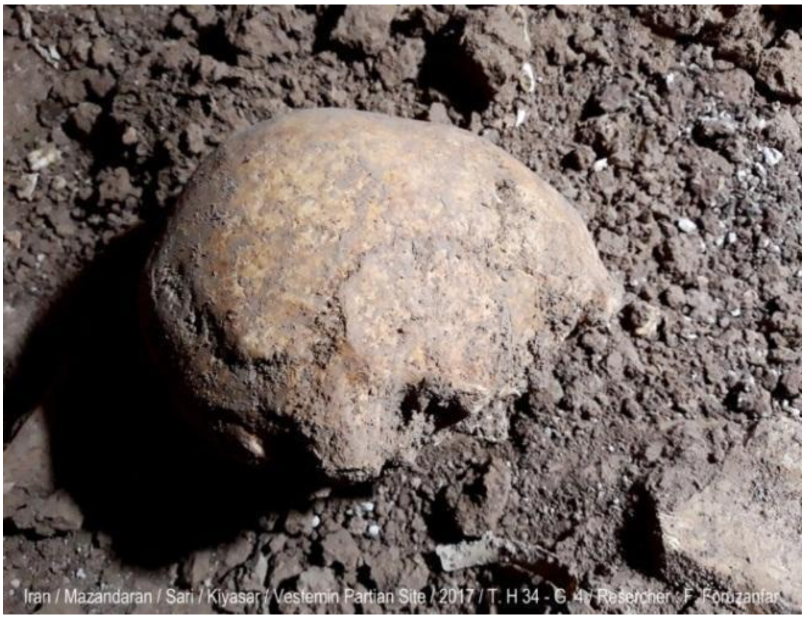


**Figure 9.14. The human skeletal remains of IRN76 in the west position within Catacomb 4 (Photo by Abdolmotalleb Sharifi).**

The third burial (IRN75) was located directly next to the entrance door on the north side. Many disturbances have been observed in this person's body, however the skull remained in its original position on the left shoulder. The individual was likely male and died at the age of 20-22. Based on the anatomical observations, (IRN75) and (IRN76) might be twins. This assumption is supported by a genetic result, where both individuals have the mtDNA haplogroup of H13a2b3. This individual is dated based on the archaeological context between 200 CE and 460 CE.

The discovered sediments in this tomb were mostly related to the collapse of the roof. The collapse of the roof has caused an apparent increase in the height of the grave. Repeated burials have also taken place at different times.

According to the evidence, it can be stated that the first burial took place on the south side and the second burial took place on the north side. After a collapse of the roof on top of them, the third burial on the west side was established.

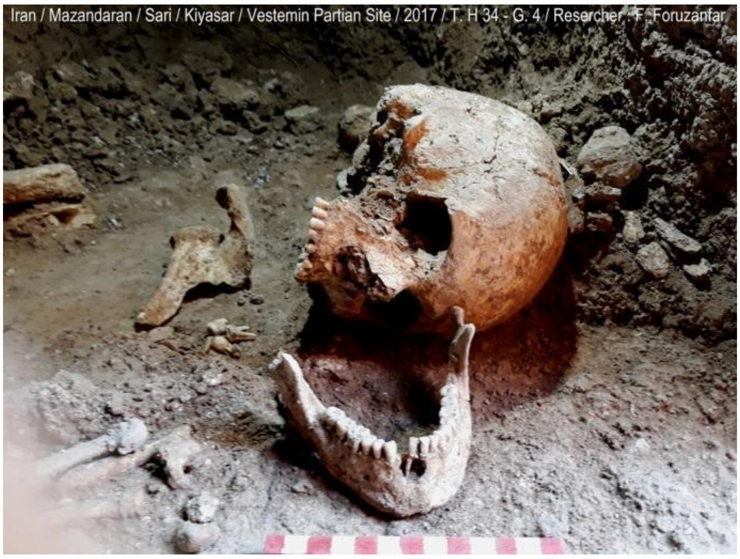


**Figure 9.15. The human skeletal remains of IRN75 in the north position within Catacomb 4 (Photo by Abdolmotalleb Sharifi).**

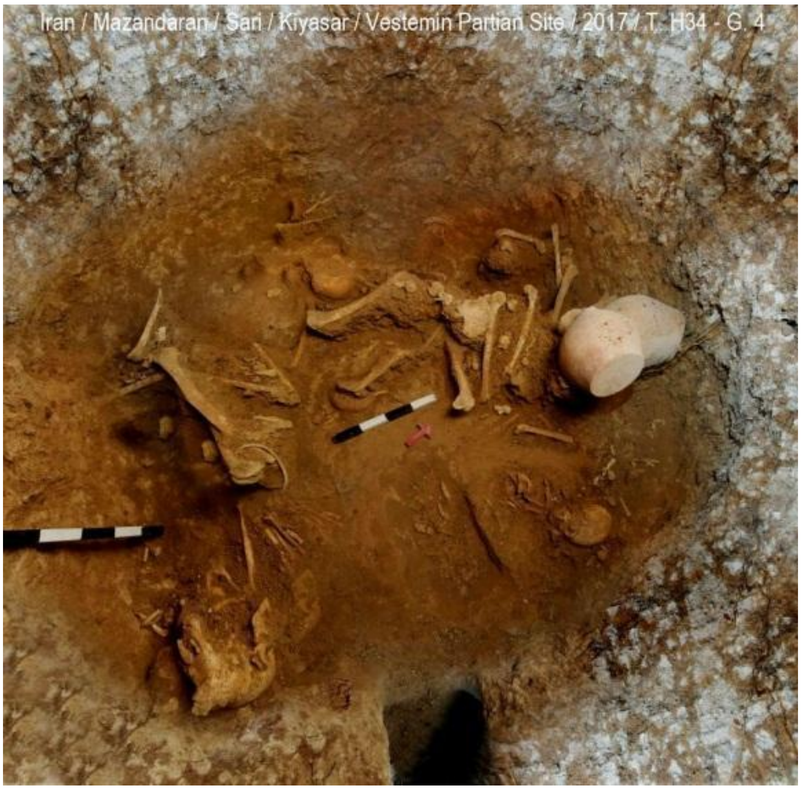


**Figure 9.16. Top view from TH34G4, showing the positions of three burials within it (Photo by Abdolmotalleb Sharifi).**

### Supplementary information to the genetic analyses

#### Chapter 10: Reconciling genetics and archaeology from a uniparental perspective

Our Historical-period mtDNA dataset from the north resembles the prehistoric and modern Iranian and South-Central Asian groups in many aspects. For example, the oldest sample from the Mesolithic Belt Cave (Narasimhan et al., 2019; Lazaridis et al., 2016) in northern Iran belongs to haplogroup HV, which has a frequency of 30% in today's northern Iran (Amjadi et al., 2024). Likewise, in our sample set, this haplogroup is detected in over 10% in the Historical period samples and reappears in our Medieval sample from the Kalmakareh cave as well.

The most frequent mtDNA sub-haplogroups, such as T2, R2, and J1 in the ancient Iranian populations can be identified in the current research from Chalcolithic to historical times (Supplementary Table S5). The new T2c sample from Chalcolithic Gol Afshan Tepe in Isfahan province, dating back to 4690-4360 cal BCE, can be recognized as the second oldest representative of this haplogroup from the Iranian Plateau, following a sample from the late 9th - early 8th millennium BCE Tepe-Abdul Hosein (Broushaki et al., 2016). Another frequent mtDNA haplogroup, R2, is specific to West Eurasia and is considered to have originated between 21,000 and 31,000 years ago, likely in southern Iran during the pre-Last Glacial Maximum (Derenko et al., 2013). R2 is described in the majority of Neolithic sites from Iran, such as in Ganj Dareh (Narasimhan et al., 2019; Lazaridis et al., 2016), Tepe Abdul Hosein and Tepe Guran (all from central Zagros area) (Broushaki et al., 2016; Allentoft et al., 2024, see details in Supplementary Table S5). T2 and R2 are present in a maximum of 21% of modern Iranian ethnic groups in total (Amjadi et al., 2024), suggesting a detectable long-term maternal genetic continuity across the region (Supplementary Table S6).

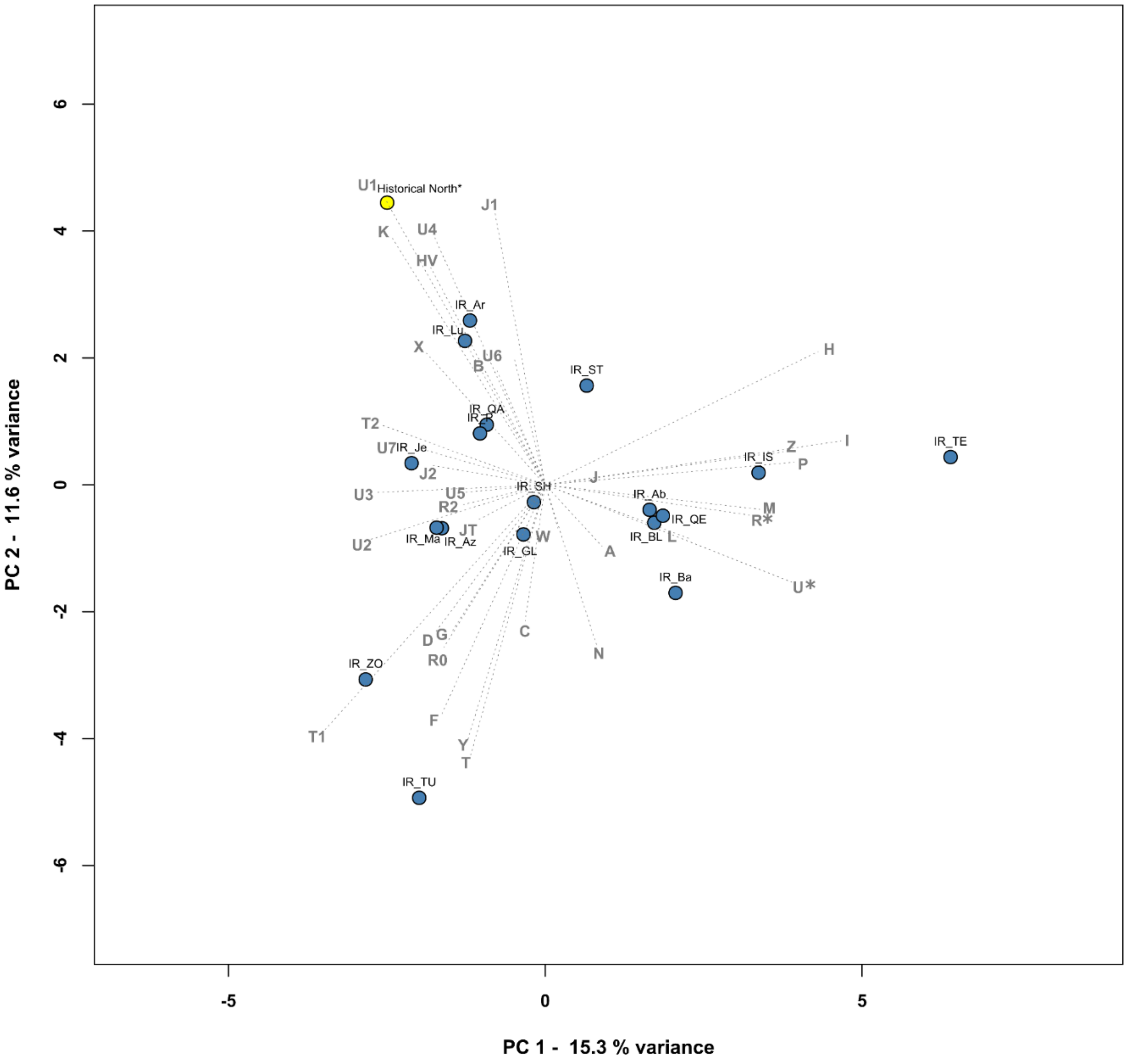


**Fig. 10.1. PCA plot based on mtDNA haplogroup frequencies in modern Iranian and selected ancient Iranian groups.** The frequencies of 36 haplogroups were used in modern and Historical North-Iranian groups. The first two principal components (PCs) capture 26.9% of the total variance (see Supplementary Table S6 for details and abbreviations). Some of the modern Iranian samples align with our historical samples along PC2, particularly the Iranian Armenians and Lurs who have similar frequencies in haplogroups H and HV, and elevated components of K and J1.

As shown by Supplementary Figure 10.1, The historical period mtDNA gene pool consists of a wide range of haplogroups (HV, J1, K, U1, U2, U4, W) from Marsin Chal (HV, J1, U1 and X), and Liarsangbon and Vestemin with H, K and T2 respectively. We combined all three northern sites into a single group, referred to as 'Historical North'. When considering the individual combination of historical samples with modern Iranian groups along PC2—via shared proportions of haplogroups K, J1, and U1—the Historical North group clusters closely with Iranian Armenians and Iranian Lurs. Notably, along the same PC2 axis, other Iranian groups such as Iranian Bakhtiari and Iranian Persian also form distinct clusters in proximity to these populations. Given the diversity of the maternal historical sample set, and considering the lack of significant differences between Iranian Lurs and Bakhtiari, as well as the high maternal similarity among Lurs, Bakhtiari, Armenians, and Persians in Iran, we underscore the strong local maternal genetic continuity from historical times to the modern era.

The Indo-Iranian borderlands (Supplementary Figure 10.2) as a part of the Iranian Plateau encompass parts of southeast Iran mainly Baluchistan area and Sistan. This region includes the Bampur Valley (Wright, 1984; Gorgi et al., 2021), Kerman province, Iranian Sistan, southern Afghanistan, Pakistani Baluchistan, and other areas of Iranian Balochistan around the fifth to the end third millennium BCE (Hiebert & Lamberg-Karlovsky, 1992; Gorgi et al., 2021). The communication channels within this region had a significant role in creating a number of cultural similarities among the communities, detected through archaeological records. Kech-Makran region geographically is the south-easternmost part of the Iranian Plateau located in southwestern Pakistan (Hiebert & Lamberg-Karlovsky, 1992; Gorgi et al., 2021).

The female burial practice of Shahi Tump archaeological site in the Kech-Makran region had parallels to Shahr-i-Sokhta in Sistan and Baluchistan province of Iran, Gonur Tepe in Turkmenistan and Sappali Tepe in Uzbekistan. Additionally, a shared characteristic of Shahr-i-Sokhta, Gonur Tepe, and Shahi Tump is the usage of metal seals as grave goods, predominantly associated with female burials (Salvatori & Tosi 2001; Jarrige et al., 2011; Sheikhi et al., 2023). All these populations exhibit a notably similar distribution of maternal haplogroup frequencies in M, HV, J1, T1, U1 and W (Figure 2A, Supplementary Table S6). These female burial practices, which were shaped in the early phase of the BA time horizon in the Indo-Iranian borderland areas, continued in the later phase of the BA, at the sites of the South Central Asian regions such as Gonur in Turkmenistan and Bustan in Uzbekistan, which became later integral parts of the Bactria Margiana Archaeological Complex (BMAC).

Notably, two genetic groups from BA Shahr-i Sokhta are the only ancient groups in our analysis with haplogroup M and also exhibit elevated frequencies of W and R*. The Shahr-i Sokhta genetic group 2 (ShahrISokhta_BA2) was previously described as part of the Indus Periphery Cline (IPC), due to the shared AHG-related ancestry with South Asian genomes, and the lack of ANF ancestry in a similar pattern to the BA Turkmenistan Gonur group 2 (Figures 3-4) (Shinde et al., 2019; Narasimhan et al., 2019). These genome-wide trends were reproduced in the current study, along with BA2 groups genetic connection to Gonur_BA2 from Turkmenistan.

Despite the genetic differences between the BA1 (n=12), BA2 (n=8), and BA3 (n=1) groups from this site, most of the archaeological and chronological contexts of these individuals are similar. On the other hand, from the total of the 21 individuals of Shahr-i Sokhta, only three graves from the genetic group BA2 had artifacts linked to the burial customs of the Harappan civilization. The mtDNA data indicate continuity with modern Iranians, such as the Baloch people from southeastern Iran, who also carry these haplogroups (Amjadi et al., 2024). This observation corresponds with the comparable geographical location of Shahr-i-Sokhta in Sistan and Baluchistan province. However, the signs of population movement from the Shahr-i-Sokhta toward the South Central Asian regions started by the initial decline of this cemetery at Period IV, phase 0 (Sarhaddi-Dadian et al., 2019). This is in accordance with the formation of the BMAC in the South Central Asian areas, where the genetic makeup of the BMAC also affected the BA non-BMAC sites in South Central Asia, such as Sarazm in Tajikistan. These areas later functioned as hot spot zones in the Indian or Indo-Iranian cline and population movement toward the North and South Indian subcontinent (Kerdoncuff et al., 2024).

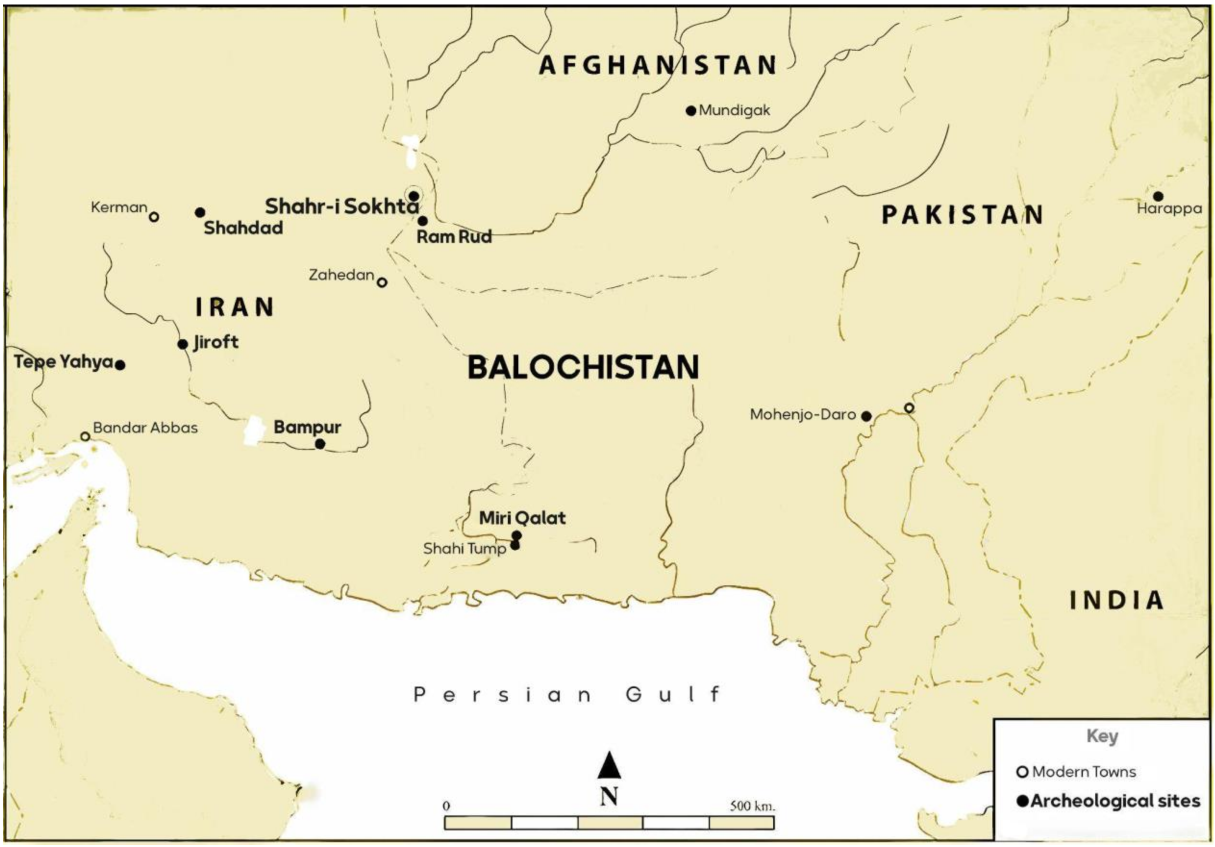


**Fig. 10.2. The map of Indo-Iranian borderland territory.**

Counterposed on the PC1 axis of Figure 2B, and with a distinction on PC2, the EN samples from Ganj Dareh are characterized by pronounced frequencies of haplogroups R2, U7, and HV. In a similar direction in the PC1-PC2 space, our historical samples show continuity in many maternal lineages from the prehistoric times. They cluster together with northeastern Iranian ancient samples from Chalcolithic Tepe Hissar, northwestern samples from IA Hasanlu, and BA site Gonur from Turkmenistan on the mtDNA PCA.

**Novel Y-chromosomal sub-lineages in ancient Iran**

In the published Neolithic Y-chromosomal datasets, R2 has, to our knowledge, been exclusively found in the Early Neolithic (EN) samples from Ganj Dareh. Our study, however, demonstrates its Chalcolithic occurrence at Gol Afshan Tepe. According to Family Tree DNA Discover (FTDNA Discover) (Gene, 2022) and YFull (https://www.yfull.com), two databases including modern and ancient genomes, subgroups of the R2a2b1 (R-FGC12582 or R-Y8763, https://www.yfull.com/tree/R-Y8763/) branch, identified in Gol Afshan Tepe has spread to distant regions. This lineage has been discovered in Chalcolithic and BA South-Central Asia, IA Pakistan, Medieval India and also Medieval Sudan. Furthermore, in these databases, it is also commonly found in a vast region between modern Italy and Singapore. This suggests the spread of this haplogroup is in correlation with the movements of the Neolithic Iranian farmers (Narasimhan et al., 2019; Lazaridis et al., 2016).

The macro-haplogroups J1 and J2 originate from the western parts of the Persian Plateau and its surroundings, where they formed approximately 20,000 and 31,600 years ago, respectively (Singh et al., 2016; Sahakyan et al., 2021). We report their sub-lineages in the Persian Plateau, which sheds light on their persistence in the region, leading to their discovery in the historical groups. We discovered two rare Y-DNA haplogroups, J1a2a2a~ in Liarsangbon (J-FGC6141 or J-FGC6031, https://www.yfull.com/tree/J-FGC6031/) and J2b2a2a~ in Marsin Chal (J-FGC61903 or J-Y28235, https://www.yfull.com/tree/J-Y28235/) (for more information see Supplementary Table S4). These two sub-lineages have formed around the Neolithic period (mean 9100-8100 and 7200-6500 years BP respectively, taken from the calculations in FTDNA Discover and YFull), however they are not common in the mentioned databases. To our knowledge, previously published ancient samples either did not trace downstream enough on the phylogenetic tree or belonged to different or sister branches compared to the two J sub-haplogroups identified here. Given the presence of these sub-haplogroups in modern Middle Eastern populations within the same databases, the limited detection in ancient samples is likely a result of the relatively small number of published ancient genomes from the region.

Y-chromosomal haplogroup T, found at variable frequencies across West Asia, Africa, and Europe with generally rare frequencies. It emerged approximately between 25,000 and 15,000 years ago, during the late glacial period, probably in the region of the Iranian Plateau (Terreros et al., 2011). It only appears in the Chalcolithic period in our dataset, representing about 7% of the Chalcolithic population within the Iranian Plateau, and absent in both earlier and later periods (Figure 2C).

#### Connections of the Caucasus and the Iranian Plateau

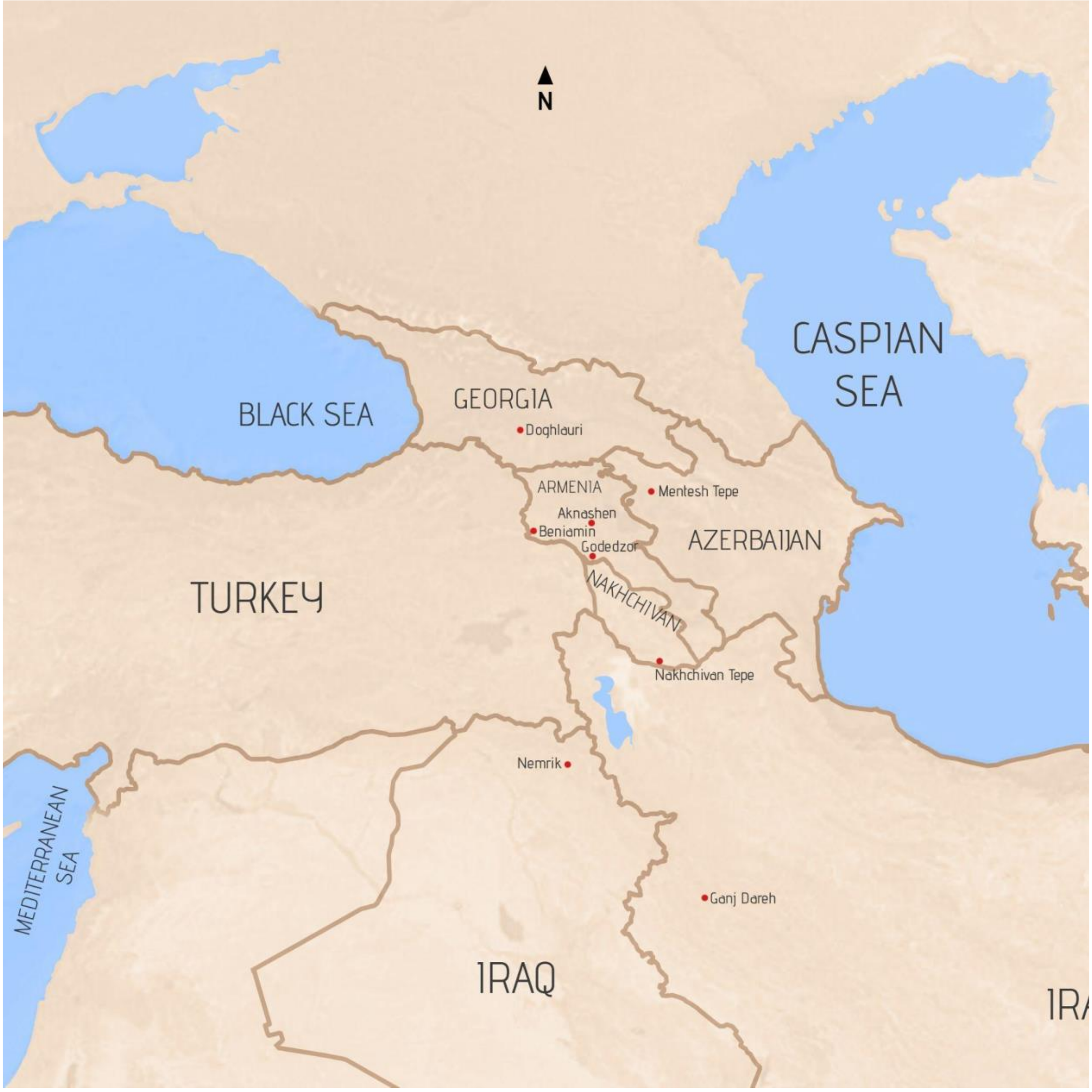


**Figure 10.3. A map of the Caucasus area with various archaeological sites, used in comparison with the sites of this study.**

The pattern of genetic continuity and homogenization is also observed in the Neolithic to Iron Age Caucasus cluster on the genomic PCA. Individuals associated with the Kura–Araxes culture in Georgia (Koptekin et al., 2023) fall close in PCA space and ADMIXTURE with published Kura–Araxes individuals from Armenia (Lazaridis et al., 2022) (Figures 3A and 3B).

During the Late Neolithic, the Zagros pastoralists expanded to the South Caucasus region, detected at Nakhchivan Tepe, a flat site on the right bank of the Nakhchivançay River–close to the modern city of Nakhchivan, in the border of present day of Iran and Azerbaijan (Palumbi et al., 2021; Amjadi et al., 2024) (Supplementary Figure 10.3). The notable genetic and cultural effect of Zagros pastoralists remained stable during the Chalcolithic in South Caucasus, detected in the Southern part of Armenia, at the Godedzor archaeological site (Margaryan et al., 2017). However, by the time of the Early Bronze Age, Kura–Araxes groups expanded into today's Iran (Maziar, 2020). These back and forth population movements might have formed the genetic homogeneity, shown by the close location of BA-IA Armenian ancient and modern groups to most of the ancient and modern Iranian maternal gene pool on the mtDNA and genomic PCA (Figures 2B, 3A and Supplementary Figure 11.2.) (Amjadi et al., 2024).

#### Chapter 11: Supplementary Figures and notes to the whole-genome based analyses

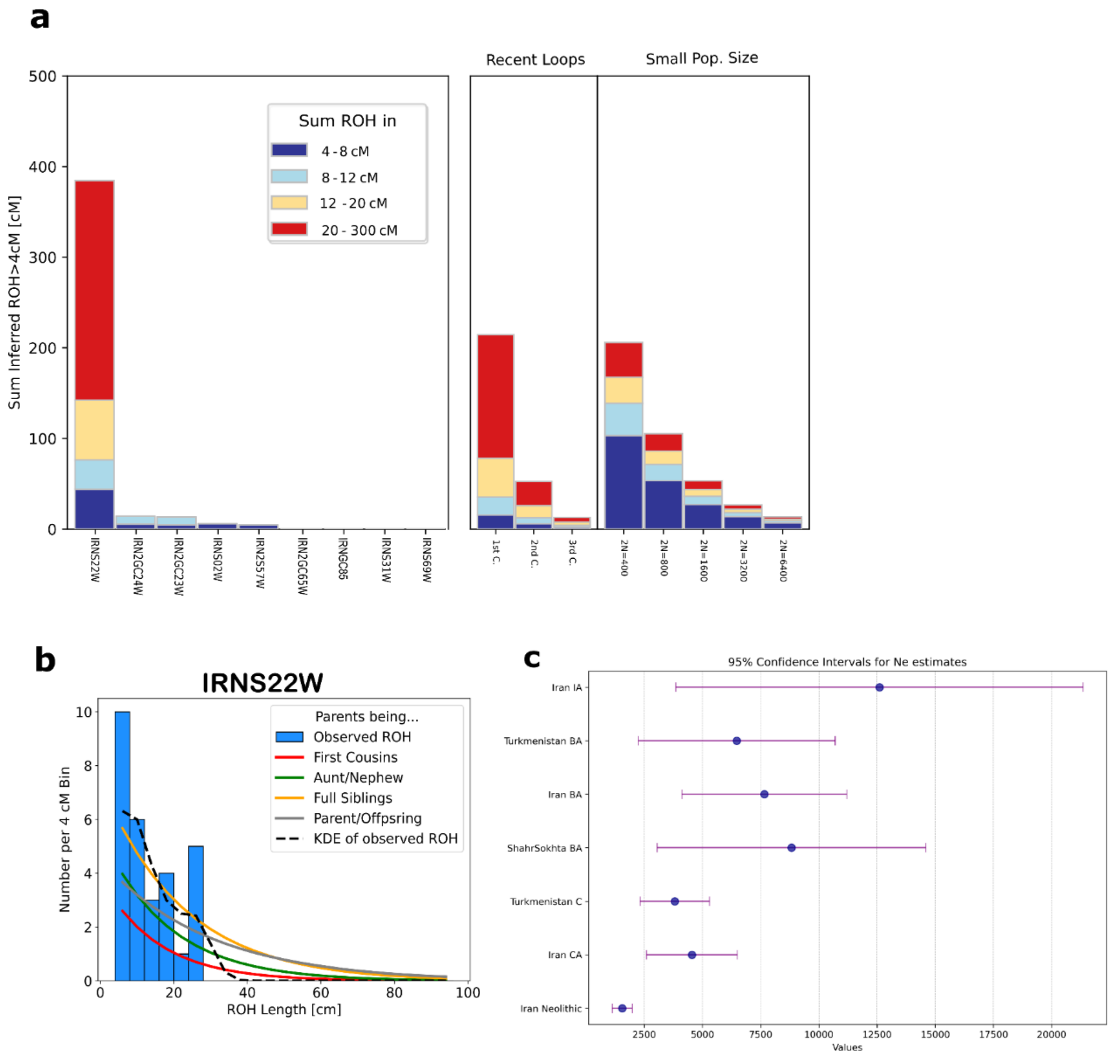


**Figure 11.1. Depicting homozygous runs in the genomes with hapROH that are indicative of small population size or parental relatedness. A**: multiplot of the newly analysed individuals, where only one sample shows signs of consanguinity and four have signals of smaller population sizes. **B**: ROH distribution in sample IRN22 (IRNS22W) from the Parthian period Liarsangbon, shows the signals of close parental relatedness. The Kernel Density Estimation (KDE) curve represents the probability density function of the observed ROH segment lengths for IRNS22W. **C**: Plot of effective population estimates in different prehistoric periods and groups within and around the Iranian Plateau. Detailed information can be found in Supplementary Table S15.

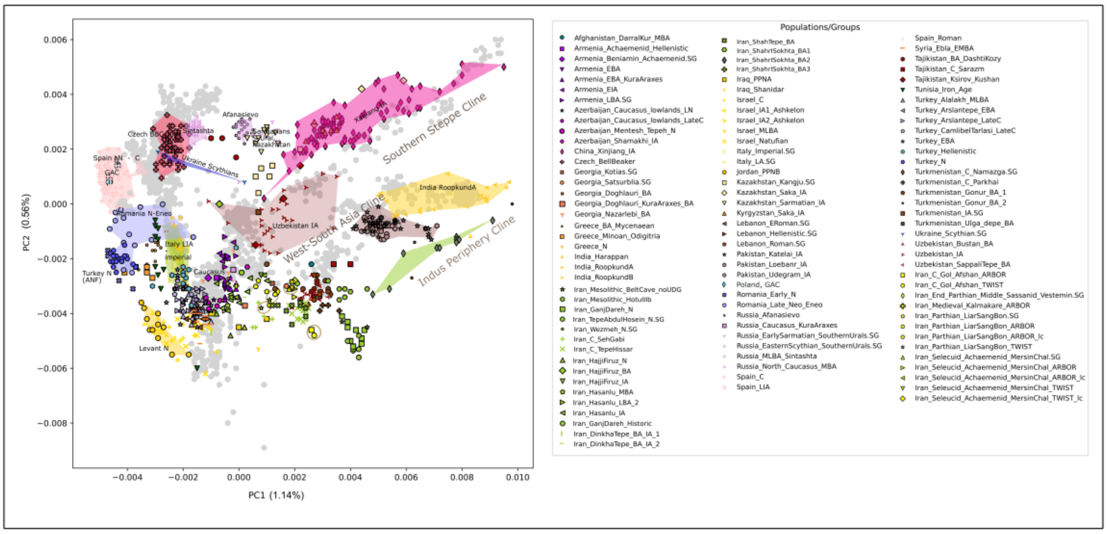


**Figure 11.2. Eurasian PCA focusing on the genetic variation of Western Asia and its adjacent territories.** Gray dots indicate modern reference populations, as in Figure 3. Olive circles indicate identical samples processed with different capture methods (see Methods and Supplementary Table S1-S4).

As shown on Supplementary Figure 11.2, the highlighted groups from Kazakhstan-Xinjiang IA, LIA-Imperial Italy and the Acheamenid-Parthian sample set, give a broader perspective on the genetic diversity associated with trade along the Silk Road and the cross-cultural interactions between major Persian empires, the Roman Empire, and Chinese populations. The genetic profiles are significantly distinct from one another, illustrating the varied genetic impacts of historical trade and interactions.

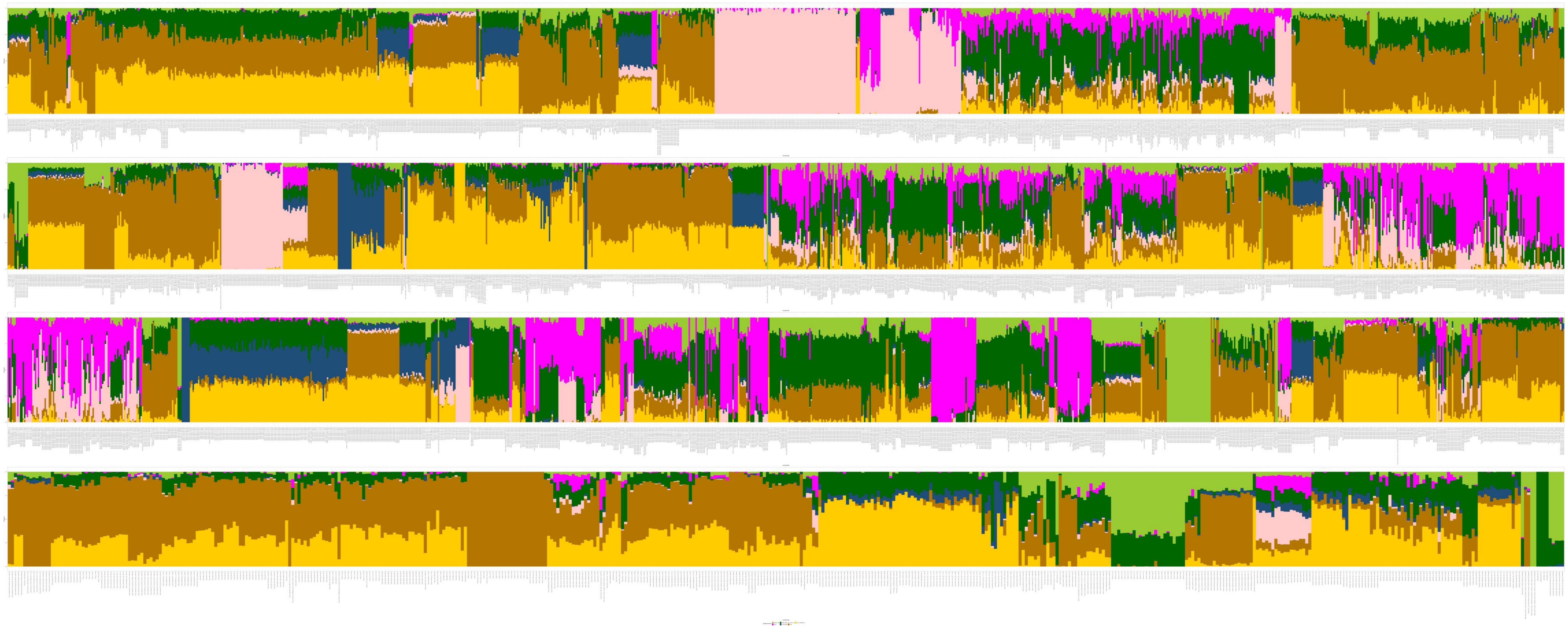

**Figure 11.3. Supervised ADMIXTURE plot with K=7, including every sample in the run.** Colours are defined as in the main text, on Figure 3.

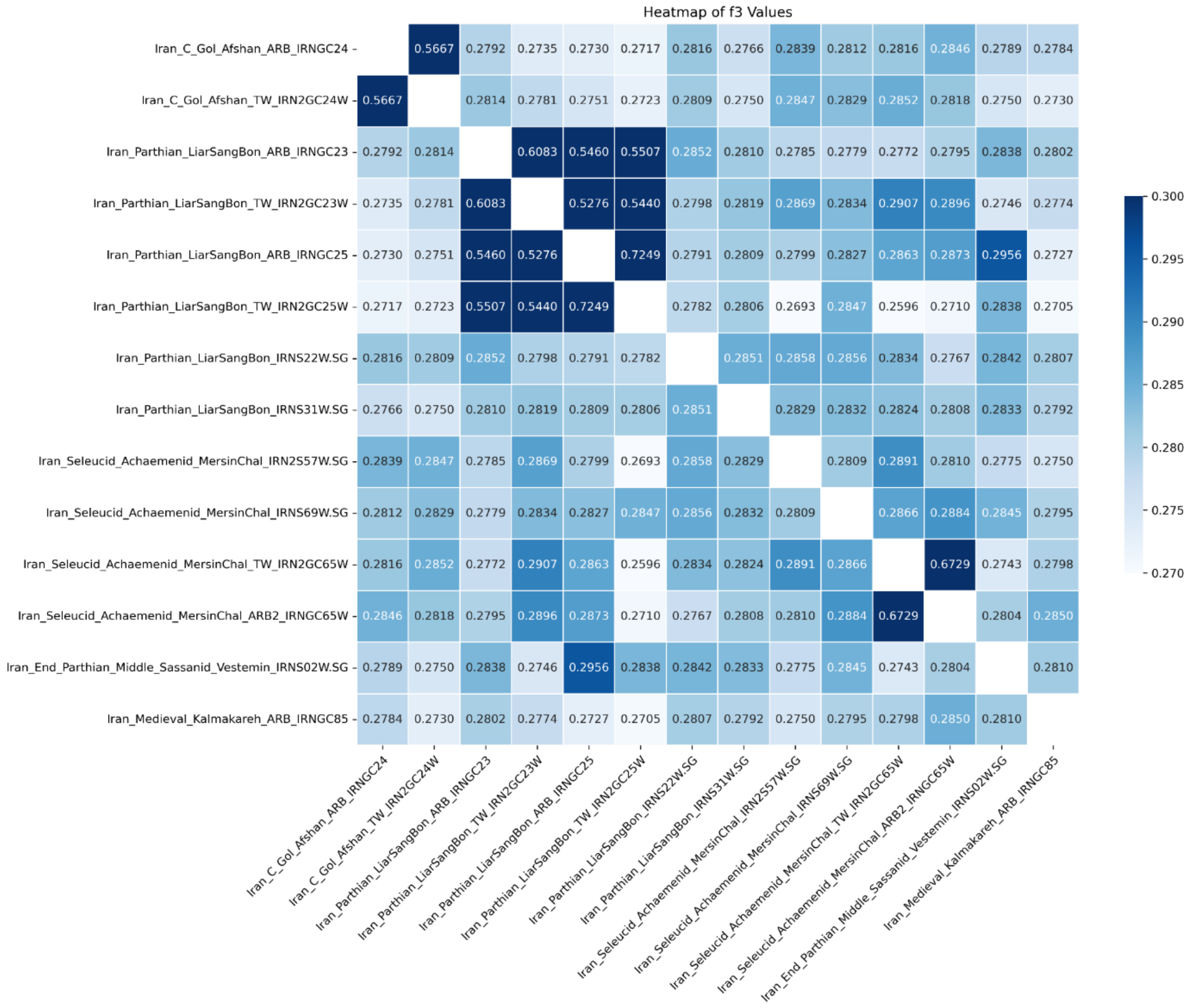


**Figure 11.4. Heatmap depicting the magnitude of f3 values between pairs of newly analysed historical samples from Iran.** Analysis includes samples from Kalmakareh, Liarsangbon, Marsin Chal, and Vestemin sites (Supplementary Table S12). F3 statistics were calculated in the form of f3(Test1, Test2, Mbuti.DG). Darker shades of blue indicate stronger genetic affinity between samples, while lighter shades mean greater genetic divergence. Standard deviations and Z scores are provided in Supplementary Table S12. This matrix shows the pairwise affinities of the individuals, prepared with different capture and sequencing approaches in this study. ARB stands for Daicel Arbor 'Prime Plus' capture type, TW for Twist 'Ancient Human DNA Panel Workflow' capture, and SG for shotgun sequencing in the sample names. The highest f3 values were observed within sites, particularly between the same individuals from the same sites in Iran prepared with different captures (Arbor and Twist).

While Arbor-captured samples tend to share more alleles with other Arbor-captured samples compared to their Twist counterparts (such as the Arbor-captured Medieval Kalmakareh sample), the difference is not significant. Using also the standard errors, the weighted t-test results in a t-statistic of approximately 1.19 and a p-value of approximately 0.32. This indicates that there is no statistically significant difference between the affinities of the two lists, Arbor and Twist, to the Kalmakareh Arbor sample, at the 0.05 significance level. A Linear Mixed Model analysis also shows no statistically significant difference between the Arbor and Twist methods when accounting for variability within the samples. The p-value for the method effect (0.238) suggests that the observed difference in means (-0.004) can be due to random chance rather than a true underlying difference.

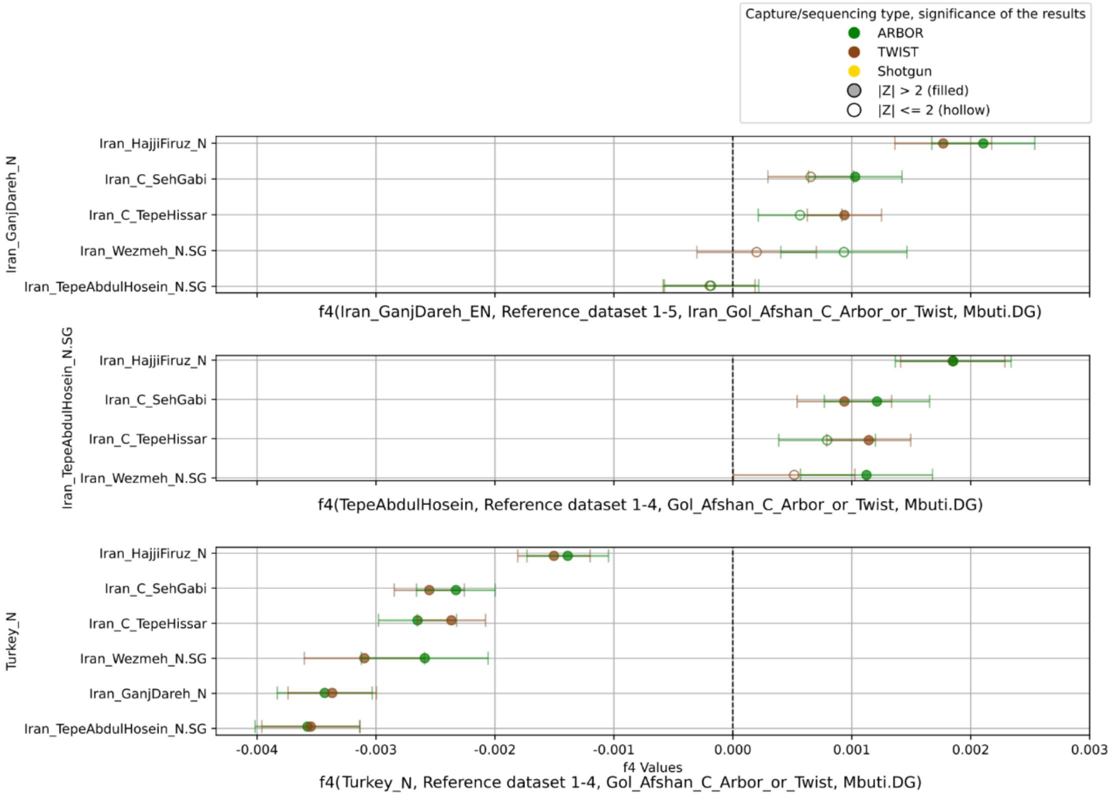


**Figure 11.5. f4-statistics in various forms. They show shared drift of the Early Copper Age Gol Afshan Tepe individual with different Neolithic and Copper Age reference sample sets (Supplementary Table S11).** f4 statistics are calculated A) in the form of f4(Iran_GanjDareh_EN, Reference_dataset 1-5, Iran_Gol_Afshan_C_Arbor_or_Twist, Mbuti.DG), B) f4(TepeAbdulHosein, Reference dataset 1-4, Gol_Afshan_C_Arbor_or_Twist, Mbuti.DG), and C) f4(Turkey N, Reference dataset 1-4, Gol_Afshan_C_Arbor_or_Twist, Mbuti.DG).

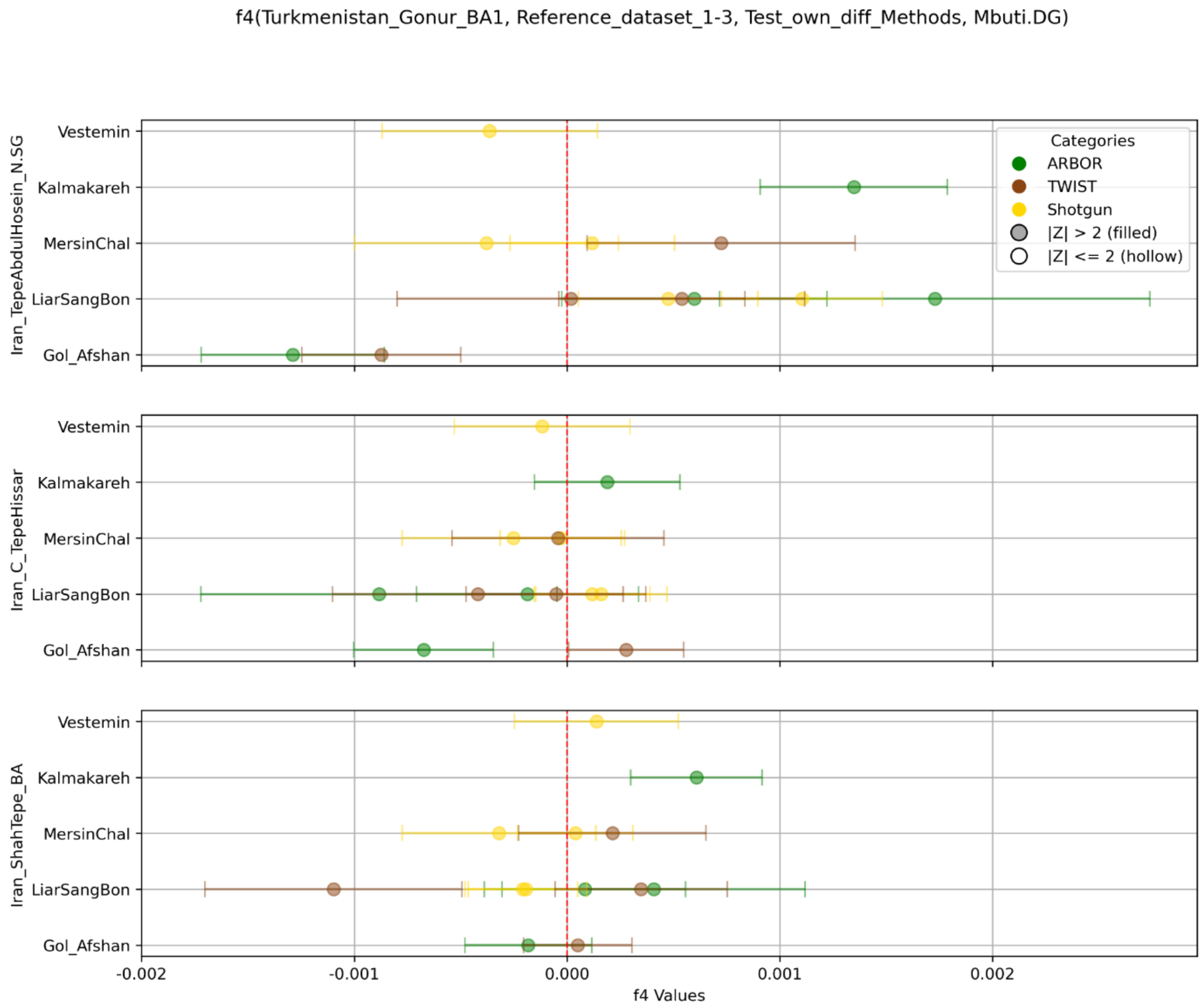


**Figure 11.6. f4-statistics in the form of f4(Turkemnistan_Gonur_BA1, Reference dataset 1-3, Test_own_diffMethods, Mbuti.DG).** Gol Afshan Tepe shares a significant excess of drift with N Tepe Abdul Hosein, compared to Gonur_BA1. Kalmakareh and Lirasangbon on the other hand have significant excess of Gonur_BA1 type allele frequencies, compared to the tested Iranian C-N datasets. Shah Tepe and Gonur_BA1 have so similar genomic compositions, that none of the analysed samples shows significant allele-sharing distributions between the two (Supplementary Table S11).

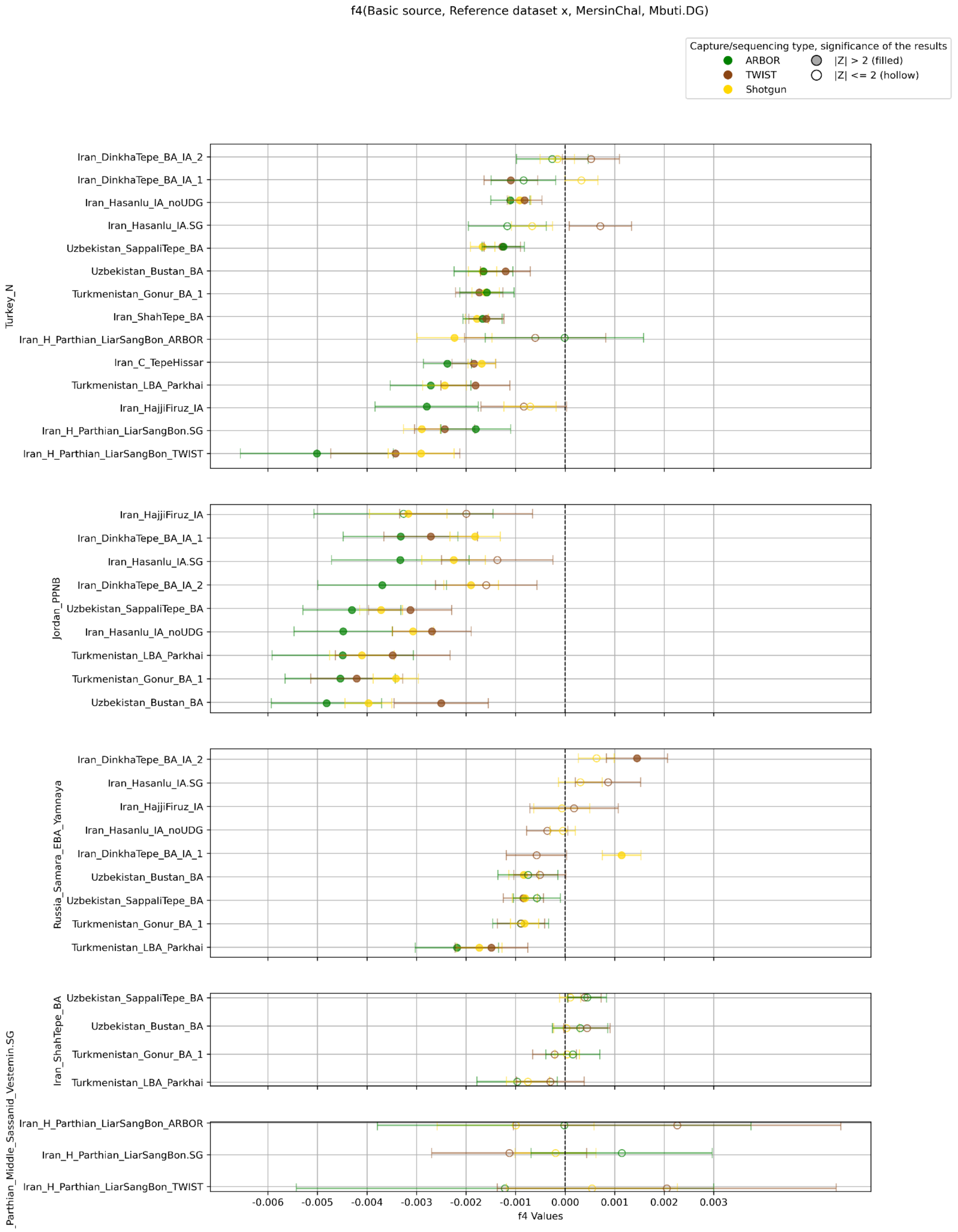


**Figure 11.7. f4-statistics with differently processed Marsin Chal samples from the Achaemenid-Seleucid period.** The tests were run in the following form: $f_4$(Basic source, Reference dataset x, Marsin Chal, Mbuti.DG). The $f_4$ tests with Shah Tepe and Vestemin are non-significant (Z<2) for the Marsin Chal samples (Supplementary Table S11). Jordan PPNB as a potential basic source separates the affinities of Arbor and shotgun/Twist results, where the former had the least data in the tests.

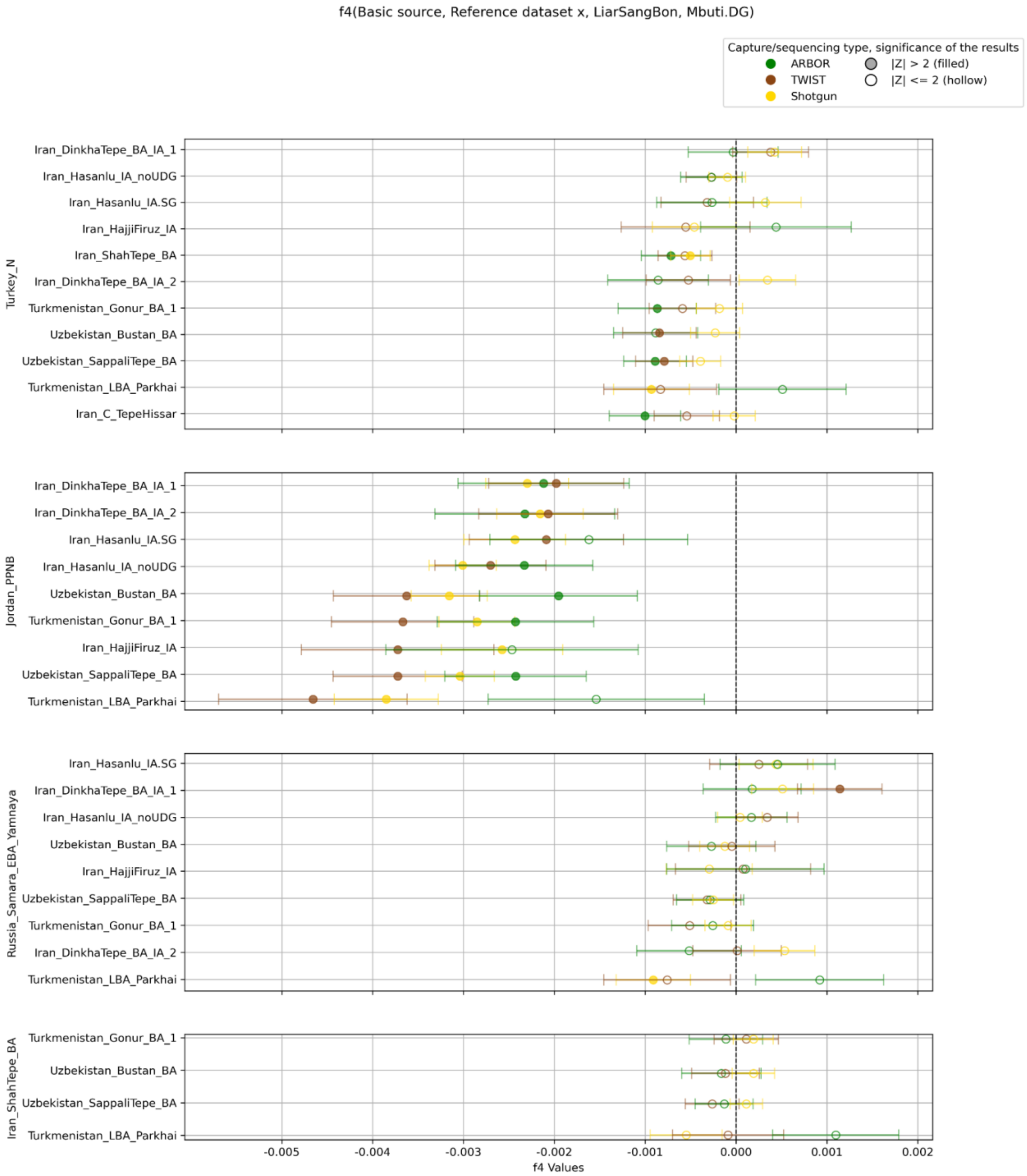


**Figure 11.8. f4-statistics with differently processed Liarsangbon samples from the Parthian period.** The tests were run in the following general form: f4(Basic source 1-4, Reference dataset x, Liarsangbon, Mbuti.DG). The $f_4$ tests with Russia_Samara_EBA_Yamnaya and Shah Tepe as basic sources are non-significant (Z<2) for the Liarsangbon samples (Supplementary Table S11), due to either the low level of allele sharing between the steppe people and the tested groups, or the general homogeneity in the BA north Iran and the contemporary Turan. Arbor samples tend to have less difference in allele sharing with Jordan PPNB and the given reference groups than the Twist and shotgun samples, although this type of capture results had also the lowest number of SNPs covered. The largest difference in f4 was observed with Parkai LBA in the test, which is a single 1240k genome with 442k SNPs covered.

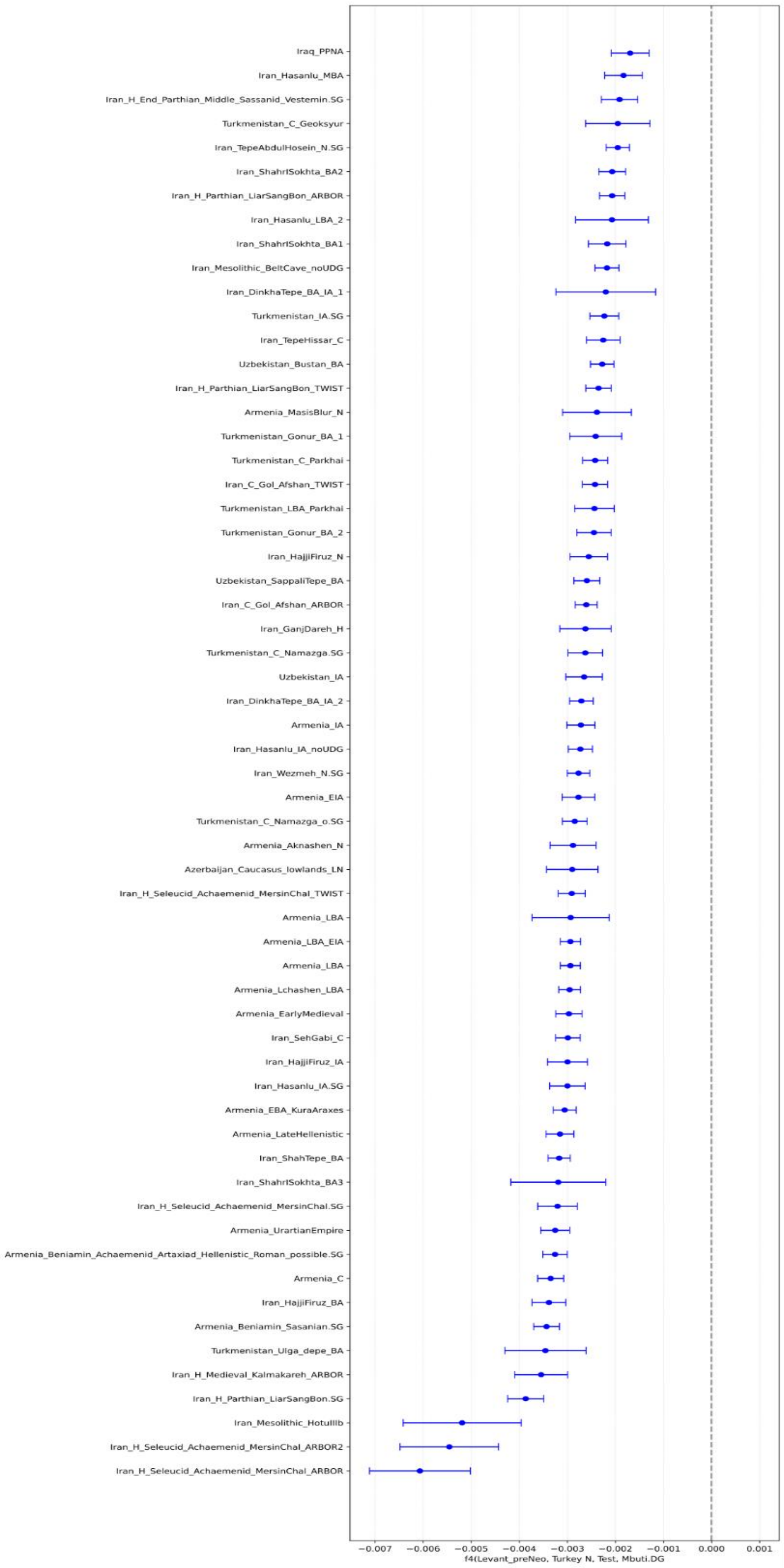


**Figure 11.9**. **Errorbar of f4-statistics in the form of (Levant pre-Chalcolithic, ANF/TurkeyN, Test, Mbuti).** The test itself is seen in Table S11.
